# Neural and Experiential Dynamics of High-Ventilation Breathwork Mirror Psychedelic States

**DOI:** 10.64898/2026.11.11.765965

**Authors:** Lana K. Porter, Ece Kaya, Kamlesh Chalise, Janina Klarmann, Armin Goffin, Muad Abd El Hay, Nina Hartter, Borja Rodriguez-de-Santiago, Joshua N. Hindmarsh, Naman Jain, Abdellatif Nemri, Barbara Breitenstein, Christopher Timmermann, Marieke L Schölvinck, Isabel Dziobek, Martha N. Havenith

**Affiliations:** Zero-Noise Lab, Ernst Strüngmann Institute of the Max Planck Society, Frankfurt a.M., Germany; Department of Neuropsychology and Psychopharmacology, Faculty of Psychology and Neuroscience, Maastricht University, Maastricht, Netherlands; Department of Psychotherapy, Institute for Psychology, Humboldt University, Berlin, Germany; Department of Clinical Psychology of Social Interactions, Institute for Psychology, Humboldt University, Berlin, Germany; Into The Deep Breathwork Coaching, Berlin, Germany; Department of Experimental Psychology, Faculty of Brain Science, University College London, London, UK

**Keywords:** high-ventilation breathwork, altered states of consciousness, EEG, complexity

## Abstract

High-ventilation breathwork (HVB) is a simple breathing technique that nevertheless frequently triggers experiences resembling psychedelic states. While recent studies have begun to pinpoint physiological and neural changes involved in altered states of consciousness (ASCs) evoked by HVB, the interplay between neural and experiential dynamics throughout an HVB session remains unclear. Here we tracked neural activity (via 64-channel EEG recordings) as well as subjective experience depth and content in 25 experienced HVB practitioners across resting-state recordings interleaved throughout a naturalistic HVB session. We find that neural dynamics evoked by HVB emulate the most robust neural signatures of classical psychedelic states, specifically (1) loosening of temporal structure, including broadband cortical desynchronization, increased signal complexity, and decreased oscillatory power in the Alpha (8 - 13 Hz) range, and (2) reorganization of spatial structure, including a flattening of cortical processing hierarchy. These neural effects directly correlated with participants’ subjective depth with minute resolution, but were not explained by isolated features of HVB such as music listening or active interoception. Together, these results identify unifying neural features of psychedelic-like ASCs regardless of induction method, demonstrating that cortical states triggered by psychedelics can also be accessed by non-pharmacological routes, and breathwork is a powerful way to do so.

## Introduction

High-ventilation breathwork (HVB) is a collection of breathing practices that feature deep and somewhat accelerated breathing, often accompanied by guided facilitation and evocative music. Prominent versions of HVB include Rebirthing, Holotropic and Conscious-Connected Breathwork (de-Wit et al., 2018; Grof & Grof, 2010; Havenith, M & Nemri, A, 2025). Unlike slow-paced breathing techniques, HVB regularly triggers highly immersive altered states of consciousness (ASCs) that resemble psychedelic states (Eyerman, 2013; Fincham et al., 2026; Havenith et al., 2025). As such, typical effects of HVB can include intense emotional breakthroughs, perceived dissolution of time, space or self, as well as sensory modifications and hallucinations, with particular emphasis on interoception-based experiences (Bahi et al., 2024; Canello et al., 2026; Fincham et al., 2026b; Havenith et al., 2025a; Havenith, M & Nemri, A, 2025; Kartar et al., 2025; Lewis-Healey, 2024).

These experiential parallels between HVB and psychedelics raise the question how the brain can generate such drastic departures from ordinary waking consciousness without pharmacological help. Recent studies have begun to trace the physiological mechanisms that play a crucial role in triggering HVB experiences, highlighting a physiological cascade in which intense breathing depletes CO2 levels (see Kety & Schmidt, 1948), thereby raising blood pH and restricting cortical blood flow (CBF) and tissue oxygenation (see Bednarczyk et al., 1990). These physiological changes correlate directly with the depth of HVB experiences, outlining their central role in supporting these subjective shifts (Havenith et al., 2025; Kartar et al., 2025; Schmitz et al., 2026). In contrast, the corresponding neural dynamics underpinning HVB experiences are only beginning to emerge.

As we begin to characterize the neural signatures of HVB, psychedelic research provides a useful benchmark, pinpointing promising candidate mechanisms. Neural responses to serotonergic psychedelics, from DMT to LSD and psilocybin, have been proposed to share several neural features, including (1) loosening of temporal structure, including broadband cortical desynchronization (Godfrey et al., 2025; Muthukumaraswamy et al., 2013; Pallavicini et al., 2019; Subramani et al., 2026; Timmermann et al., 2019), increased complexity (i.e. lack of predictability) of neural signals (Carhart-Harris et al., 2016; Carhart-Harris, 2026; Kroupi et al., 2026; Lebedev et al., 2016; Lewis-Healey, 2024b; Lyons et al., 2026; Schartner et al., 2017), and decreased oscillatory power in the Alpha range (8 - 13 Hz; Carhart-Harris et al., 2016; Godfrey et al., 2025; Muthukumaraswamy et al., 2013; Timmermann et al., 2019), and (2) a reorganization of spatial structure, including a flattening of the cortical processing hierarchy, and disintegration of large-scale networks (Alamia et al., 2020; Carhart-Harris et al., 2012, 2014; Carhart-Harris & Friston, 2019; Gattuso et al., 2023; Girn et al., 2022, 2026; Pines et al., 2026; Preller et al., 2018; Tagliazucchi et al., 2014).

Each of these features has been associated with core aspects of subjective psychedelic phenomenology, including ego dissolution, visual hallucinations as well as experiential and emotional intensity (Carhart-Harris et al., 2016; Kroupi et al., 2026; Lewis-Healey, 2024; Madsen et al., 2021; Preller et al., 2018; Schartner et al., 2017; Subramani et al., 2026; Tagliazucchi et al., 2016; Van Elk et al., 2019). Thus, given the experiential similarity between psychedelics and HVB, it is conceivable that HVB experiences are also underpinned by similar neural features. This hypothesis has only recently begun to be explored. In 2024, Bahi et al. recorded resting-state EEG following an HVB session, reporting post-session decreases of oscillatory power in Delta (1-4 Hz), Theta (5-8 Hz) and Beta (14-30 Hz) frequencies, together with increased Gamma power (30 - 45 Hz) observed only in experienced practitioners. Also in 2024, Lewis-Healey et al. traced neural signatures within an HVB session, analysing portable 8-channel EEG recordings from 14 participants practicing SOMA breathwork, a version of HVB that includes regular breath holds. Their study identified increased EEG signal complexity (Lempel & Ziv, 1976) as a hallmark of breathwork-evoked ASCs, a promising parallel to psychedelic states (Lewis-Healey, 2024; Schartner et al., 2017).

In the present study, we build on these findings by collecting high-density (64-channel) EEG recordings together with time-resolved experience reports throughout a naturalistic HVB session, compared to two control conditions that mimic central aspects of HVB (Progressive Muscle Relaxation (PMR) to account for active interoception and guided facilitation, and passive music listening). This enables us to directly trace the neurophenomenological dynamics of HVB, with a focus on testing the relation between real-time ASC intensity and neural signatures that have been reported for psychedelic states, including broadband cortical desynchronization, decreased Alpha power, increased signal complexity, and a reshuffling of the hierarchical organization of cortical activity. With this approach, we aim to delineate shared and unique features of psychedelic-like ASCs evoked by pharmacological and non-pharmacological routes.

## Results

The study includes 25 participants (18 female, 7 male, aged 43.60 ± 8.16 (all values given as mean ± standard deviation unless stated otherwise; see Supp. Table S1 for full demographics), who had all completed a minimum of five HVB sessions prior to their study participation. Following basic participant screening, pre-session surveys were administered online one week before the in-person HVB session (Fig. 1A). Specifically, we surveyed aspects of affect and well-being via the Warwick-Edinburgh Mental Well-being Scale (WEMWBS; Tennant et al., 2007), State-Trait Anxiety Inventory (STAI: Spielberger, 1983), Positive and Negative Affect Schedule (PANAS; Watson et al., 1988), and Watts Connectedness Scale (WCS; Watts et al., 2022).

**Figure 1.**
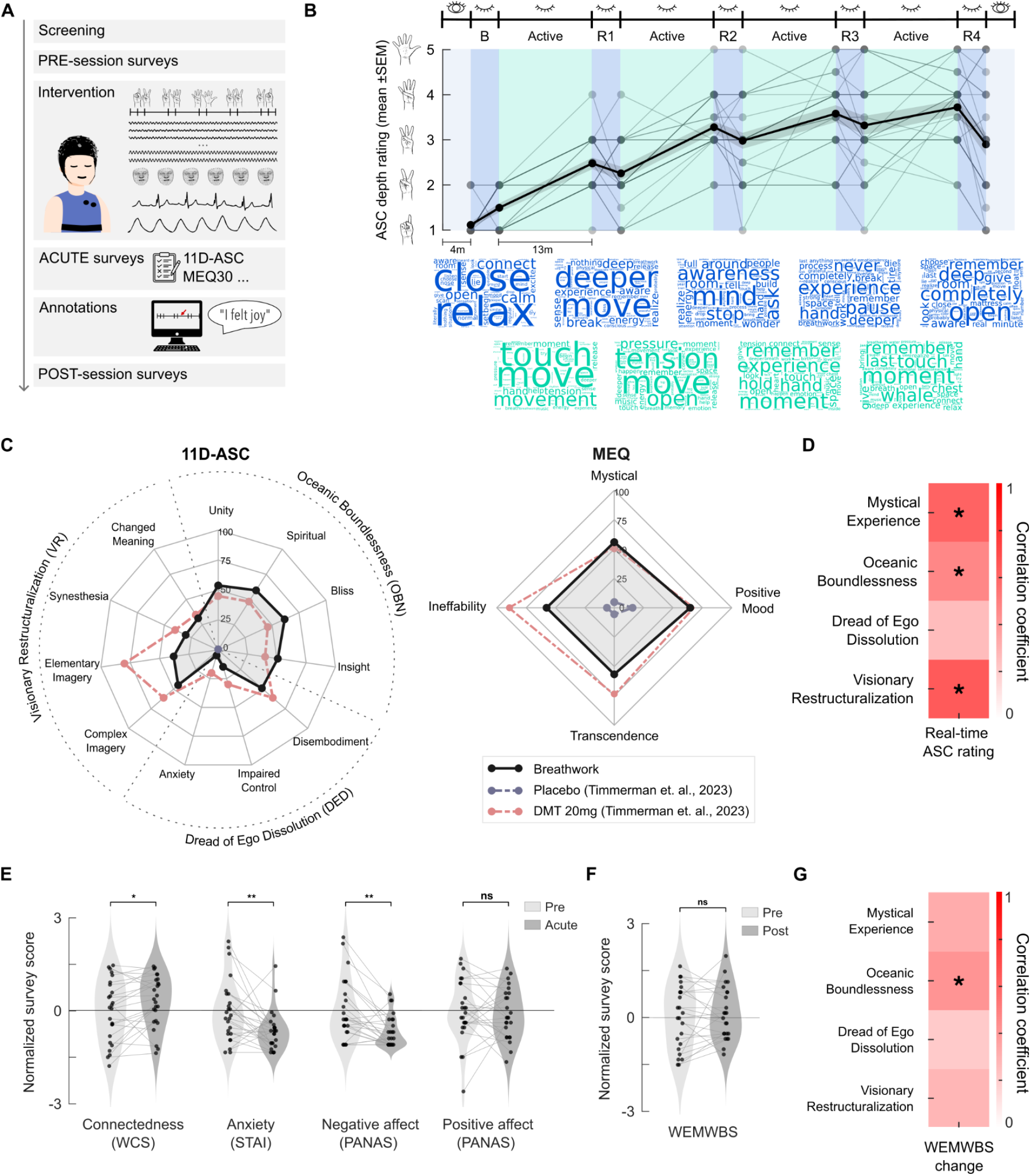
Phenomenological effects of HVB. **A)** Graphical summary of experimental design. **B)** Top: Graphic illustration of session epochs, including eyes-open and eyes-closed resting-state epochs (blue, 4 minutes duration), and active-breathing epochs (turquoise, 13 minutes duration). Black line: Average real-time ASC intensity rating. Shading: SEM. Grey lines: Raw ratings of individual participants, reported by hand signs (see y axis illustration). Bottom: Word clouds from post-session video annotations corresponding to each timepoint. Blue: Annotations for each resting-state epoch. Turquoise: Same for active-breathing epochs. **C)** Average 11D-ASC (left) and MEQ30 (right) scores (n = 25 participants). Black lines: Average ratings for breathwork. Red lines: Ratings for ASCs triggered by intravenous DMT (from Timmermann et al., 2023). Grey lines: Corresponding placebo scores from the same study. **D)** Correlation coefficients between real-time ASC ratings and retrospective survey scores (MEQ30 total score, OBN, VR and DED). Color represents correlation coefficient (Pearson’s r); * p <0.05 after BH-FDR correction. **E)** Acute changes in connectedness and affect. Violin plots: Z-scores of survey scores, normalized by average pre-session baseline scores, for WCS, STAI PANAS-positive and PANAS-negative. Light grey: One week pre-session, i.e. baseline. Dark grey: Acute post-session. Connected dots: Individual participants (n = 25). Stars: Statistical significance based on paired-samples t-tests (* p <0.05, ** p <0.01). **F)** Same as E for WEMWBS scores collected one week pre- and post-session (n = 25). **G)** Same as D for the correlation coefficients between 11D-ASC and MEQ30 scores, and WEMWBS changes at one-week follow-up (uncorrected).

The HVB session was designed based on typical formats of Conscious-Connected Breathwork, with participants lying down on a mattress with their eyes closed, and practicing deep, somewhat accelerated breathing in the presence of a trained breathwork facilitator. During the active-breathing portions of the session, experiences were accompanied by music, and facilitators were free to support participants verbally and through supportive touch (‘bodywork’), as agreed prior to the session (see Intervention Protocol and Touch Protocol provided in Supp. Texts S1-S2). For the purpose of EEG recording, these active-breathing phases were interleaved with four short resting-state epochs throughout the session, during which music and active facilitation were discontinued, and participants were asked to breathe normally. To enable time-resolved tracking of experience intensity throughout the session, participants gave real-time ratings of altered-state-of-consciousness (ASC) intensity at the beginning and end of each resting-state epoch, i.e. across a total of 10 timepoints throughout the session (Fig. 1B). To minimize the disruption to breathwork experiences, these ratings were given non-verbally, via hand signs, on a 5-point scale (1 = normal waking consciousness; 5 = near-complete loss of time/space perception, strongly reduced responsiveness, and unrestrained emotional or bodily expression): Anchoring examples of scoring criteria were given to participants before the session (see Methods for scoring criteria).

As such, one session lasted a total of 80 minutes, comprising five four-minute eyes-closed resting state epochs (Baseline (B) and Resting State (R1 - R4; see Fig. 1B), interleaved with four thirteen-minute phases of active breathing epochs (Active (BR1 - BR4). These elements were book-ended by two four-minute eyes-open resting state recordings, which were not analyzed further for the purpose of this study (Fig. 1B). Throughout the session, we recorded 64-channel EEG (BrainCap-Fast’n Easy 64Ch) together with bipolar ECG (BrainAmp ExG system) to derive a cardiac-derived respiration signal, as well as video footage of participants’ faces for the post-session annotation of experiential content (see below; Fig. 1A). All EEG analyses were based on the five recorded eyes-closed resting-state epochs, with the first epoch (preceding the first active-breathing epoch) serving as baseline.

After a brief post-session break, participants were first asked to complete standard surveys about their experience, specifically the 11-Dimensions of Altered States of Consciousness (11D-ASC; Dittrich, 1998; Studerus et al., 2010) and Mystical Experience Questionnaire (MEQ30; Barrett et al., 2015). We also collected acute post-session scores of affect, including the STAI, PANAS and WCS (see above). Note that WEMWBS scores were not collected acutely post-session, because the WEMWBS is not designed to track short-term shifts in well-being (Tennant et al., 2007). Finally, participants were asked to review the video footage of their session in the presence of experimenter and facilitator, with the opportunity to verbally annotate descriptions of their subjective experience at any point in the recording (see Supp. Text S3 for example annotations). In this way, video and audio footage served as anchors to prompt and temporally organize remembered experiences, an important scaffold given that time perception is often drastically disrupted during breathwork experiences (Havenith et al., 2025; Lewis-Healey, 2024; Sviderskaya & Bykov, 2006). One week post-session, participants again completed the online surveys collected pre-session, i.e. the WEMWBS, STAI, PANAS and WCS.

### HVB evokes altered states of consciousness

To characterize subjective experiences evoked by breathwork, we collected 10 real-time ratings of ASC intensity throughout the session (see above and Methods) which enabled us to trace experiential dynamics in a time-resolved way. Throughout the session, participants reported a gradual increase in ASC intensity from 1.12 (± 0.33) at baseline (B) to 3.72 (± 0.82) at the end of the final active-breathing epoch (B - R4; Wilcoxon signed-rank; n = 25; r = 1.0; d = 3.18; p < 0.001), with an overall average ASC rating of 3.09 (± 0.62) across within session timepoints (R1 - R4; Fig. 1B). This temporal progression of experiences was also reflected in our post-session phenomenological annotations: word clustering of all verbal annotations linked to different epochs of the HVB session hints at a transition of experiential qualities from frequently used descriptors like ‘close’ and ‘relax’ at the beginning of session to words like ‘tension’ and ‘move’ in the active phase of the HVB session, arriving at an increasingly diverse collection of descriptions including phrases like ‘remembering’ and ‘completely open’ towards the end of the session (Fig. 1B). Interestingly, ASC ratings dropped slightly but consistently by the end of each resting-state epoch, during which participants returned to ordinary breathing (epoch start: 3.11 ± 0.64; epoch end: 2.85 ± 0.77; mean difference: 0.26 ± 0.59; paired t-test; n = 25, t(24) = 2.20, d = 0.44, p = 0.038). The small magnitude of this consistent drop suggests that while active breathing is instrumental in maintaining HVB states, subjective experiences were largely able to self-sustain throughout the four-minute resting-state epochs in which active breathing, music and active facilitation were interrupted.

In addition to real-time ratings, the two standard phenomenological surveys of ASC experiences we collected acutely post-session, the 11D-ASC and MEQ30 (Fig. 1C), reveal that HVB evokes strong ASCs, approximating some of the typical effects of classical psychedelics, particularly the positively-valenced ego-dissoluting dimensions, with reference scores for dimethyltryptamine (DMT) shown as an example (Fig. 1C; DMT reference data reproduced from Timmermann et al., 2023). For HVB, the three subscales of the 11D-ASC reached average scores of 56.4 (± 23.7) for Oceanic Boundlessness (OBN), 36.1 (± 22.2) for Visionary Restructuralization (VR), and 22.8 (± 11.1) for Dread of Ego Dissolution (DED: see Supp. Table S2 for all scores). OBN scores were statistically indistinguishable between HVB and DMT (one-sample t-tests against DMT reference, n = 25; DMT reference = 44.75; t(24) = 2.46, d = 0.49, p = 0.063; significance Šidák-corrected across all three comparisons; Šidák, 1967). In contrast, VR and DED both reached lower scores for HVB than DMT (VR: DMT reference = 54.0; t(24) = 4.03, d = 0.81, p = 0.0015; DED: DMT reference = 37.0; t(24) = 6.40, d = 1.28, p < 0.001; Šidák-corrected). Similarly, the four subscales of the MEQ30 reached average scores of 56.7 (± 24.7) for Transcendence, 64.7 (± 17.1) for Positive Mood, 57.6 (± 25.5) for Ineffability, and 55.8 (± 22.8) for Mystical Experience (see Supp. Table S2 for all scores). Compared to DMT reference scores, Transcendence, Positive Mood and Mystical Experience did not differ from DMT (one-sample t-tests against DMT reference; n = 25; Transcendence: DMT reference = 70; t(24) = 2.69, d = 0.54, p = 0.050; Positive Mood: DMT reference = 60; t(24) = 1.37, d = 0.27, p = 0.55; Mystical Experience: DMT reference = 49; t(24) = 1.49, d = 0.30, p = 0.48; significance Šidák-corrected across four subscales). Ineffability scored significantly lower for HVB than DMT (DMT reference = 85; t(24) = 5.37, d = −1.07, p < 0.001; Šidák-corrected). As a result, nine HVB participants (36%) met the criteria for a complete mystical experience as set out by Barrett et al. (2015), a proportion that is lower than the 60% reported for high-dose psilocybin (Garcia-Romeu et al., 2014), but higher than the 12.5% reported for moderate-dose LSD (Liechti et al., 2017).

Importantly, average real-time ratings of ASC depth correlated well with retrospective survey scores (Fig. 1D). Average real-time ASC ratings significantly correlated with Total Mysticical Experience (MEQ Total) scores as well as with the OBN and VR (Pearson’s correlation; n = 25; MEQ Total: r = 0.61, p = 0.001; OBN: r = 0.46, p = 0.022; VR: r = 0.65, p < 0.001; corrected for multiple comparisons across four scales using Benjamini Hochberg-False Discovery Rate (BH-FDR) correction; Benjamini & Hochberg, 1995). The exception was DED, which was not significantly linked to real-time ASC intensity ratings (Pearson’s r = 0.31, p = 0.126; see Supp. Fig. S1). These relations suggest that real-time ASC ratings are a reliable metric to track ASC depth throughout the session, and are more closely linked to positive than aversive elements of ASCs.

### HVB acutely alters affect

Acutely post-session, participants reported a significant decrease in state anxiety (STAI) and negative affect (PANAS-negative), as well as a significant increase in connectedness (WCS) compared to one week pre-session (Fig. 1E; paired t-test, n = 25; STAI: mean score 33.48 ± 10.05 to 26.88 ±6.46; t(24) = −2.98, p = 0.006; PANAS negative: mean score 15.36 ±4.91 to 12.12 ±2.51; t(24) = −3.69, p = 0.001; WCS: mean score 74.93 ±15.32 to 79.73 ±12.79; t(24) = 2.27, p = 0.033; all values significant after BH-FDR correction). In contrast, positive affect (PANAS-positive), remained stable compared to baseline (Fig. 1E; see Supp. Table S3 for complete statistics).

The observed changes largely returned to baseline one week post-session (Supp. Fig. S2; Supp. Table S3). This lack of significant long-term impact also applied to participants’ WEMWBS ratings, which were measured one week post-session. WEMWBS scores did not significantly change from the pre-session baseline to the post-session follow up (Fig. 1F; paired t-test; n = 25; mean score 51.64 ± 6.39 vs. 52.12 ± 6.21, t(24) = 0.45, p = 0.654). However, exploratory analysis revealed that participants who experienced stronger OBN during the session showed improved well-being one-week post session (Fig. 1G; Pearson’s correlation; n = 25; r = 0.42, p = 0.037; uncorrected), an effect that is consistent with previous reports both for breathwork and psychedelic experiences (Havenith et al., 2025; Ko et al., 2022).

Together, these analyses suggest that HVB consistently evoked deep ASC experiences, which to a large extent resembled ASCs evoked by classical psychedelics such as DMT (see also Bahi et al., 2024; Havenith et al., 2025; Kartar et al., 2025; Lewis-Healey, et al., 2024). In terms of temporal dynamics, ASC depth typically reached a peak towards the second half of the HVB session and continued largely undisrupted throughout each resting-state epoch.

### HVB dynamically reorganizes EEG spectral power

To characterize the neural dynamics associated with HVB experiences, we first analyzed EEG spectral power across the session’s five resting-state epochs (baseline and R1-4) by dissecting total spectral power across the frequency range of 1-45 Hz into its aperiodic and periodic components (Fig. 2A). To quantify changes in spectral power, two complementary levels of analysis were applied: (1) a participant-level modulation index (Fig. 2B; see Methods), summarizing each individual’s normalized change from baseline across time points R1-4, and (2) a sensor-level cluster-based permutation test (Fig. 2C; see Methods), which localizes reliable changes across the scalp at the population level.

**Figure 2.**
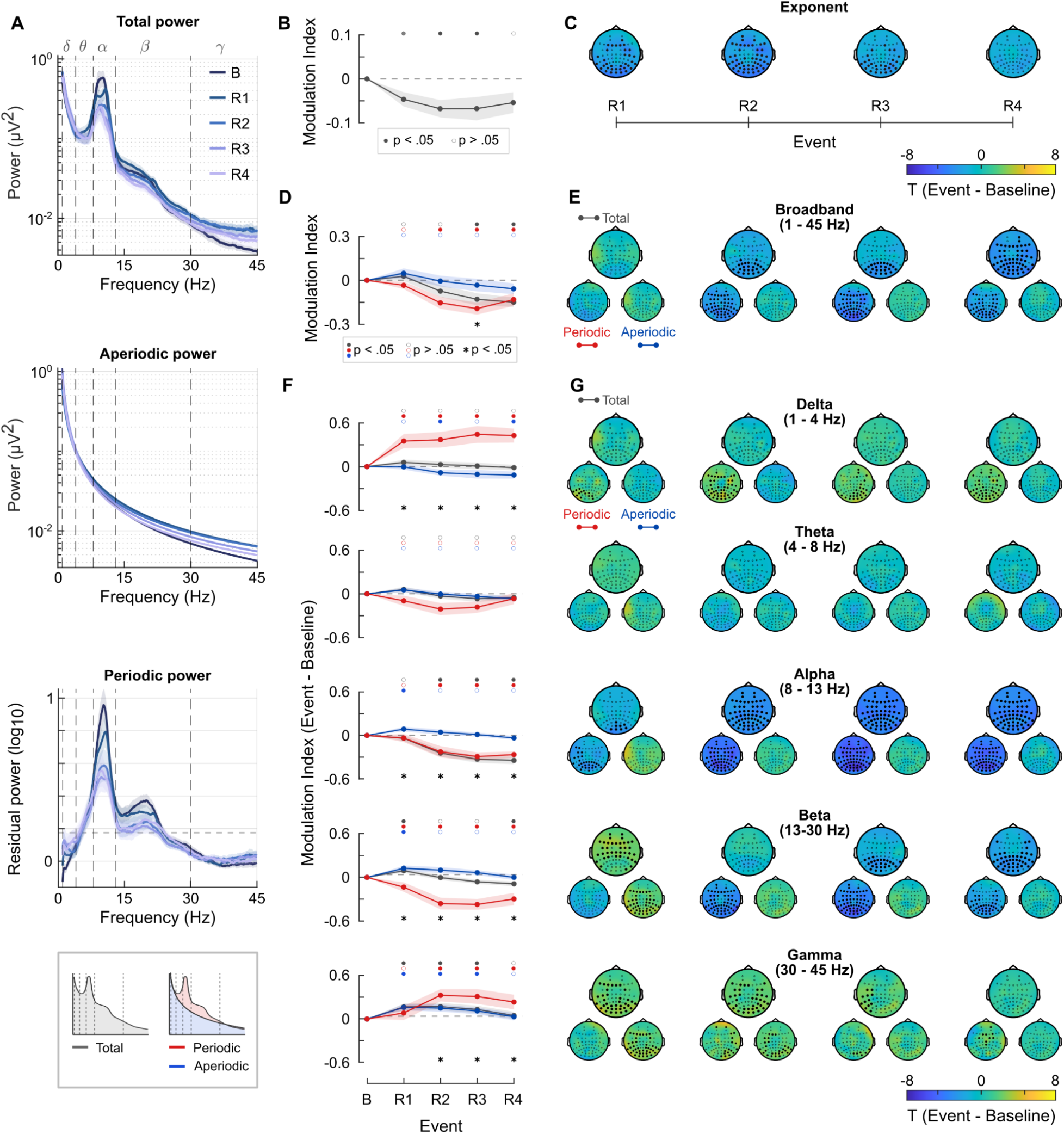
EEG spectral power throughout HVB. **A)** Density of total spectral power (top), aperiodic power (center), and periodic power (bottom) across frequencies of 1-45 Hz. Colored lines: Power spectra across five resting-state epochs (see in-figure legend). Dashed lines delineate frequency bands from Delta to Gamma (see labels above top panel). **B)** Modulation index relative to baseline for aperiodic exponent across R1 - R4. Shaded region: SEM. Filled circles above the panel indicate significance according to signed-rank Wilcoxon tests for deviation from zero (filled symbol: p <0.05; BH-FDR). **C)** Topological sensor maps of aperiodic exponent at each timepoint (R1-R4) relative to baseline. Color: t-value of deviation from baseline (see in-figure color bar). Black markers: electrode locations that form part of significant spatial clusters after two-sided cluster-permutation inference and BH-FDR correction. Markers indicate cluster membership. **D)** Same as B for total (grey), periodic (red), and aperiodic (blue) broadband power. Black asterisks below panel indicate a significant difference between modulation indices for periodic and aperiodic power (also tested by signed-rank Wilcoxon test). **E)** Same as C for broadband spectral power, segregated by total power (top), periodic power (left), and aperiodic (right) power. **F)** Same as D for each canonical frequency band (see labels in G). **G)** Same as E for canonical frequency bands (see in-figure labels).

In terms of overall spectral power, the aperiodic exponent of power spectra (Fig. 2A, center) decreased significantly across time points R1-R3 compared to baseline (Fig. 2B; all analyses use Wilcoxon signed-rank modulation indices, matched-pairs rank-biserial; n = 25; R1: r = −0.58, p = 0.012; R2: r = −0.69, p = 0.0025; R3: r = −0.55, p = 0.017; significance corrected across four epochs using BH-FDR correction across epochs unless otherwise stated; see Supp. Table S4 for full statistics). This effect was most prominent in posterior electrodes, as well as some frontal electrodes (Fig. 2C; permutation-based cluster test; n = 25; 5,000 permutations; R1: Σt = 103.7, p = 0.008; R2: Σt = 158.4, p = 0.003; R3: Σt = 68.6, p = 0.025; BH-FDR; Supp. Table S4). This decrease in the spectral 1/f slope does not necessarily suggest a change in total power, but rather a shift in the balance between low-frequency and high-frequency components, with high-frequency activity becoming relatively more pronounced (see Fig. 2A, center) throughout a large part of the HVB session.

Complementary to this, total broadband power also decreased relative to baseline, particularly in the second half of the HVB session (Fig. 2D; Wilcoxon signed-rank; R3: r = −0.61, p = 0.0074; R4: r = −0.78, p < 0.001; BH-FDR; Supp. Table S4). This decrease in spectral power was first localized to posterior electrodes at R2, before extending across nearly the whole scalp by R4 (Fig. 2E; cluster permutation; R2: Σt = 63.5, p = 0.025; R3: Σt = 70.9, p = 0.025; R4: Σt = 180.1, p < 0.001; BH-FDR; Supp. Table S4). This reduction appeared to be largely driven by drastic decreases in periodic broadband power from R2 onward (Fig. 2D; Wilcoxon; R2: r = −0.70, p = 0.002; R3: r = −0.88, p < 0.001; R4: r = −0.60, p = 0.009; BH-FDR), which was distributed across the entire scalp (Fig. 2E; cluster permutation; R2: Σt = 166.9, p = 0.002; R3: Σt = 215.1, p < 0.001; R4: Σt = 121.0, p = 0.008; BH-FDR). In contrast, aperiodic broadband power showed no significant deviation from baseline at any timepoint (Supp. Table S4). As a result, changes in broadband periodic power were generally larger than in aperiodic power, a difference that reached statistical significance for timepoint R3 (Wilcoxon signed-rank test on paired difference in modulation index; n = 25; r = −0.58, p = 0.012, BH-FDR). These results indicate that the decrease in broadband spectral power was driven specifically by decreased neural oscillations rather than simply a shift in the slope or offset of aperiodic spectral power.

To explore how these dynamics mapped onto different frequencies, we computed total, periodic and aperiodic power for five canonical frequency bands from Delta to Gamma (Figs. 2F-G). These analyses showed a highly frequency-specific impact of HVB on neural activity. For instance, oscillatory power in the Delta band (1-4 Hz) increased consistently from baseline across the entire session (Fig. 2F; Wilcoxon; R1: r = 0.74, p = 0.0012; R2: r = 0.71, p = 0.0019; R3: r = 0.71, p = 0.0019; R4: r = 0.79, p < 0.001; BH-FDR) with changes localized mostly towards posterior electrode locations (Fig. 2G; cluster permutation; R1: Σt = 35.8, p = 0.007; R2: Σt = 128.6, p < 0.001; R3: Σt = 67.3, p < 0.001; R4: Σt = 98.4, p < 0.001: BH-FDR). In contrast, aperiodic Delta power did not increase, and in fact showed a significant decrease at time points R2 and R4 (Wilcoxon; R2: r = −0.52, p = 0.023; R4: r = −0.52, p = 0.023; BH-FDR), though this decrease did not translate to a significant sensor cluster at either time point (Supp. Table S4). As such, changes in periodic Delta power were significantly larger than in aperiodic Delta power across the session (Fig. 2F; Wilcoxon signed-rank test on the paired difference in modulation index; n = 25; R1: z = 3.08, r = 0.71, p = 0.002; R2: z = 3.24, r = 0.74, p = 0.001; R3: z = 3.16, r = 0.72, p = 0.002; R4: z = 3.73, r = 0.85, p < 0.001; BH-FDR). In contrast, activity in the Theta range (4 - 8 Hz) did not significantly change from baseline at any time point (Supp. Table S4).

For the Alpha band (8 - 13 Hz), periodic and total power significantly decreased from baseline, an effect that emerged earlier at the sensor level than in the participant-level modulation: cluster-level decreases were significant from R1 onwards, spreading from posterior electrode locations towards the entire scalp (Fig. 2G; cluster permutation; periodic: R1: Σt = 77.4, p = 0.019; R2: Σt = 307.3, p < 0.001; R3: Σt = 354.4, p < 0.001; R4: Σt = 313.2, p < 0.001; total: R1: Σt = 21.7, p = 0.036; R2: Σt = 195.6, p < 0.001; R3: Σt = 244.9, p < 0.001; R4: Σt = 232.2, p < 0.001; BH-FDR), whereas participant-level modulation indices reached significance from R2 onwards (Fig. 2F; Wilcoxon; periodic: R2: r = −0.78; R3: r = −0.95; R4: r = −0.95; total: R2: r = −0.82; R3: r = −0.97; R4: r = −0.91; all p < 0.001; BH-FDR). Contrary to this global periodic decrease, aperiodic Alpha power remained largely unchanged, with a small but significant participant-level increase at R1 (Fig. 2F; Wilcoxon; r = 0.59, p = 0.009; BH-FDR) that did not translate to a significant sensor cluster (Fig. 2G; Supp. Table S4). Changes in periodic Alpha power also differed significantly from those in aperiodic Alpha power at every time point (Fig. 2F; Wilcoxon; R1: z = -2.27, r = −0.52, p = 0.023; R2: z = −3.05, r = −0.70, p = 0.002; R3: z = −3.97, r = −0.91, p < 0.001; R4: z = −3.38, r = −0.77, p < 0.001; BH-FDR).

Spectral power in the Beta band (13-30 Hz) showed the most diverse temporal sequencing: aperiodic and total power initially increased significantly at R1 (Fig 2F; Wilcoxon; aperiodic: R1: r = 0.71, p = 0.002; total: R1: r = 0.63, p = 0.006; BH-FDR), an effect that was globally visible across electrode locations (Fig. 2G; cluster permutation; aperiodic: R1: Σt = 124.8, p = 0.002; total: R1: Σt = 104.7, p = 0.010; BH-FDR). As the session progressed, total and aperiodic Beta power then declined again, with total Beta power reaching significant decreases at the participant level at R4 (Fig. 2F; Wilcoxon; R4: r = −0.58, p = 0.012; BH-FDR), largely in posterior regions (Fig. 2G; cluster permutation; R3: Σt = 65.5, p = 0.021; R4: Σt = 122.2, p = 0.006; BH-FDR; Supp. Table S4). At the same time, periodic Beta power decreased consistently from R1 through R4 at the participant level (Fig. 2F; Wilcoxon; R1: r = −0.48, p = 0.035; R2: r = −0.81, p < 0.001; R3: r = −0.88, p < 0.001; R4: r = −0.65, p = 0.005; BH-FDR) and from R2 onwards at the electrode cluster level (Fig. 2G; cluster permutation; R2: Σt = 192.2, p = 0.002; R3: Σt = 257.0, p < 0.001; R4: Σt = 113.8, p = 0.010; BH-FDR). As a result, changes in periodic Beta power differed significantly from aperiodic Beta power at every time point (Fig. 2F; Wilcoxon; R1: z = −2.84, r = −0.65, p = 0.005; R2: z = −3.40, r = −0.78, p < 0.001; R3: z = −3.57, r = −0.82, p < 0.001; R4: z = −2.41, r = −0.55, p = 0.016; BH-FDR).

Finally, periodic, aperiodic and total power in the Gamma band (30-45 Hz) all significantly increased from baseline, though with distinct timecourses. Aperiodic Gamma power rose earliest, with significant increases at R1 to R3 for the participant level (Fig. 2F; Wilcoxon; R1: r = 0.73, p = 0.001; R2: r = 0.59, p = 0.009; R3: r = 0.51, p = 0.026; BH-FDR), and at R1 and R2 for the cluster level (Fig. 2G; cluster permutation; R1: Σt = 119.7, p = 0.002; R2: Σt = 68.3, p = 0.016; BH-FDR). Complementary to this, periodic power emerged later, rising significantly from R2 onward (Fig. 2F,G; Wilcoxon; R2: r = 0.71, p = 0.002; R3: r = 0.63, p = 0.006; R4: r = 0.51, p = 0.025; cluster permutation; R2: Σt = 46.8, p = 0.002; R3: Σt = 16.3, p = 0.006; R4: Σt = 21.3, p = 0.003; BH-FDR). As a result, total Gamma power increased throughout time points R1–R3 (Fig. 2F; Wilcoxon; R1: r = 0.72, p = 0.002; R2: r = 0.62, p = 0.007; R3: r = 0.53, p = 0.021; cluster permutation; R1: Σt = 102.9, p = 0.0056; R2: Σt = 93.4, p = 0.008; R3: Σt = 30.0, p = 0.033; BH-FDR). As such, periodic Gamma power significantly exceeded aperiodic Gamma at R2 - R4 (Fig. 2F; Wilcoxon; R2: r = 0.52, p = 0.023; R3: r = 0.51, p = 0.025; R4: r = 0.51, p = 0.026; BH-FDR).

Together, these results not only suggest a dynamic and frequency-specific reorganization of oscillatory neural activity, they also mirror the pronounced desynchronization of broadband cortical activity and the drop of spectral Alpha power that has been consistently reported as a marker of psychedelic states (Carhart-Harris et al., 2016; Godfrey et al., 2025; Muthukumaraswamy et al., 2013; Timmermann et al., 2019). One conceivable explanation for these observations might be that the intensified breathing involved in HVB raises general physiological arousal, which could in turn modulate and possibly entrain cortical activity (Azzalini et al., 2019). Heart rate does elevate from baseline throughout active breath epochs, but is only significantly elevated during resting epochs R2 and R3, whereas breath rate is unchanged in R1 and R2 and decreases in R3 and R4 (Supp. Figs S3 and S4). Neither changes in heart rate nor respiratory rate correlated with ASC intensity in any resting-state epoch (Supp. Figs. S3 and S4), and the reported neural effects were present in epochs where neither was elevated, suggesting acute cardiac and respiratory activity alone are unlikely to account for the observed neural changes.

Beyond such direct physiological factors, these results raise the question to what extent the observed shifts in spectral power are triggered by the core breathing technique of HVB, and to what extent they are evoked by its context and setting. Specifically, HVB features two aspects that have been shown to generate significant neural effects in their own right: passive music listening (Aparicio-Terrés et al., 2025; Blood & Zatorre, 2001; Daly et al., 2014) and deliberate body awareness, i.e. interoception (Critchley et al., 2004; Villena-González et al., 2017). To disentangle the overall effects of HVB from the contributions of these two central features, we conducted two control experiments: In the first, a subgroup of HVB participants (n = 10) additionally attended a Progressive Muscle Relaxation (PMR) session (see Supp. Text S4); in the second, an independent sample of breathwork-naive participants (n = 8) attended a passive music listening session featuring the same music playlist as the HVB sessions themselves.

Both control conditions followed the same experimental structure as HVB sessions, but resulted in significantly lower real-time ASC scores than HVB, and distinct profiles of ASC survey dimensions compared to HVB (Supp. Figs. S5 - S6; Supp. Tables S5 - S6; see also Supp. Fig. S7 for distinct post-session changes in affect evoked by PMR). At the neural level, none of the spectral effects of HVB were replicated in either control condition. Instead, both control conditions showed unique neural effects that were unrelated to, and in some instances opposite to, those observed for HVB. PMR significantly decreased aperiodic power in the Theta, Alpha, Beta and Gamma bands (Supp. Fig. S8; Supp. Table S7), while music listening resulted in decreased aperiodic power in the Beta and Gamma bands (Supp. Fig. S9; Supp. Table S8). Neither control condition evoked periodic spectral changes. Together, these controls suggest that the spectral signature of HVB is not explained by facilitator presence, guided interoceptive practice or music alone.

### HVB increases neural complexity in frequency-specific ways

Given that HVB predominantly modulated periodic rather than aperiodic spectral power across multiple frequency bands, we hypothesized that this would further dissolve the temporal structure of neural activity, potentially diminishing rhythmic elements and thereby increasing signal complexity in a similar way as reported for psychedelics (Carhart-Harris, 2026; Kroupi et al., 2026; Lebedev et al., 2016; Schartner et al., 2017). To test this hypothesis, we computed Lempel-Ziv complexity (LZC; Lempel & Ziv, 1976; see Methods) for broadband EEG signals and for EEG signals filtered according to the canonical frequency bands applied in Figs. 2F-G.

HVB produced a pronounced increase of LZC in broadband signals across all time points (Fig. 3A; all analyses use Wilcoxon signed-rank tests on modulation indices based on matched-pairs rank-biserial r; n = 25; R1: r = 0.66, p = 0.004; R2: r = 0.82, p < 0.001; R3: r = 0.63, p = 0.006; R4: r = 0.54, p = 0.019; significance BH-FDR corrected across four epochs; see Supp. Table S9 for full statistics), with significant sensor clusters distributed across most of the scalp (Fig. 3B; cluster-based permutation test; n = 25; 5,000 permutations; R1: Σt = 177.86, p = 0.002; R2: Σt = 214.43, p < 0.001; R3: Σt = 126.00, p = 0.010; R4: Σt = 95.29, p = 0.020; BH-FDR; see Supp. Table S9).

**Figure 3:**
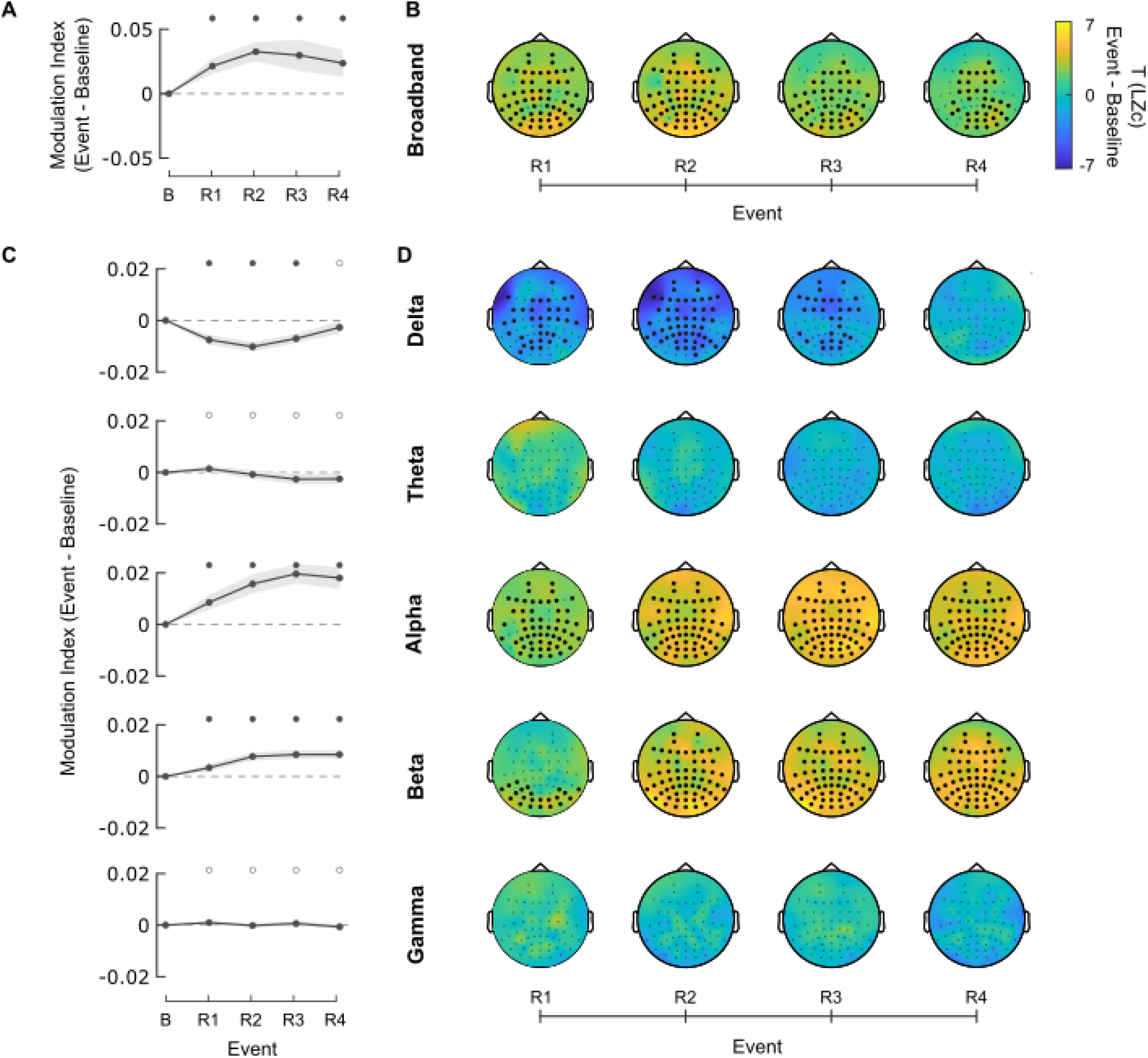
Lempel–Ziv complexity changes dynamically during HVB sessions. **A)** Average modulation index of LZC relative to baseline, based on broadband EEG signals. Shaded region: SEM. Filled circles above panel indicate significance of signed-rank tests (p<0.05) as in Fig. 2B. **B)** Same topological map as in Fig. 2C. Color: T value of population-level difference in broadband LZC between R1–R4 and baseline (see in-figure color bar). Black markers identify electrodes that belong to statistically significant spatial clusters, based on two-sided cluster-permutation inference and BH-FDR. **C)** Same as Fig. 2A for canonical frequency bands (see in-figure labels). **D)** Same as Fig. 2B for canonical frequency bands.

Decomposing this broadband effect into frequency bands revealed frequency-specific changes in LZC that complemented the changes in spectral power described previously. Counter to the broadband increase described above, LZC actually decreased in Delta frequencies across R1–R3 and the entire scalp (Figs. 3C-D; Wilcoxon: R1: r = −0.71, p = 0.002; R2: r = −0.90, p < 0.001; R3: r = −0.70, p = 0.002; cluster permutation: R1: Σt = 102.40, p = 0.002; R2: Σt = 183.51, p < 0.001; R3: Σt = 86.57, p < 0.001; BH-FDR; Supp. Table S9). In contrast, LZC in Alpha increased drastically across all four time points and the entire scalp (Figs. 3C-D; Wilcoxon; R1: r = 0.61, p = 0.007; R2: r = 0.76, p < 0.001; R3: r = 0.88, p < 0.001; R4: r = 0.81, p < 0.001; cluster permutation: R1: Σt = 141.98, p = 0.007; R2: Σt = 240.06, p < 0.001; R3: Σt = 288.89, p < 0.001; R4: Σt = 240.63, p < 0.001; BH-FDR). Beta frequencies largely followed the same pattern of globally raised LZC as Alpha, with a somewhat slower onset which was initially focused on posterior regions before shifting into a widely distributed pattern (Figs. 3C-D; Wilcoxon: R1: r = 0.46, p = 0.042; R2: r = 0.80, p < 0.001; R3: r = 0.82, p < 0.001; R4: r = 0.86, p < 0.001; cluster permutation: R1: Σt = 59.35, p = 0.026; R2: Σt = 239.32, p < 0.001; R3: Σt = 249.30, p < 0.001; R4: Σt = 267.32, p < 0.001; BH-FDR). Neither Theta nor Gamma bands showed significant shifts in LZC (Supp. Table S9).

Neither of our two control conditions reproduced this pattern of HVB-related LZC changes. In the music control, LZC did not change significantly in broadband or in any frequency band (Supp. Fig. S9 and Supp. Table S10). In the PMR control, the only effect was a modest increase in Alpha-band LZC at the participant level, which did not reach significance at the sensor-cluster level (Supp. Fig. S10; Supp. Table S11).

These results indicate that HVB loosens the overall temporal structure of neural activity, thereby increasing signal complexity in similar ways as psychedelics (Carhart-Harris, 2026; Carhart-Harris et al., 2016; Kroupi et al., 2026; Lebedev et al., 2016; Lyons et al., 2026; Schartner et al., 2017). These effects are frequency-specific and mirror the shifts in periodic spectral power shown in Fig. 2, with neural activity in the Delta band actually becoming more rhythmic and less complex, while activity in Alpha and Beta bands becomes less rhythmic and more complex, contributing to a boost in overall signal complexity.

### HVB reorganizes functional connectivity

Next, to test whether HVB alters cross-cortical organization, as has been reported for psychedelic states (Alamia et al., 2020; Carhart-Harris & Friston, 2019; Girn et al., 2022; Preller et al., 2018), we computed the weighted phase-lag index (wPLI; Potash et al., 2025; Vinck et al., 2011) to track phase synchronization and connectivity across electrode locations.

Broadband connectivity was largely unaffected by HVB, with wPLI showing no significant changes across all four time points (Fig. 4A; Supp. Table S12). Instead, our analysis highlighted frequency-specific shifts in phase synchronization. Across the Delta and Theta bands, wPLI nodal connectivity strength increased across the session, with significant positive clusters emerging throughout R1–R4 (Fig. 4A; cluster-based permutation test; n = 25; 5,000 permutations; Delta: R1: Σt = 21.26, p = 0.028; R2: Σt = 97.64, p = 0.006; R3: Σt = 106.79, p < 0.001; R4: Σt = 74.33, p = 0.002; Theta: R1: Σt = 143.94, p < 0.001; R2: Σt = 68.08, p = 0.004; R3: Σt = 61.75, p = 0.005; R4: Σt = 114.55, p < 0.001; BH-FDR across four epochs; Supp. Table S12). Conversely, wPLI nodal strength in Alpha decreased significantly towards the second half of the session, with negative clusters at R3 and R4 located particularly in posterior regions (Fig. 4A; cluster permutation; R3: Σt = 128.55, p = 0.012; R4: Σt = 138.67, p = 0.004; BH-FDR). wPLI did not change significantly in Beta and Gamma bands (Supp. Table S12).

**Figure 4:**
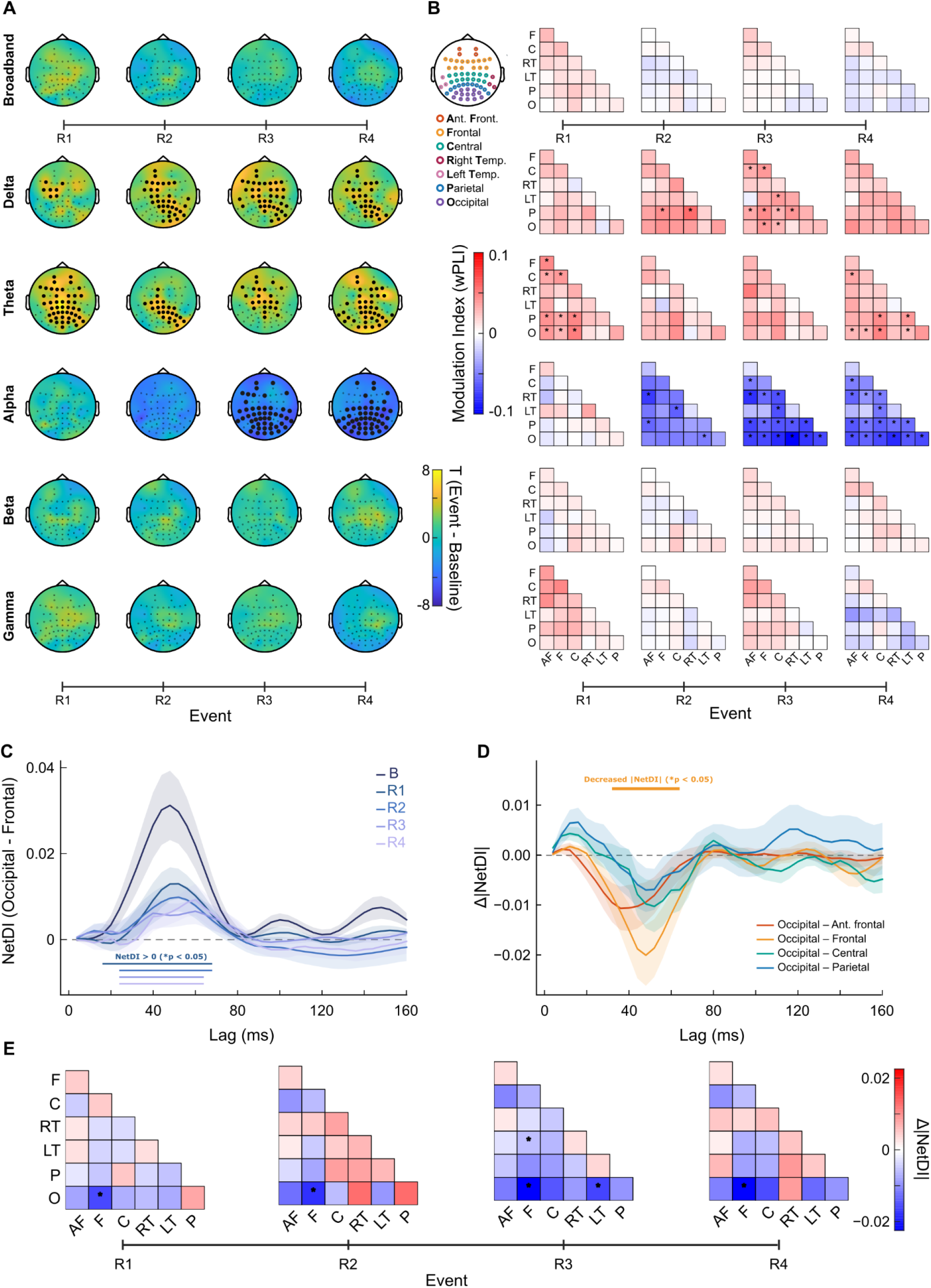
Phase-lagged connectivity and directional symmetry. **A)** Event-minus-baseline t (24) maps of wPLI nodal strength. Markers identify significant node clusters after resting-map BH-FDR correction within band and sign. Colors represent t values = −7 to +7. **B)** Mean modulation index of wPLI for 21 unique ROI edges; stars denote q <0.05 across edges within each band/recording. **C)** Mean ± SEM signed occipital–frontal NetDI over 4–160 ms. Positive values favor occipital-to-frontal lagged information. Bars mark resting-versus-baseline clusters retained after correcting all ten recording comparisons. **D)** Mean ± SEM resting-averaged Δ|NetDI| for four occipital connections, each tested against zero. The bar identifies the frontal–occipital reduction (q = 0.0272 across four connections). **E)** Group-selected peak Δ|NetDI| at R1–R4. Stars denote lag-curve tests retained after correcting four recordings within each pair. Matrices are symmetric and diagonals undefined; negative values mean weaker directional imbalance than baseline.

While this analysis quantifies the average connectivity at each electrode location, we also explored the pair-wise functional connectivity between brain regions. To this end, we defined seven canonical brain regions (see in-figure legend in Fig. 4B and Supp. Table S13 for region definitions) and computed the modulation index for pair-wise wPLI across time points R1-R4. Overall, pair-wise regional connectivity followed the frequency-specific changes of per-electrode connectivity highlighted in Fig. 4A. Specifically, pair-wise broadband wPLI did not change significantly, and neither did pair-wise wPLI in the Beta and Gamma bands (Supp. Table S14). At the same time, pair-wise wPLI increased in Delta and Theta, and decreased in Alpha (Fig. 4B; Wilcoxon signed-rank tests on modulation indices, matched-pairs rank-biserial r; n = 25; Delta: R2: 2/21 edges, r = 0.80–.82, p < 0.001; R3: 9/21 edges, r = 0.53–.80, p < 0.021; Theta: R1: 9/21 edges, r = 0.56–.79, p < 0.015; R4: 7/21 edges, r = 0.58–.78, p 0.012; Alpha: R2: 4/21 edges, r = −.59 to −.70, p < 0.009; R3: 16/21 edges, r = −.50 to −.81, p < 0.028; R4: 16/21 edges, r = −.49 to −.70, p = 0.032; significance corrected using BH-FDR across 21 pairwise comparisons within each band and epoch; Supp. Table S14).

Closer analysis revealed several interesting patterns: first, wPLI in Delta and Theta bands mainly increased between posterior (i.e. electrodes assigned as occipital or parietal) and anterior-central regions (frontal, anterior frontal, and central electrodes), as well as within anterior regions (see Girn et al., 2026). At the same time, wPLI in the Alpha band reflected a complete desynchronization of posterior regions from the rest of the brain at R3 and R4, a pattern that also largely applied to right-temporal electrode locations (Fig. 4B; Supp. Table S14).

Neither control condition reproduced HVB-related changes in connectivity. In the PMR control, wPLI nodal strength did not change significantly in any frequency band (Supp. Fig. S10; Supp. Table S15). In the music control, theta connectivity increased at R1 and R4 at the participant level and at R1 at the cluster level, with no other significant changes, and without the sustained cluster-level Theta increases seen during HVB (Supp. Fig. S9; Supp. Table S16).

### HVB increases symmetry of frontal-occipital communication

Based on the shifts in functional connectivity highlighted by our wPLI analyses, we next explored if the directionality of functional connections was also impacted by HVB, mirroring declines in hierarchical processing reported for psychedelic states (Alamia et al., 2020; Girn et al., 2022, 2026; Preller et al., 2018). To this end, we computed Net Directed Information (NetDI; Ince et al., 2017; Potash et al., 2025). NetDI is a metric of directional information transfer between two regions, computed as the difference between the transferred information Region A → Region B versus Region B → Region A (see Methods).

Fig. 4C shows an example of the NetDI between occipital and frontal regions. At baseline, occipital-frontal NetDI shows a pronounced peak at a lag of approx. 48 ms, indicating that activity in occipital regions drives activity in frontal regions at a small delay. This peak is significantly attenuated for time points R1 - R4, suggesting that communication between occipital and frontal cortices becomes less unidirectional and more symmetric, consistent with a flattening of cortical hierarchy (Fig. 4C; sign-flip cluster-based permutation tests of each epoch against baseline at peak delay; n = 25; 5,000 permutations; R1: peak lag 44 ms, mean Δ = 0.019 bits, cluster 16–68 ms, cluster mass = 47.03, p <0.001; R2: 48 ms, 0.022 bits, 24–68 ms, cluster mass = 40.51, p = 0.002; R3: 48 ms, 0.026 bits, 24–64 ms, cluster mass = 40.24, p = 0.002; R4: 48 ms, 0.025 bits, 24–64 ms, cluster mass = 39.10, p <0.001; significance corrected using BH-FDR across 21 region pairs × 4 epochs; Supp. Tables S17–S18). To test if this effect was also apparent in other cross-cortical connections, we computed the absolute change in NetDI between baseline and an average of R1 - R4.

Fig. 4D shows the change in NetDI between occipital cortices and four other regions (see in-figure legend). For all four regions, NetDI decreased during HVB, but this decrease was only significant for occipital–frontal communication at time lags of 32–64 ms (Fig. 4D; cluster-based permutation test on the absolute change in NetDI, baseline vs mean R1–R4, n = 25; 5,000 permutations; occipital–frontal, 32–64 ms: cluster mass = −25.53, mean Δ = −0.0142 bits, p = 0.007; significant after BH-FDR across four region pairs; Supp. Table S19; see Supp. Fig. S11 for all hierarchical relationships between ROIs).

Fig. 4E shows NetDI compared to baseline at the group-selected peak lag for all pairings of cortical regions. While asymmetric connectivity between occipital and frontal regions declines overall across the HVB session, a significant decrease in connectivity also occurs between left temporal regions and occipital/ frontal regions occurs at R3 (Fig. 4E; cluster-based permutation tests of change in |NetDI|, n = 25, 5,000 permutations; 21 region pairs × 4 epochs; occipital–frontal: R1: 36–72 ms, cluster mass = −27.85, p = 0.006, R2: 32–64 ms, −24.58, p = 0.009, R3: 32–64 ms, −26.20, p = 0.006, R4: 32–60 ms, −22.15, p = 0.008; occipital–left temporal at R3: 36–76 ms, −35.13, p = 0.002; frontal–left temporal at R3: 28–64 ms, −25.37, p = 0.006; BH-FDR correction across R1–R4 within each pair; see Supp. Table S19 and Supp. Fig. S11). As such, HVB weakened directed, hierarchical information flow, most clearly between posterior-anterior connections. This is consistent with the reorganization of cortical hierarchy reported for psychedelic states (Alamia et al., 2020; Carhart-Harris & Friston, 2019; Girn et al., 2022, 2026; Preller et al., 2018). Neither control condition (PMR or music listening) evoked significant changes in directed information flow (Supp. Figs. S8 and S9).

### Neural signatures of HVB relate to real-time but not retrospective ratings of ASC intensity

Based on the insight that HVB triggered consistent shifts in spectral power, signal complexity, and connective symmetry, all of which mirrored effects of classical psychedelics to a large extent, we next assessed if these neural shifts were directly related to subjective ASC experiences. To this end, we first explored the correlation between neural markers within individual resting-state epochs and the real-time ASC ratings given at the start of the same epoch.

Real-time ASC ratings at the start of each resting state epoch were significantly predicted by a range of neural metrics during the epoch, specifically decreased oscillatory activity and increased signal complexity in the broadband, Alpha and Beta ranges, as well as increased oscillations and decreased signal complexity at Delta frequencies (Fig. 5A, top; participant level repeated measures-correlations; n = 25 participants x 5 time points; df = 99; periodic power: broadband: r = −0.476, p < 0.001; Delta: r = 0.254, p = 0.010; Alpha: r = −0.607, p < 0.001; Beta: r = −0.408, p < 0.001; LZC: broadband: r = 0.338, p < 0.001; Delta: r = −0.240, p = 0.016; Alpha: r = 0.550, p < 0.001; Beta: r = 0.525, p < 0.001; effects significant after BH-FDR correction across 25 neural metrics tested; see Supp. Table S20). Additional predictors of higher ASC ratings were increased wPLI connectivity in Delta and decreased wPLI connectivity in Alpha (Fig. 5A; repeated-measures correlations; Delta: r = 0.294, p = 0.003; Alpha: r = −0.294, p = 0.003; BH-FDR), as well as an overall decrease in cross-region asymmetry (Fig. 5A, top; Δ|NetDI|; repeated-measures correlations; n = 25 participants x 4 time points, df = 74; r = −0.370, t(74) = −3.42, p = 0.001; BH-FDR).

**Figure 5:**
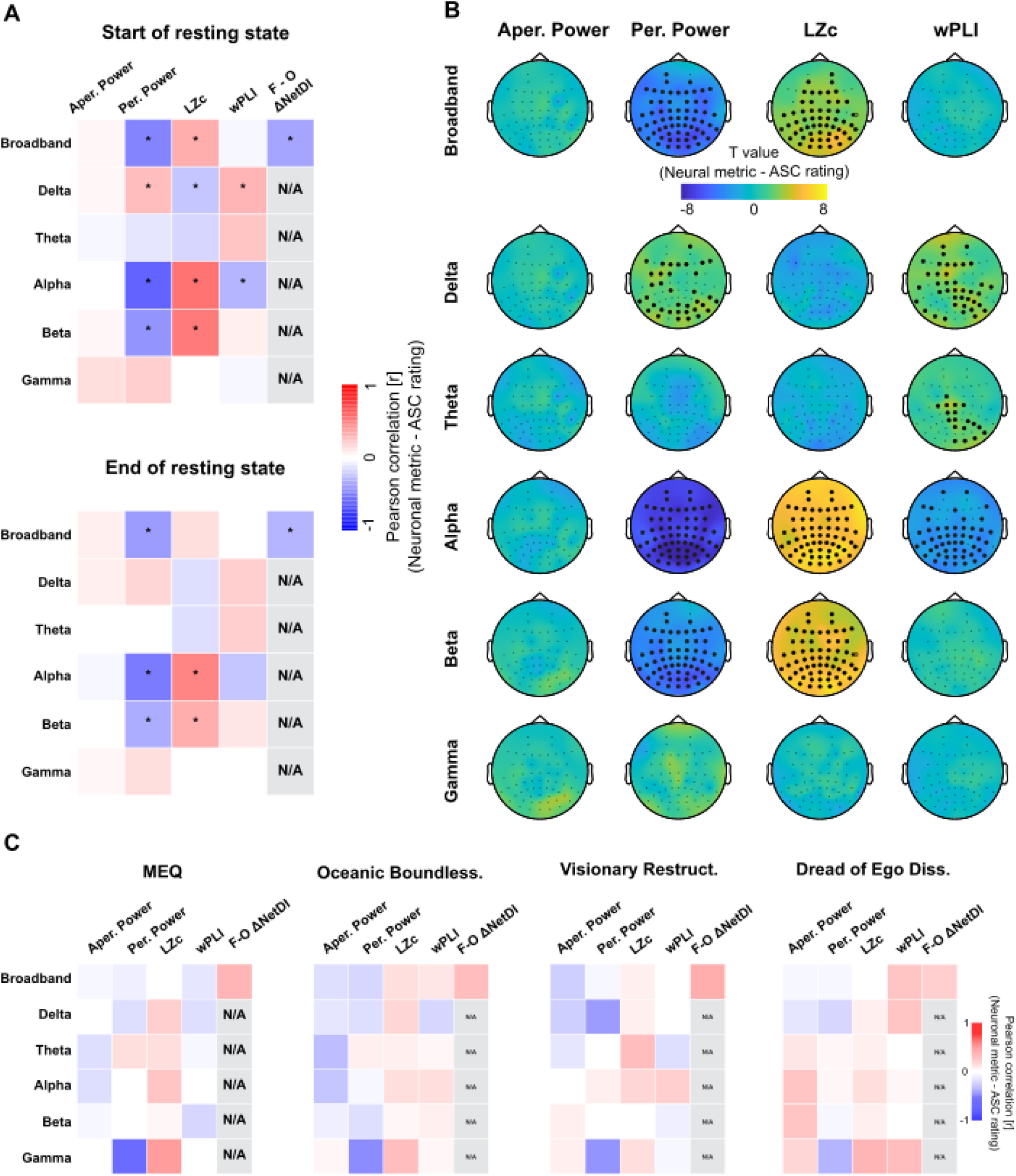
Associations between frequency-specific neural metrics and experiential ratings. **A)** Repeated-measures correlations (rm-corr; within-participant) between ASC ratings at the start of a resting-state epoch and neural metrics for the same epoch. Colors indicate the repeated-measures correlation coefficient (see in-figure color bar). Asterisks denote correlations that were statistically significant at p < 0.05 after BH-FDR correction for multiple comparisons. Aperiodic power, periodic power, LZC, and wPLI are computed across baseline and R1 - R4 epochs (n = 25 participants x 5 time points = 125). Peak frontal–occipital Δ|NetDI| applies to epochs R1–R4, as Δ|NetDI| was computed relative to baseline (n = 25 participants x 4 time points = 100). Correlations for each of these metrics were evaluated for broadband activity as well as the canonical delta-gamma frequency bands. **B)** Same as Fig. 4A for the correlation coefficients between ASC ratings at the start of resting-state epochs and the corresponding neural markers (see in-figure labels). Topographies show sensor-level t-statistics (sensor-level cluster-based permutation test; see in-figure color bar). Black markers identify significant electrode clusters retained after participant-blocked permutation testing and BH-FDR correction across six bands. **C)** Same as A for Pearson’s correlation coefficients (Spearman’s correlation coefficient for DED reported in Supp. Table S20) between retrospective survey subscale and neural metrics at each participant’s peak-ASC recording (n = 25).

Cluster-level results largely mirrored the participant-level correlations (Fig. 5B). Decreased periodic power in the broadband, Alpha and Beta ranges and increased LZC in the broadband, as well as decrease wPLI in the Alpha band predicted ASC intensity at the cluster level. Broadband, Alpha and Beta ranges were each supported by clusters spanning most of the scalp (sensor level permutation tests; n = 25; df = 99; periodic power: broadband, 62 sensors, Σt = −257.81, p = 0.0002; Alpha, 63 sensors, Σt = −431.24, p = 0.0002; Beta, 62 sensors, Σt = −230.87, p = 0.0002; LZC: broadband, 56 sensors, Σt = 181.59, p = 0.0004; Alpha, 63 sensors, Σt = 358.53, p = 0.0002; Beta, 63 sensors, Σt = 310.39, p = 0.0002; Alpha wPLI: 51 sensors, Σt = −140.46, p = 0.005; BH-FDR). Delta LZC did not reach significance at the cluster level, and Delta periodic power and wPLI clusters were smaller than broadband and Alpha (cluster permutation; periodic Delta power: 36 sensors, Σt = 89.45, p = 0.0004; wPLI: 39 sensors, Σt = 101.74, p = 0.0006; BH-FDR). Theta wPLI additionally showed a significant cluster, largely in posterior electrodes (Fig. 5B; cluster permutation; 18 sensors, Σt = 43.48, p = 0.010; BH-FDR). Real-time ASC ratings at the start of each resting-state epoch were also significantly predicted by a decrease in the aperiodic exponent, with a lower exponent accompanying higher ASC intensity (cluster permutation; n = 25; negative cluster; p = 0.0144; see Supp. Fig. S13).

Qualitatively, the same pattern was observed when predicting real-time ASC ratings at the end rather than start of each resting-state epoch, but associations were generally somewhat weaker, and reached significance for fewer metrics (Fig 5A, bottom; see Supp. Fig. S14). Change in ASC intensity throughout an epoch (ASC end - ASC start) does not correlate with any metric, suggesting absolute ASC intensity, rather than dynamic ASC intensity shift, correlates to our observed neural changes (see Supp. Fig. S14).

Together, these results suggest that deep subjective ASCs were mainly predicted by an overall decrease in the temporal and spatial structure of neural activity, as well as a frequency-specific decrease of structure and power in Alpha frequencies, and a complementary increase of structure and power in Delta frequencies. Interestingly, in both control conditions, ASC intensity only correlated to a decrease in the aperiodic power of higher frequency bands, and additionally an increase in the aperiodic exponent in the music control (Supp. Fig. S12; Supp. Fig. 13).

To test if these relations also extended to the specific experiential contents of ASCs, as reported post-session via the 11D-ASC and MEQ30, we computed Pearson’s correlations between the same neural metrics tested above for each participant’s resting-state epoch at their respective highest ASC rating (see Methods), and the corresponding post-session scores for the MEQ30 as well as the three sub-scales of the 11D-ASC. We found no significant correlations between neural metrics and survey scores (Pearson’s correlations; n = 25; 25 tested neural metrics per survey; all |r| ≤ 0.57, all p ≥ 0.003, none significant after BH-FDR across 25 neural metrics tested within each survey; see Supp. Table S20). These findings suggest that neural dynamics were more closely linked to real-time assessments of subjective experience than to retrospective assessments.

## Discussion

In this study, we report that HVB not only evokes subjective experiences akin to classical psychedelics, but also modulates neural activity in comparable ways. Specifically, we found several markers of dissolving temporal structure, including reduced broadband power, and increased broadband signal complexity; as well as a redistribution of spectral power composition, with activity in Alpha (8-13 Hz) and Beta (13-30-Hz) bands losing spectral power and gaining complexity, while activity in Delta (1-4 Hz) and Gamma (30-45 Hz) gained spectral power and lost complexity. At the spatial level, we found a reconfigured constellation of functional connectivity and hierarchical information flow. Together, these effects closely mirror typical neural signatures of classical psychedelics (Gattuso et al., 2023; Godfrey et al., 2025; Kwan et al., 2022; Lebedev et al., 2016; Schartner et al., 2017; Tagliazucchi et al., 2014; Tolle et al., 2024; Yu et al., 2024). The observed neural effects were not explained by individual features of HVB, specifically the presence of music or deliberate interoception (see Supp. Figs S8-10 and Supp. Tables S7-S8, S10-11, S15-16). Instead, they robustly predicted real-time changes in reported ASC depth. These findings are consistent with previous work by Lewis-Healy et al. (2024), who reported an increase in LZC for portable EEG recordings collected during SOMA breathwork. Our time-resolved analysis of high-density EEG signals and real-time ASC ratings allows us to expand on these findings by highlighting ongoing links between neural metrics and experiential states during an HVB session.

### Set and setting

In interpreting these results, it is important to take into account the contribution of contextual variables. While HVB practices are centered around a change in breathing rhythm towards deeper, accelerated breathing, they also typically feature emotionally evocative music, as well as supportive facilitation including consensual bodywork and guided interoception. In addition, this study was conducted in experienced HVB practitioners, whose past HVB experiences may have set expectations that further modulated the session. As such, the effects reported here cannot be seen as generated by a breathing technique in isolation, but by a holistic intervention incorporating intensified breathing, music, interpersonal support as well as the participants’ own intentions and expectations. This resembles the ‘set and setting’ principle established in psychedelic practice, whereby the outcome of a psychedelic intervention does not just depend on its pharmacological action but also on the ‘set’ i.e. the participant’s mind set, and the ‘setting’, i.e. the environmental context (Brouwer & Carhart-Harris, 2021; Carhart-Harris et al., 2018; Hartogsohn, 2016, 2017).

While ‘set and setting’ variables certainly contribute to HVB experiences, and are in fact deliberately included in HVB practices e.g. in the form of pre-session intention setting, evidence suggests that these contextual variables are not the main driver of the effects reported here. First, by analysing neural and experiential dynamics only for resting-state epochs interleaved throughout the session, we removed acute effects of music, facilitation and intensified breathing from our measurements. Second, two control conditions - music listening and progressive muscle relaxation (PMR), which features deliberate interoception in the presence of a facilitator - yielded significantly lower ASC scores and none of the neural changes reported for HVB (Supp. Figs S5-S6 and Supp. Tables S5-S6). Although these control conditions had smaller sample sizes than the total HVB sample, they were nevertheless specifically designed to emulate the contextual variables of HVB as closely as possible by taking place in the same setting, with the same temporal structure of recording sessions, and featuring the same music (for music listening) and the same facilitator-participant dyad (for PMR). Finally, a matched breathwork sample to the PMR condition (n = 9) reproduced the main effects seen in the total HVB sample (albeit with smaller effect sizes), indicating that sample size alone does not explain the differences, and supporting the validity of the comparison (see Supp. Fig. S8, S10; Supp. Tables S7, S11, S15). As such, it is striking to note that the experiential and neural effects of HVB failed to replicate qualitatively or quantitatively in either control condition.

In both controls, real-time ASC intensity was significantly lower than for HVB, and correlated mainly with decreased aperiodic power in higher frequency bands, an effect that occurred in the opposite direction for HVB. This suggests that ASCs in control sessions may have been linked to a shift towards decreased cortical excitability (Gao et al., 2017), a completely separate mechanism for deviation from ordinary consciousness compared to HVB. These findings are also in line with prior work that estimated the contribution of contextual variables to the experiential effects of HVB at approx. 30%, suggesting that context is not the core driver of HVB experiences (Canello et al., 2026; Havenith et al., 2025).

At the same time, the effects of HVB also cannot be reduced to a simple physiological response to controlled hyperventilation. Voluntary hyperventilation has been reported to produce diffuse EEG slowing and an increase in Delta power, attributed to hypocapnia-driven reductions in cerebral perfusion (Bednarczyk et al., 1990; Debatisse et al., 2024; Kraaier et al., 1988). As such, while HVB-driven hypocapnia may contribute to the observed increase of Delta power, it does not explain any of the other neural changes reported here. These findings suggest that while the HVB practice studied here combines voluntary hyperventilation, music, deliberate interoception and facilitation, its effects cannot be understood as a simple sum of these components. Rather, HVB seems to act as a holistic trigger condition for extensive deviations from ordinary neural and conscious states, which are synergistically supported by its components.

### Neural effects of HVB

At the neural level, HVB triggered a pronounced shift in activity, characterized by a global loosening of temporal structure, and a reorganization of spatial structure. Spatial reorganization appears in three ways: (1) the pronounced reduction of Alpha oscillations and anterior-posterior Alpha-band coupling, which suggests reduced integrity within the DMN (see below), (2) concurrent increases in low frequency connectivity, possibly reflecting an increase in long-range coupling (Buzsáki & Draguhn, 2004; Von Stein & Sarnthein, 2000), and (3) decreased directionality of posterior-to-anterior information flow, indicating less hierarchical cross-cortical communication. This reduction in hierarchical differentiation and directional asymmetry echoes reports of weakened processing hierarchies (Gattuso et al., 2023; Pines et al., 2026; Shinozuka et al., 2025) and reduced differentiation between unimodal and transmodal cortex under LSD and psilocybin (Girn et al., 2022, 2026), and the altered travelling waves observed under DMT (Alamia et al., 2020).

Temporally, the attenuation of neural structure appears as a broadband (1-45 Hz) decrease in rhythmic activity (Fig. 2), concurrent with an overall increase of LZC (Fig. 3), signalling a shift towards more diverse, less rhythmic or otherwise predictable neural dynamics. This effect echoes decreases in broad band power and increases in broadband signal complexity observed consistently across classical psychedelics (Lebedev et al., 2016; Pallavicini et al., 2019; Schartner et al., 2017; Subramani et al., 2026; Timmermann et al., 2019), as well as in a previous study of HVB (Lewis-Healey, 2024). In addition, the aperiodic exponent of spectral power (Donoghue et al., 2020) was significantly reduced during HVB, and this flattened exponent correlated with higher real-time ASC intensity. Such flattened power spectra have been proposed to reflect a shift towards increased cortical excitability (Gao et al., 2017), and have also been observed during LSD exposure (Muthukumaraswamy & Liley, 2018; Subramani et al., 2026). Together, enhanced cortical excitability and more diverse, less rhythmic signalling evoke the notion of a more entropic, less constrained regime of cortical processing, as encapsulated by the ‘Entropic Brain’ concept popular in psychedelic research (Carhart-Harris, 2026; Carhart-Harris et al., 2014; Lyon, 2026).

In addition to this overall spatiotemporal disintegration, HVB also produces striking frequency-specific reorganization. Most strikingly, neural activity in the Alpha band was drastically desynchronized, with an extensive loss of oscillatory power and corresponding rise in signal complexity, accompanied by a decline of cross-cortical Alpha coupling, particularly along the posterior-anterior axis. Beta-band activity largely mirrored these changes at a somewhat lower level, and without corresponding connectivity changes. These effects were counterbalanced by enhanced oscillatory power and cross-cortical connectivity, and decreased signal complexity, in the Delta band, and increased connectivity in the Theta band, indicating a more structured oscillatory drive in these lower frequencies.

The observed weakening of Alpha and Beta oscillations is particularly interesting because oscillations in both frequency bands have been proposed to represent maintenance of the cognitive “status quo”, marking processes that carry top-down predictions towards sensory and sensorimotor areas to gate irrelevant sensory inputs (Engel & Fries, 2010; Jensen & Mazaheri, 2010; Klimesch et al., 2007; Michalareas et al., 2016). As such, both frequency bands are attenuated by novel, unexpected and salient events (Engel & Fries, 2010; Ferrari et al., 2022; Trautwein et al., 2024), as well as during psychedelic states (Alamia et al., 2020; Barrett & Griffiths, 2017). In line with this, Alpha oscillations, as well as frontoparietal coupling in the Alpha band, have been proposed as neural markers signalling involvement of the Default-Mode-Network (DMN; Guldenmund et al., 2012; Knyazev, 2013; Knyazev et al., 2011; Mo et al., 2013), a widespread network of brain regions which jointly support the maintenance of a coherent sense of self, including thoughts about personal history, future plans and perception of self by others (Babo-Rebelo et al., 2016; Davey et al., 2016; Menon, 2023; Yeshurun et al., 2021).

In this context, a sustained decrease in oscillatory activity and cross-region coupling in the Alpha range is consistent with a relaxation of DMN-related top-down feedback and sensory gating, releasing neural dynamics from their usual constraints (Carhart-Harris & Friston, 2019; Gattuso et al., 2023; Van Elk et al., 2019). At the experiential level, such a shift is expected to reduce the influence of habitual processing patterns and of self-referential thinking, creating space for experiences of heightened sensory vividness, dissolving boundaries of self and perspective change (Carhart-Harris & Friston, 2019; Gattuso et al., 2023; Jedlicka & Havenith, 2025). Consistently with this, reduced functional connectivity of the DMN has recently been observed to predict stronger ASCs during HVB (Schmitz et al., 2026).

Interestingly, these observations stand in contrast to other forms of ASCs such as meditative practices and flow states, which have instead been linked to increased Alpha oscillations (Katahira et al., 2018; Kiely & Gellhorn, 1972; Lomas et al., 2015; Timmermann, Barba, et al., 2025). In these contexts, increased Alpha power is thought to mediate the inhibition of external sensory input in favour of internally directed processing (Cooper et al., 2003; Kiely & Gellhorn, 1972). From this perspective, meditative states may be understood as a processing regime in which sensory input is held in line with top-down priors, whereas psychedelic and breathwork states appear to relax those priors, opening the system to novel and unexpected input, and thereby providing a basis for long-term shifts in perspective (Carhart-Harris & Friston, 2019; Erritzoe et al., 2019; Lebedev et al., 2016; MacLean et al., 2011).

The complementary increase of oscillatory communication in Delta frequencies may be supported by a conjunction of factors. First, enhanced Delta oscillations are a known consequence of controlled hyperventilation (Kraaier et al., 1988), attributed to reductions in cerebral blood flow and increases in neuronal excitability triggered by hyperventilation-induced hypocapnia (Gotoh et al., 1965). In addition, Delta oscillations could also be supported by cortical entrainment to cardiac rhythms (Park & Blanke, 2019; Tort et al., 2018), as heart rate during HVB tends to vary from approx. 60-100 beats per minute (see Supp. Fig. S3), all of which would fall into the Delta range.

Beyond these physiological factors, Delta oscillations have also been associated with disengagement from external sensory input, dominating for instance as sleep deepens (Harmony, 2013). However, Delta oscillations during sleep are typically accompanied by a steepening of the 1/f power spectrum (Lendner et al., 2020), indicating reduced cortical excitability (Gao et al., 2017), whereas during HVB, the power spectrum flattened. This constellation observed during HVB would be in line with a conscious state that is somewhat removed from the perception of external sensory stimuli, while still actively engaged in cognitive processing.

Finally, together with the reshuffling of spectral power from medium towards slow frequencies, activity in the Gamma band was also dynamically reorganized. Aperiodic Gamma power increased in the first half of the session, potentially reflecting a general rise of cortical excitability (see below). The second half of the session was then characterized by increased periodic Gamma power. Given that Gamma oscillations are thought to carry feed-forward information computed by local sensory circuits (Buzsáki & Wang, 2012; Michalareas et al., 2016), increased Gamma oscillations are consistent with a shift towards bottom-up processing (Fincham et al., 2023; Van Kerkoerle et al., 2014). It is however important to note that increased Gamma power was not accompanied by significant changes in Gamma LZC or cross-cortical connectivity, suggesting that it may not be the main factor driving a shift in cross-regional processing trajectories.

While the profile of neural shifts reported here strongly resembles the neural signatures of classical psychedelics, interestingly the closest matches appear to be with DMT or 5-MeO-DMT, particularly given the observed increases in Delta and Gamma power (see Blackburne et al., 2025; Godfrey et al., 2025; Timmermann et al., 2019; Timmermann, Sanders, et al., 2025). This suggests that the neuronal mechanisms engaged by HVB may be more similar to those of DMT and related tryptamines than to other psychedelics, a parallel of particular interest given evidence that DMT is upregulated by, and provides neuroprotection against, hypoxic environments (Schimmelpfennig & Jankowiak-Siuda, 2025; Szabo et al., 2016; Vaccari et al., 1978), such as the one created in the brain by HVB (Kartar et al., 2025; Schmitz et al., 2026).

Notably, none of the neural dynamics discussed so far were replicated in our two control conditions of music listening and PMR (Supp. Figs S8 - S10). The only neural signature of HVB that was mirrored by our control experiments was a moderate increase in cross-cortical Theta coupling, which only appeared during music listening. Increases in frontal theta synchronisation have previously been linked to music-evoked pleasure and emotional engagement (Ara & Marco-Pallarés, 2020; Sammler et al., 2007), suggesting that this neural effect may echo the auditory and affective dynamics evoked by music during the active breathing epochs, which may have persisted at least somewhat into our resting-state recordings. As increased Theta coupling was among the findings that diverged most from the typical neural signatures of psychedelic states, it might reflect a music-related component of HVB rather than of HVB-evoked ASCs. Together, the results of both control experiments indicate that the neural signatures of HVB reported here are largely not attributable to a simple linear summation of its component parts.

### Linking neural and experiential effects of HVB

The neural effects of HVB reported here reliably predicted ASC intensity: real-time ASC ratings correlated robustly with several previously identified neural markers of HVB, specifically with decreased periodic power in Alpha and Beta but increased periodic power in Delta, as well as increased LZC in Alpha and Beta but decreased LZC in Delta and decreased asymmetry of posterior-anterior information flow. In contrast, the same neural metrics did not correlate with any of the retrospective ASC evaluations surveyed via the 11D-ASC and MEQ30. This intriguing discrepancy between real-time and post-hoc ASC ratings may arise from several different factors.

First, it may reflect a difference in the criteria by which ASCs are evaluated: during the session, participants were asked to rate exclusively the perceived intensity of an ASC, whereas post-session surveys honed in on specific experiential qualities like ineffability. Our results may suggest that the identified neural markers of HVB mainly reflect how *intensely* altered an ASC is, but not necessarily *in which way* it is altered, thus correlating better with pure intensity ratings than with specific experiential contents.

Complementary to this, the dissociation between real-time and retrospective ASC metrics may also simply be a matter of timing: given that real-time ratings by definition provide a snapshot of ongoing experience, they are arguably more faithful to the “experiencing self,” whereas retrospective assessments capture related-but-distinct information held by the “remembering self” (Kahneman & Riis, 2005). As such, aspects of HVB experiences that are most tightly linked to ongoing neural dynamics may not be the ones that are later accessed by the ‘remembering self’. This immediacy of neurophenomenological dynamics is further underlined by the fact that the relation between neural metrics and real-time ASC ratings had already somewhat faded by the end of each resting-state epoch, suggesting that even a delay of just four minutes was beginning to obscure the link between neural and experiential dynamics. This quick decay is in line with the observation that respiratory-driven EEG changes resolve within minutes (Hoshi et al., 1999). It also provides a potential explanation why previous work by Lewis-Healey et al., (2024) did not find significant links between time-resolved changes in EEG signal complexity and experience depth. Since their study collected time-resolved but retrospective ratings of experience depth via Temporal Experience Tracing (Jachs, 2021), measurements may well have been hampered both by their reliance on the ‘remembering self’, and by even minor temporal inaccuracies in the estimated time course of experience depth throughout the HVB session. These observations highlight the crucial importance of metrics that can track neurophenomenological effects of ASCs as they happen in real time.

Finally, the relation between neural dynamics and post-session ASC ratings may also be attenuated by the fact that both MEQ30 and 11D-ASC were originally designed to quantify psychedelic experiences (Barrett et al., 2015; Dittrich, 1998), and may thus fail to reflect central experiential elements unique to HVB. For instance, real-time ASC ratings shared only about 50% of variance with positively-valenced survey dimensions like Oceanic Boundlessness, Mystical Experience and Positive Mood, but did not significantly correlate with negatively-valenced elements like Dread of Ego Dissolution. This suggests that while HVB and psychedelic experiences certainly overlap, HVB likely induces unique experiential dimensions that are not adequately captured by existing surveys. The time-resolved free-form annotations of HVB experiences collected in this study can provide a first indication of potential HVB-specific experiential contents (see Supp. Text S3). Qualitative word-clustering of these free-form reports (see Fig. 1) points towards prominent interoception-based themes of movement, touch, tension, release, and themes of embodiment that are generally not covered by existing ASC surveys. This observation is also in line with previous work by Canello et al. (2026), who showed that body awareness was one of the main themes expressed in free-form reports of HVB experiences.

Intriguingly, the prominence of embodied sensations in HVB appears somewhat at odds with several observed neural signatures of attenuated interoception during HVB, including the Delta increase observed here and reduced cerebral blood flow in the insular cortex, a core interoceptive region (Kartar et al., 2025). Delta oscillations have been linked to impaired heart beat detection as well as dampened heart-evoked potentials, suggesting that they track a dissociation from interoceptive awareness (Haslacher et al., 2025). We propose that this apparent paradox may be resolved within a predictive-processing framework: in such a framework, markers of attenuated interoception may reflect not a loss of bodily sensation, but a loss of top-down interoceptive inference, i.e. the predictive evaluation and regulation of bodily states. Reduced precision-weighting of interoceptive priors would allow unexpected afferent signals to propagate, and thereby be experienced as novel, vivid and unconstrained; this is in line with the interoceptive prediction error hypothesis of HVB proposed by Fincham et al., (2023). This hypothesis could be tested in future by tracking heartbeat-evoked potentials, insular activity and experiential metrics of embodiment throughout HVB.

Together, these points raise the question how to capture the content of HVB (and more broadly ASC) experiences in a more nuanced and temporally precise way without disturbing ongoing experiential states. In terms of experiential content, deep-learning analyses provide exciting new avenues to extract experiential themes from free-form reports without the need for preformulated surveys, an approach that has been successfully applied in psychedelic and breathwork research (Canello et al., 2026; Cowen & Keltner, 2017; Friedman & Ballentine, 2022; Timmermann, Bauer, et al., 2023). Such approaches can also be used to build an empirical base for surveys that better capture the lived experiences of different ASCs, for instance by incorporating embodied sensations in the evaluation of HVB experiences. In terms of time-resolved tracking of experiences, the real-time rating system applied in this study as well as in Havenith et al., (2025) appears to successfully reflect ongoing ASC intensity. However, it does require repeated interruptions as well as a general understanding from participants concerning the intensity range they may encounter. Indirect behavioural inference of experience depth and content e.g. via facial expressions, posture or movement may provide a complementary and highly time-resolved approach to track experiential states via video footage, circumventing the need for explicit evaluation by the participant.

Finally, while this study focuses explicitly on the central nervous system, it stands to reason that physiological processes, including autonomic nervous system dynamics, also contribute to HVB experiences. For instance, under DMT, the level of autonomic arousal has been observed to track with affective intensity (D’Amelio et al., 2026) and sympathetic-parasympathetic coactivation has been reported as a marker of spiritual and mystical-type experience (Bonnelle et al., 2024). During HVB, heart rate increases at the beginning of the session, tracking increased experiential intensity, but gradually declines independently of ASC depth in the second half of the session, suggesting a dynamic and potentially non-linear relation between autonomic arousal and conscious state (see Supp. Fig. S3). In addition, heart rate variability has also been observed to fluctuate dynamically throughout HVB sessions, consistent with the notion that during HVB, both sympathetic and parasympathetic system exit their ordinary trajectories (Canello et al., 2026; see also Bonnelle et al., 2024).

Taken together, our findings indicate that brain states which are typically referred to as ‘psychedelic states’ do not in fact have to rely on psychedelic (or other) substances, but can arise by non-pharmacological means. In this context, HVB provides a powerful and lab-compatible benchmark for non-pharmacological access points to psychedelic-like ASCs. This suggests that similar to ASCs like dreaming or meditation, psychedelic states can be understood as an in-built processing mode of the human brain, which can be engaged via various avenues including, but not limited to, psychedelic substances. This phenomenon makes the case for an updated framework of altered consciousness, in which ASCs are studied less in terms of their induction method, and more in terms of their unique profile of physiological, neural and experiential effects; less as separate instances and more as a continuous spectrum of accessible and innate subjective experience (see (Jedlicka & Havenith, 2025; Martial et al., 2020) - in other words, a framework that maps the neurophenomenological landscape of human experience, and all the places we can go in it.

## Acknowledgements

We thank Ana Cukic, Francesca D’Amico and Tea Bogdanovic for their help with experiments, Tomás Berjaga i Buisan and Gustavo Deco for advice on data analysis, and Robin Carhart-Harris for input on earlier versions of this project. We also thank all study participants for their dedication in contributing to this study. This work was funded by the Tiny Blue Dot Foundation. LP received additional funding by PsyDAO.

## Methods

### 1. Participants

The study was approved by the ethics committee of the J.W. Goethe University Hospital Frankfurt (Germany). Participants were recruited through online platforms and breathwork communities. Inclusion criteria were: age 20–67 years; at least five previous high-ventilation breathwork (HVB) sessions; English proficiency; not pregnant; no current psychiatric diagnosis (Structured Clinical Interview for DSM; First, 2015); and a self-reported history of childhood trauma (Adverse Childhood Experiences questionnaire, Felitti et al., 1998; Childhood Trauma Questionnaire, Bernstein, D. P. & Fink, L., 1998). Of 168 candidates screened, 51 were confirmed eligible and 32 provided written informed consent and completed an in-laboratory HVB session; a subset of 9 also completed a progressive muscle relaxation (PMR) control session in randomized, counterbalanced order. Three sessions were excluded for technical or procedural reasons (playlist fault, n = 1; facilitator–participant misunderstanding of the breathing technique, n = 1; amplifier failure, n = 1), and four for excessive EEG noise (see Section 6), leaving a final sample of n = 25 (mean age 43.60 (±8.16), range 30–64; 18 female, 7 male). Full demographics are summarized in Supp. Table S1. A separate group of 8 breathwork-naive participants, recruited from the general population and who were not screened for psychiatric diagnosis or trauma history, completed a music control session.

### 2. Experimental Protocol

#### 2a. Session Structure

Each session comprised a pre-session intention-setting period followed by an 80-minute HVB session guided by a trained facilitator, with music and facilitator-administered supportive touch (See Supp. Text S1, S2 for Facilitator Protocol and Touch Protocol, respectively). EEG was recorded across seven passive resting periods and four 13-minute active breathing periods: a 4-min eyes-open rest, a 4-min eyes-closed rest, four breathing periods each followed by a 4-min eyes-closed rest, and a final 4-min eyes-open rest (Fig. 1B). Only passive resting epochs were analysed; the eyes-closed pre-session rest served as baseline.

Participants gave real-time altered-state-of-consciousness (ASC) intensity ratings at 10 timepoints (Fig. 1B), on a 1–5 scale anchored to their own previous HVB experience (1 = normal waking consciousness; 5 = near-complete loss of time/space perception, strongly reduced responsiveness, and uncontrolled emotional or bodily expression).

Questionnaires were completed via REDCap. 11 Dimensions of Altered States of Consciousness (11D-ASC; Dittrich, 1998; Studerus et al., 2010) and Mystical Experience Questionnaire (MEQ30; Barrett et al., 2015) were administered acutely post-session (within 30 minutes). Positive and Negative Affect Schedule (PANAS; Watson et al., 1988), State Trait Anxiety (STAI: Spielberger, 1983)), and Watts Connectedness Scale (WCS; Watts et al., 2022) were collected at 1 week pre- and acutely post-session, and Warwick-Edinburgh Mental Well-being Scale (WEMWBS; Tennant et al., 2007) at 1 week pre- and 1 week post-session.

Following each in lab-session, participants watched a video recording of their session and gave time-resolved spoken reports of their emotional, bodily and cognitive state. Participants paused the video and spoke freely and were additionally prompted by the experimenter at each epoch boundary. Instructions were worded to avoid leading, e.g. “describe what you were experiencing at that moment”. Facilitator was present during annotation, with participants consent. Spoken reports were audio-recorded, time-stamped against the session video, and later transcribed.

#### 2b. Music

All four active-breath epochs of the HVB session were accompanied by music that was selected to support HVB processes. For each epoch, selected music pieces were merged to avoid noticeable transitions, and towards the final minute of the epoch, music faded out gradually before a chime tone signalled the beginning of the resting-state epoch. The first two 13-minute active-breath epochs featured music with strong rhythmic components at frequencies compatible with somewhat accelerated breathing. The third epoch featured still somewhat rhythmic music with a stronger emotionally evocative (‘moving’) component. The fourth and final epoch featured more calming and meditative music with fewer rhythmic components, including a music piece featuring whale song, which was repeatedly mentioned as soothing in post-session video annotations (see Fig. 1).

#### 2c. Control conditions

PMR session followed the same structure, questionnaires and annotations, but participants were guided to tense and release muscle groups during the active periods (hands and arms, feet and legs, abdomen and back, head and neck) and breathed normally and lay still during rest, with no music or touch (See Supp. Text S4 for PMR Session Protocol). The music control had the same structure, but no facilitator or intention-setting was present; researchers described the timeline, prompted ASC ratings at epoch boundaries, and participants listened to the music during active periods.

### 3. Recording

Continuous physiological data were recorded at 2500 Hz using a BrainAmp ExG system (Brain Products GmbH, Gilching, Germany) and BrainVision Recorder. EEG was recorded from a 64-channel active electrode cap (BrainCap - Fast’n Easy, Brain Products GmbH, Gilching, Germany) referenced to Cz with ground electrode as ground. The EEG, auxiliary channels and event markers were recorded in the same file, with event markers at the start of each rest and breathing period and at each ASC rating.

ECG was recorded for removal of cardiac artefacts and for cardiac-derived respiration (CDR) signal. ECG was recorded via bipolar Ag/AgCl electrodes at two thoracic sites: a primary anterior (chest) configuration used for all our ECG-based analyses, and a posterior (back) configuration used solely for R-peak cross-validation. R peaks were detected on the filtered ECG (5–30 Hz) using a custom sliding-window peak-voting procedure, and implausible inter-beat intervals (< 0.3 s or > 3 s, or statistical outliers) were automatically excluded.

CDR traces were extracted from the primary anterior ECG signal via QRS complex modulation. Specifically, R-peak amplitudes were interpolated to form a continuous respiratory envelope, which was smoothed using a 1-second moving average. Breaths were then identified as envelope peaks (minimum separation 1.5 s). Segments in which the envelope amplitude saturated over several seconds were excluded. Breathing rate (breath per minute) was computed for each epoch as 60 divided by the mean inter-breath interval.

Video and audio footage of the participant’s face and shoulders were recorded throughout the session. All participants gave written and verbal consent to being recorded.

### 4. Word clouds

Annotations were transcribed with Whisper large-v3 (Radford et al., 2022) and manually corrected. Each transcript was segmented by timestamp into nine chronological session phases (five resting-state and four active-breath epochs). Segments that could not be assigned to a definite epoch were labelled “timeless” and excluded from phase-specific analyses. Text was tokenized and lemmatized, and filtered against standard and context-specific stopword lists (NLTK; Bird et al., 2009; see Github https://github.com/zero-noise-lab/BreathworkEEG for stopwords) to remove discursive fillers unrelated to the reported experience. Word clouds were generated per phase to descriptively visualise how reported content changed over the session.

### 5. EEG Pre-processing

EEG pre-processing was done using a customized pipeline combining functions from the Fieldtrip (Oostenveld et al., 2011) and EEGLab (Delorme & Makeig, 2004) toolboxes, implemented in MATLAB (see www.github.com/zero-noise-lab/BreathworkEEG). EEG and auxiliary channels were first downsampled from 2500 to 500 Hz and then cropped to the full experimental duration (81 min, including 5 s padding either side). Data were band-pass filtered at 1–45 Hz. Line noise at 50 Hz was removed using the pop_cleanline function in EEGLAB. Channels with excessive variability (>3 SD above the mean of the channel signal across the session) were first identified and removed using custom code implemented in MATLAB. Channels with variability of 2-3 SD above the mean of the channel signal were manually inspected, and removed if their variability was significantly higher than the majority of recording channels.

#### 5b. Independent component analysis (ICA)

Following preprocessing, an ICA was run on the pre-processed continuous data using the EEGLAB *runica* function with infomax algorithm (Delorme & Makeig, 2004). Components were then classified using the ICLabel function (Pion-Tonachini et al., 2019), which yields automated per-IC classification probabilities (eye/muscle/brain/heart/etc.) A custom matlab function then correlated the ICLabel output with AUX-derived cardiac/blink detection in order to assign each IC exactly one label — brain, eye, cardiac, muscle, or ambiguous — along with a full diagnostic record of probabilities per criterion. Based on these labels, components classed as eye- or muscle-related with a probability of 80% or above were removed automatically, components classed as brain-based with a probability of 80% or above were retained, and components of ambiguous type were smoothed manually by wavelet denoising (custom function).

The cleaned data were then segmented into the seven 4-min resting-state epochs (Fig. 1B): eyes-open resting-state baseline, eyes-closed resting-state baseline (B), four eyes-closed resting-state epochs interleaved within the HVB session (R1-R4), and a final eyes-open resting-state epoch.

Next, we identified channels that still retained excessive variability (>3 SD above the mean of the channel signal in any of the seven resting-state epochs), and removed them. All channels removed due to excessive noise levels were then interpolated using spherical splines (Perrin et al., 1989), and data were re-referenced to the average of all 63 scalp channels.

### 6. Window Selection and Signal Representations

EEG data comprised 63 scalp channels downsampled to 500 Hz. Out of the seven resting-state recordings per participant (see above), we analysed the four-minute eyes-closed pre-session baseline (B) together with four resting-state epochs interleaved throughout the HVB session (R1–R4). Each rating-state epoch was divided into 48 non-overlapping 5-second windows (2500 samples per window; 48 windows covering 4 minutes, i.e. 240 seconds in total).

Based on these windows, artifact rejection was applied to the EEG traces via a custom MATLAB function, using the following criteria: A window was rejected if any channel in it contained NaN values, an absolute amplitude >150 μV, a peak-to-peak amplitude >180 μV, or a within-epoch SD > 35 μV. From the remaining epochs, those whose mean channel variance exceeded three times the median channel variance across epochs were also rejected.

Recording sessions were included in the final analysis if at least 70% of 48 windows (= 33 windows) remained in each of the seven resting-state epochs (see above). To avoid a situation where some recorded sessions would contribute significantly more data to the analysis than others, recordings with more than 80% (= 38) of clean windows were randomly subsampled to 38 windows (fixed MATLAB randomization seed: 42). Retained recordings therefore contained 33–38 windows (165–190 s) each.

### 7. Frequency Analysis

Power spectra were estimated for each retained epoch using multitapers implemented in FieldTrip (DPSS tapers, 0.5 Hz spectral smoothing, 1–45 Hz) and averaged across 5-second windows to yield one power spectral density (PSD) per channel and epoch. Band power was computed as the area under the PSD curve (trapezoidal integration in Fieldtrip) within each frequency band.

Spectra were additionally parameterized into periodic and aperiodic components using the FOOOF algorithm (Fitting Oscillations & One-Over-F; Donoghue et al., 2020), applied in fixed aperiodic mode (no spectral knee) across 1–45 Hz, with peak widths constrained to 1–12 Hz and a peak-detection threshold of 1.5 standard deviations. This yielded, for each channel and epoch, an aperiodic offset and exponent describing the broadband background trend, and the corresponding fitted aperiodic curve. Aperiodic band power was computed as the area under this fitted curve within each band, using the same trapezoidal integration as for total band power. A residual periodic, i.e. oscillatory, spectrum was obtained by subtracting the aperiodic curve from the raw PSD in log-power space, and oscillatory band power was computed as the area under this residual spectrum within each band. Specifically, for each channel and recording, let P(f) be the multitaper PSD and A(f) the fitted aperiodic spectrum (both in linear power units). For a band b with limits [f₁, f₂], three quantities were computed by trapezoidal integration over the frequency bins within the band:

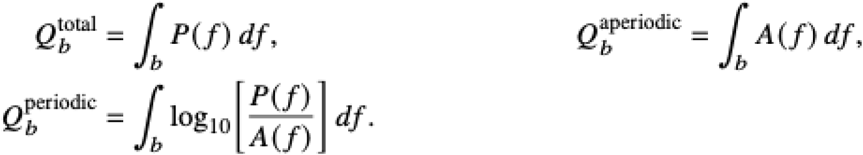

Periodic power is therefore a log-ratio spectral area, not linear total power minus aperiodic power, nor a fitted peak height. Broadband used the full frequency grid (approximately 0.9766–45.0439 Hz; nominally 1–45 Hz). Canonical bands used the stated limits: delta (1–4 Hz), theta (4–8 Hz), alpha (8–13 Hz), beta (13–30 Hz) and gamma (30–45 Hz). Global values were averaged across electrodes.

The mean model fit quality (R²) across all channels, epochs and participants was 0.967. To nevertheless account for potential contribution of remaining recording noise to the estimates of aperiodic power, we tested the relation between fit quality and estimated total, aperiodic and periodic band power. Model fit quality (R²) was unrelated to estimated band power in all bands except gamma, where it correlated negatively with total and aperiodic power, mainly due to two outlier recordings (Supp. Fig. S15).

### 8. Modulation index

To compare baseline-relative change across measures on a common scale, we computed a bounded modulation index for each measure, recording, participant and band:

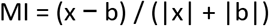

where x is the value in a post-baseline epoch and b is the value in the corresponding baseline epoch (see Seghier, 2008). As such, MI can take values from −1 to 1.

### 9. Complexity

Signal diversity was quantified with Lempel–Ziv complexity (LZc), which counts the number of distinct subsequences needed to describe a symbolic sequence (Kaspar & Schuster, 1987; Lempel & Ziv, 1976); higher values indicate a less compressible, more diverse signal. Each channel’s epoch was binarised around its own median (sᵢ = 1 if xᵢ > median(x), otherwise 0), and the binary sequence was parsed left to right with a counter c(N) incremented at each previously unseen subsequence. LZc was normalised as c(N) / [N / log₂(N)], where N is the sequence length, and computed using a compiled C implementation adapted from NeuroKit2 (Makowski et al., 2021). LZc was calculated on EEG traces (either broadband or band-pass filtered for each frequency band) per channel in each 5-second window. Estimates were first averaged across windows to obtain one LZC value per electrode, participant and epoch to be used in the topoplots (e.g. Fig. 3B). Global LZC per participant and epoch was computed as the arithmetic mean across all 63 electrodes.

### 10. Functional Connectivity

As a first step for the analysis of functional connectivity, we computed a scalp current density (CSD) map per 5-second window using spherical splines (Perrin et al., 1989) as implemented in FieldTrip (Oostenveld et al., 2011; regularisation λ = 10⁻⁵, spline order 4, maximum Legendre degree 14, conductivity 0.33 S/m). Analyses were run for broadband data and within frequency bands (see above).

Connectivity analyses used seven non-overlapping scalp regions of interest (ROIs; Supp. Table S13). The ROIs are defined as electrode groupings, not reconstructed cortical sources. Functional connectivity was then quantified with the weighted phase-lag index (wPLI; Vinck et al., 2011), which measures consistent non-zero-lag phase coupling while reducing sensitivity to spurious connectivity arising from volume conduction and common-reference effects. For electrodes i and j at frequency f, wPLI was defined as

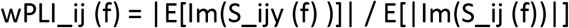

where S_ij(f) is the complex cross-spectrum between electrodes i and j, Im(·) is its imaginary part, and the expectation is taken over [time samples/tapers] within an epoch. Values range from 0 to 1 with higher values indicating more consistent non-zero-lag phase coupling.

All wPLI analyses relied on the MNE-Connectivity package implemented in Python (Binns et al., 2026). To compute an overall wPLI estimate per participant and epoch, all 5-second windows were analyzed jointly using MNE-Connectivity’s *spectral_connectivity_epochs* function with *method=’wpli’* setting and multitaper spectral estimation (Binns et al., 2026). DPSS tapers were used with a 1-Hz bandwidth (low-bias selection, non-adaptive weighting), yielding a frequency resolution of 0.2 Hz. Frequency-resolved wPLI estimates were then averaged within the respective frequency bands, while broadband wPLI was obtained by averaging all frequency bins from 1 to 45 Hz.

For each band, this yielded a symmetric 63 channel × 63 channel connectivity matrix with a zero diagonal. Global wPLI was defined as the mean across the 1,953 unique electrode pairs. Sensor-level connectivity maps were calculated for each electrode as the sum of its connections to the remaining 62 electrodes. Between-region wPLI was calculated as the mean connectivity across all electrode pairs included in two ROIs, excluding within-ROI connections. Regional analyses then summarized the 21 unique between-ROI connections.

### 11. Directed Information

The analysis of directed information flow was based on the same initial CSD analysis as the connectivity analysis described above (see Point 10). Directional interactions between the seven ROIS defined in Point 10 (see also Supp. Table S13) were quantified using a lagged conditional-information metric, termed directed information (dirINFO; Canales-Johnson et al., 2023; Ince et al., 2017; Schreiber, 2000). DirINFO is a model-free metric which compares the transfer entropy from region A to region B to the opposite direction for a range of time lags, thereby providing an estimate of both linear and nonlinear dependencies.

To compute dirINFO, CSD signals per participant and window were averaged across electrodes within each of the seven scalp ROIs and copula-normalised (Ince et al., 2017). For a source region X and target region Y at lag τ, dirINFO quantified how much information the past CSD of X provided about the present CSD of Y beyond the information already contained in Y’s own past CSD (Ince et al., 2017).

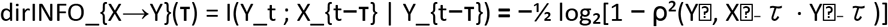

where I(·;·|·) denotes conditional mutual information and ρ is the corresponding partial correlation after Gaussian-copula transformation. Estimates were expressed in bits. Both directions were calculated independently for 40 positive time lags from 4 to 160 ms in 4-ms steps, and window-level estimates were averaged for each epoch. Zero lag was excluded because the conditional-information estimate becomes degenerate.

The primary analysis of directionality compared occipital-to-frontal and frontal-to-occipital transfer, with directional asymmetry defined as

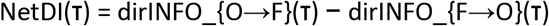

such that positive values indicate relatively stronger O→F transfer.

To quantify changes in the magnitude of directional asymmetry irrespective of direction, absolute NetDI was computed relative to baseline epochs:

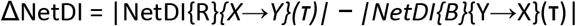

where R denotes each post-baseline resting-state epoch (R1–R4) and (B) denotes baseline. Negative values therefore indicate a decrease in directional imbalance or regional hierarchy relative to baseline.

For each participant, ΔNetDI was calculated separately for R1–R4 and then averaged to obtain an average lag curve for each ROI pair. The peak of this curve across 4–160 ms was used to summarize the strongest baseline-relative change. For Fig. 4E, this procedure was repeated across all 21 unique ROI pairs, yielding a symmetric 7 ROI × 7 ROI matrix in which each cell represents the peak change in absolute directional asymmetry between the corresponding ROIs.

### 12. Statistical Analysis

All statistical tests were two-sided (α = 0.05) with n = 25 participants. Each of the four resting-state epochs (R1–R4) was compared with the pre-session baseline (B). Multiple comparisons were controlled with the Benjamini Hochberg False Discovery Rate (BH-FDR; Benjamini & Hochberg, 1995), with the family defined for each analysis below. Effect sizes for non-parametric tests are matched-pairs rank-biserial correlations (Kerby, 2014).

#### 12a. Comparison with DMT reference data

To compare the phenomenology of HVB with DMT, 11D-ASC and MEQ30 scores were compared with published group-mean scores for intravenous DMT administration (20 mg; Timmermann et al., 2023; reference data reproduced with the authors’ permission). Because only group means were available for the reference condition, each HVB subscale was tested with a two-sided one-sample t-test of HVB scores (n = 25) against the DMT mean, treating the reference mean as a constant. Effect sizes are reported as Cohen’s D, computed as the mean difference divided by the standard deviation of the HVB scores (Cohen, 1988). To control the familywise error rate, p-values were adjusted with the Šidák correction (Šidák, 1967) separately within each questionnaire: m = 3 for the three 11D-ASC dimensions (OBN, VR, DED) and m = 4 for the four MEQ30 subscales (Transcendence, Positive Mood, Ineffability, Mystical). Adjusted p-values are reported in the Results.

#### 12b. Modulation index

Participant-level modulation indices were computed for total, aperiodic and periodic band power, global LZc, and ROI-pair wPLI. MI values were tested against zero using two-sided Wilcoxon signed-rank tests (Wilcoxon, 1945), with matched-pairs rank-biserial correlation coefficients as effect size (Kerby, 2014). For LZC, BH-FDR correction was applied across R1–R4 separately within each frequency band; for wPLI, correction was applied across the 21 ROI pairs separately within each frequency band and epoch.

#### 12c. Spatial cluster-based permutation tests

For topographic maps of each metrics and band, each epoch R1–R4 was compared to baseline for each electrode, using paired t-tests with cluster-based permutation correction (Maris & Oostenveld, 2007). Spatially adjacent electrodes exceeding a two-sided cluster-forming threshold of p < 0.05 were grouped into clusters, with summed t-values as cluster mass. Significance of this cluster mass was then assessed using 5,000 cluster permutations (randomization seed 42 in MATLAB). Probabilities for all significant clusters were then BH-FDR corrected across R1–R4 separately for each frequency band, metric, and cluster sign.

#### 12d. Directed information

To find significant sustained peaks in directed information, for each of 40 time lags (4–160 ms in 4ms steps), pairwise cross-ROI NetDI was statistically evaluated against zero using two-sided sign-flip cluster-based permutation tests (5,000 permutations, cluster-forming p < 0.05, random seed 42). Lags whose uncorrected two-sided *t*-test exceeded this threshold criterion were considered supra-threshold. Consecutive suprathreshold lags with the same sign were grouped into temporal clusters, and cluster mass was calculated as the sum of their lag-wise *t*-statistics. Each observed cluster was evaluated against the permutation distribution of the largest cluster mass. The reported map-level p-value was the p-value of the strongest observed cluster. Statistical inference therefore applies to the lag-resolved pattern as a whole, rather than to individual lags.

For frontal–occipital NetDI curves, all ten pairwise comparisons among baseline (B) and recordings R1–R4 were evaluated, with BH-FDR correction across the ten contrasts. The absolute-asymmetry analyses (Δ|NetDI|) comprised four occipital connections (anterior frontal, frontal, central, and parietal), each tested relative to baseline for each resting-state epoch. In the full seven-ROI network analysis, event-minus-baseline contrasts were evaluated for each of 21 ROI pairs across R1–R4, with BH-FDR correction across all 84 ROI-pair-by-recording tests.

#### 12e. Relationships with real-time ASC intensity

ASC ratings obtained at the start and end of each recording were analyzed separately. Associations with global power, LZC, and wPLI were assessed using within-participant correlations across B and R1–R4 (Bakdash & Marusich, 2017). BH-FDR correction was applied across 25 tests within each ASC-start and ASC-end panel (four neural metrics across six frequency bands plus one NetDI measure).

For frontal–occipital ΔNetDI, only R1–R4 were included because the baseline-relative measure is zero at baseline by definition. For each epoch, a common ΔNetDI lag was selected as the lag showing the largest absolute group-mean peak, and all participants were evaluated at that joint peak lag.

Sensor-level ASC-start associations were evaluated separately for total, periodic, and aperiodic power, LZC, and wPLI nodal strength using 5,000 within-participant permutations of ASC recording labels (seed 2143). Spatially adjacent, same-sign electrodes exceeding (p<.05) formed clusters, with BH-FDR correction applied across the six frequency bands separately within each metric and cluster sign (see Point 12c).

#### 12f. Questionnaire associations

Analyses relating survey scores to neural metrics were computed based on each participant’s epoch with the maximum ASC rating. In cases in which several epochs received the same rating, the earliest epoch was chosen. Band-specific neural metrics at this epoch were expressed relative to baseline. Frontal–occipital ΔNetDI was evaluated using the same epoch-specific peak lag described above (see Point 12e). Neural metrics were correlated separately to MEQ total, Oceanic Boundlessness (OBN), Visionary Restructuralization (VR), and Dread of Ego Dissolution (DED) using Pearson correlations. BH-FDR correction was applied across the 25 neural tests (four neural metrics across six frequency bands plus one NetDI measure) separately for each survey dimension.

#### 12g. Nonparametric tests

For each ASC dimension (11D-ASC and MEQ30 survey scores and real-time ASC ratings), normality was assessed with the Lilliefors test (p < 0.05 taken as non-normal). Where the assumption of normality is not met, we additionally tested non-parametric equivalents of the statistical tests described in Point 12e - 12f (Mann–Whitney U test for unpaired comparisons, with matched-pairs or rank-biserial r as effect size; Spearman’s rank correlations for survey–neural correlations). The Lilliefors statistics and the parametric and non-parametric results for the appropriate dimension are given in corresponding Supp. Table 20.

### 13. Software and Computation

Computations were performed in Python using NumPy, SciPy, MNE-Python, and MNE-Connectivity. Spatial inference and scalp topographies were generated in MATLAB R2024b using FieldTrip. LZC analysis was implemented in C. Participant-level inputs, permutation parameters, statistical outputs, and multiple-comparison correction definitions were retained with the analysis outputs. The code used for all computations as well as plotting can be found here: https://github.com/zero-noise-lab/BreathworkEEG

