## Supplemental Materials for "Neural and Experiential Dynamics of High-Ventilation Breathwork Mirror Psychedelic States"

**Supplemental Figures:**

Supplemental Figure S1: Correlations of mean ASC ratings and phenomenological surveys

Supplemental Figure S2: Post-session changes in affect

Supplemental Figure S3: Heart rate and real-time ASC ratings across the session

Supplemental Figure S4: Breath rate across the session

Supplemental Figure S5: Phenomenological scores of PMR and matched BW sample

Supplemental Figure S6: Phenomenological scores for music control

Supplemental Figure S7: Post-session affect and well-being for PMR

Supplemental Figure S8: Spectral EEG changes in PMR and matched BW sample

Supplemental Figure S9: Spectral Power, Complexity, and Connectivity in Music Control

Supplemental Figure S10: Complexity and connectivity in PMR and matched BW sample

Supplemental Figure S11: All hierarchical netDI relationships between area clusters in BW

Supplemental Figure S12: EEG associations with real time ASC intensity in PMR

Supplemental Figure S13: EEG associations with real time ASC intensity in music control

Supplemental Figure S14: EEG associations with ASC end and ASC delta in total breathwork sample

Supplemental Figure S15: Fit dependencies (R2) of area under the curve (AUC) powers on exponent fit

**Supplemental Tables:**

Supplemental Table S1: Demographic Information

Supplemental Table S2: 11D-ASC and MEQ30 scores in breathwork sample

Supplemental Table S3: Post-Session Changes in Affect and Wellbeing

Supplemental Table S4: Modulation Index and Sensor-Level Cluster Statistics for Power Spectra

Supplemental Table S5: 11D-ASC and MEQ30 Scores in PMR and matched HVB Sample

Supplemental Table S6: 11D-ASC and MEQ30 Scores in the Music Control compared to HVB

Supplemental Table S7: Power-Spectrum Modulation Index for PMR and Matched BW sample

Supplemental Table S8: Power-Spectrum Modulation Index for Music Control

Supplemental Table S9: LZC Modulation Indices for HVB

Supplemental Table S10: LZC in the Music Control

Supplemental Table S11: LZC in PMR and Matched HVB sample

Supplemental Table S12: Weighted Phase Lag Index for HVB

Supplemental Table S13: Electrode Assignments for Seven Scalp Regions of Interest

Supplemental Table S14: Pair-wise cross-region Weighted Phase-Lag Index for HVB

Supplemental Table S15: Weighted Phase-Lag Index in PMR

Supplemental Table S16: Weighted Phase-Lag Index in the Music Control

Supplemental Table S17: Net Directed Information for Frontal–Occipital Regions

Supplemental Table S18: Peak-Lag Net Directed Information for Frontal–Occipital Regions

Supplemental Table S19: Absolute Net Directed Information from Occipital to Four Other Regions

Supplemental Table S20: Associations Between Neural Metrics and Real-Time ASC Ratings

Supplemental Table S21: Correlations Between Peak-ASC Neural Metrics and Post-Session Survey Scores

**Supplemental Texts:**

Supplemental Text S1: Breathwork Intervention & Facilitation Protocol

Supplemental Text S2: Protocol for Facilitator Touch Interventions

Supplemental Text S3: Examples of verbal session annotations

Supplemental Text S4: PMR Intervention Protocol

__**Supplemental Figure S1: Correlations of mean ASC ratings and phenomenological surveys**__

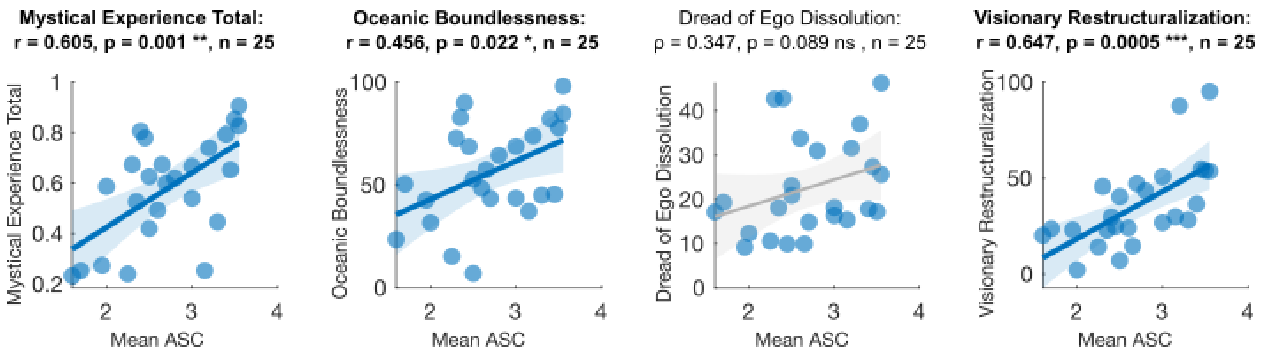

*Scatter plots and linear regressions of four post-session survey scores of subjective phenomenology predicted by mean ASC rating per participant (n = 25) across R1-R4. Mean ASC intensity was significantly associated with Mystical Experience Total (Pearson's correction; n = 25; r = 0.605, p = 0.0013), OBN (Pearson's; r = 0.456, p = 0.0220), and VR (Pearson's; r = 0.647, p = 0.0005); all three remained significant after BH-FDR correction across the four correlations. No significant association was found with DED (Pearson's r = 0.314, p = 0.1258; Spearman's ρ = 0.347, p = 0.0891; DED deviated from normality (Lilliefors tests; p = 0.032) so both Pearson's and Spearman's are reported). * = p < 0.05, ** = p < 0.01, *** = p < 0.001.*

#### Supplemental Figure S2: Post-session changes in affect

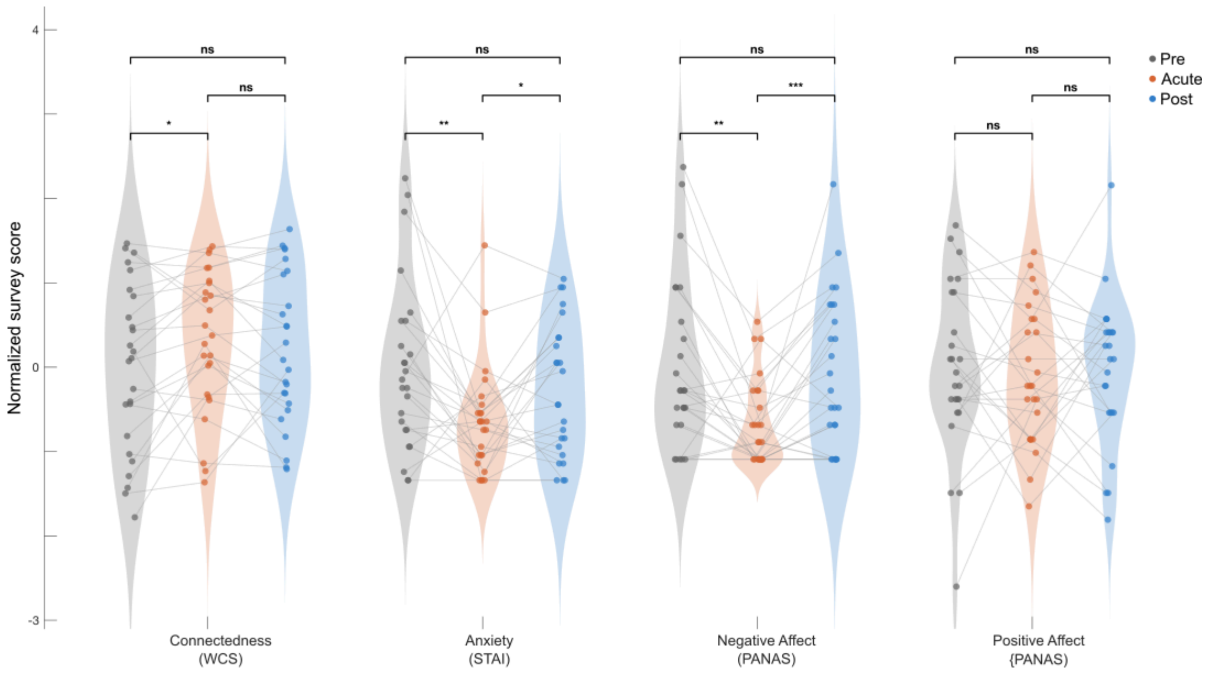

*Violin plots of survey scores from four surveys of affect (WCS, STAI, PANAS-positive and PANAS-negative). Scores were normalized for visualization normalized by z-scoring relative to average and standard deviation of pre-session scores. Grey: 1 week pre-session. Red: Acute post-session. Blue: 1 week post-session. Connected dots: participants (n = 25). * p <0.05, ** p <0.01. See Supp. Table S3 for full statistics. Statistical significance was determined on raw (non-normalized) values using paired-samples t-tests or Wilcoxon signed-rank tests, as appropriate based on normality (Lilliefors test; see main text for statistics). Connectedness increases and anxiety and negative affect decrease significantly at acute post-session measurement, but return to baseline at a one-week follow-up.*

#### Supplemental Figure S3: Heart rate and real-time ASC ratings across the session

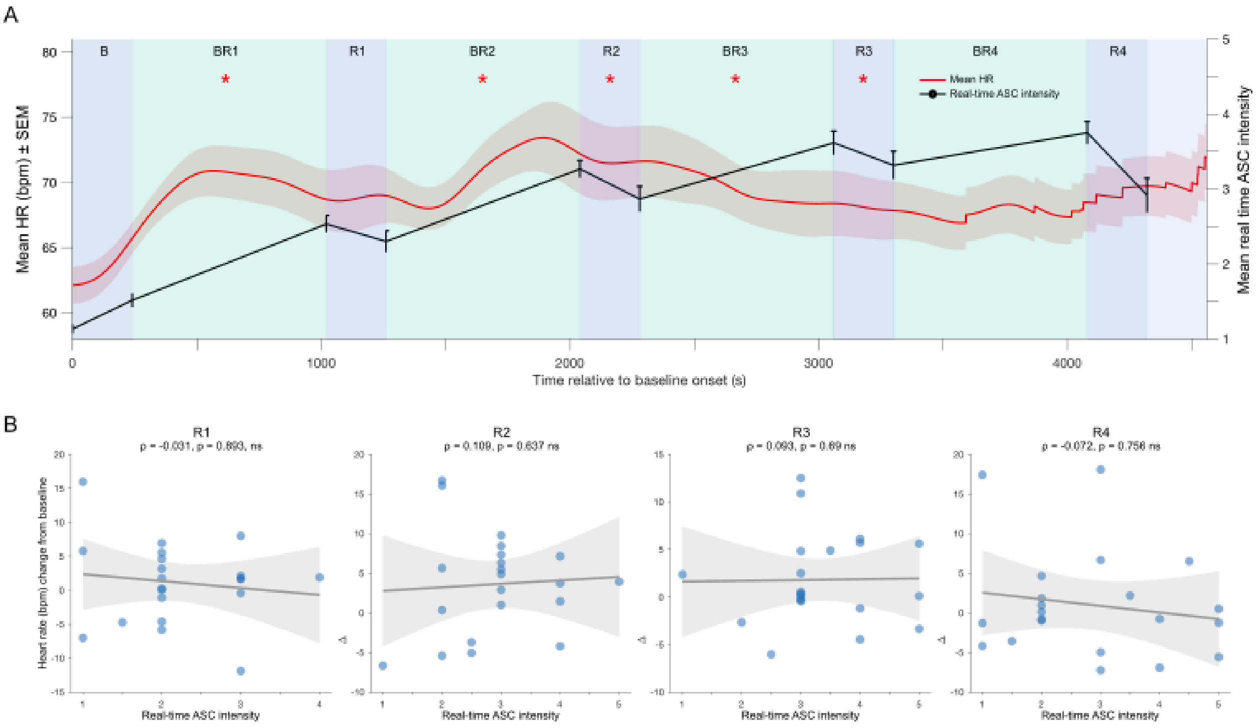

*Relation between heart rate and ASC depth ratings across the session.* ***A)*** *Time course of heart rate and ASC ratings throughout the HVB session for 25 participants. Red line: mean heart rate. Red shading: SEM. Black line: mean ASC intensity rating. Error bars: SEM. Turquoise background: active breathing epochs 1-4. Blue background: five resting-state epochs (Baseline, R1-R4). * $p < 0.05$ for comparisons of heart rate per epoch to baseline heart rate (Wilcoxon signed-rank tests; $n = 25$). Mean heart rate increases early in the session from the first breathing trial (Wilcoxon; BR1: $r = 0.73$, $p = 0.00024$; BR2: $r = 0.72$, $p = 0.00030$; R2: $r = 0.56$, $p = 0.0054$; BR3: $r = 0.75$, $p = 0.00017$; R3: $r = 0.43$, $p = 0.030$; all significant after BH-FDR). By BR4, heart rate no longer differs from baseline (Wilcoxon; $r = 0.28$, $p = 0.16$; BH-FDR).* ***B)*** *Scatter plot and linear regression between real-time ASC ratings and heart rate change from baseline per resting-state epoch R1-R4 (see A). Heart rate and ASC ratings did not correlate signficantly (Spearman's rank correlations; $n = 25$: R1: $\rho = -0.031$, $p = 0.89$; R2: $\rho = 0.109$, $p = 0.64$; R3: $\rho = 0.093$, $p = 0.69$; R4: $\rho = -0.072$, $p = 0.76$; none significant before or after BH-FDR).*

#### Supplemental Figure S4: Breath rate across the session

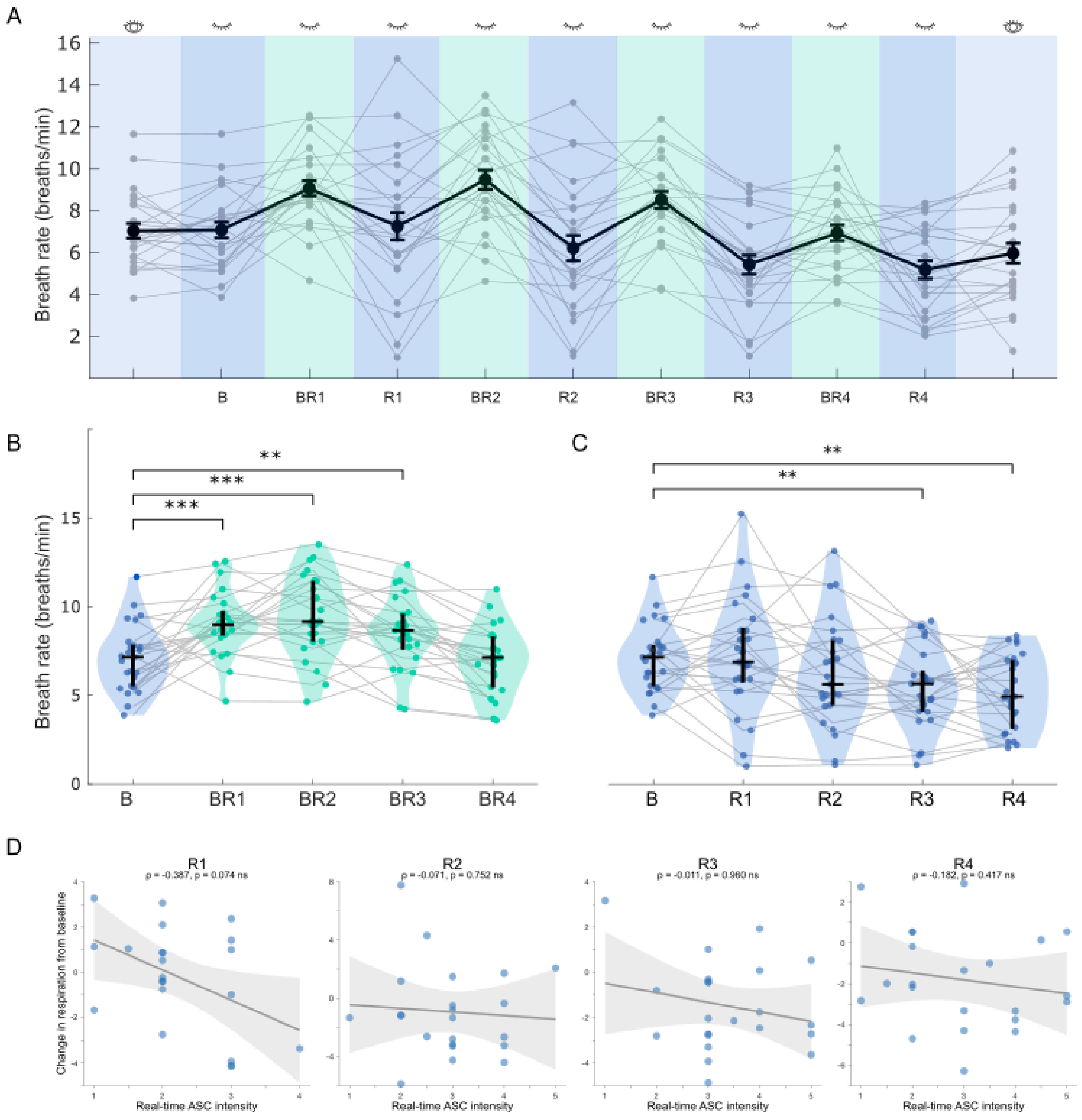

*Relation between respiratory rate and ASC depth ratings across the session.* ***A)*** *Black line: Mean respiratory rate (breaths per minute). Error bars: SEM. Grey lines: Individual respiration rates per participant (n = 25).* ***B)*** *Violin plots of breathing rate across baseline (blue) and four active breathing epochs (turquoise, BR1 - BR4). Horizontal black line: Median. Vertical black line: 25th to 75th percentile.*

*Grey lines connect values for individual participants. * p < 0.05, **p < 0.01, ***p < 0.001. Breath rate increased significantly from baseline in the first three active-breath epochs, and returned towards baseline in the final epoch (paired t-test against baseline; n = 25; df = 24; BR1: t = 4.14, d = 0.83, p = 0.00037; BR2: t = 4.30, d = 0.86, p = 0.00024; BR3: t = 3.40, d = 0.68, p = 0.0024; effects remain significant after BH-FDR).* ***C)*** *Same as B for four resting-state epochs (R1-R4). Breath rate does not change significantly from baseline in the first half of the session, but decreases in the second half (paired t-test against baseline; n = 25; df = 24; RS3: t = 4.22, d = 0.84, p < 0.001; RS4: t = 4.02, d = 0.80, p < 0.001).* ***D)*** *Scatter plot and Spearman rank correlation between ASC ratings and breath rate change from baseline across time points R1-R4. Black line: linear regression. Grey shade: 95% Confidence Interval of fitted regression line. There were no significant correlations between breath rate and ASC depth (n = 22; R1: ρ = -0.388, p = 0.075; R2: ρ = -0.071, p = 0.752; R3: ρ = -0.011, p = 0.960; R4: ρ = -0.182, p = 0.417; none significant before or after BH-FDR).*

**Supplemental Figure S5: Phenomenological scores of PMR and matched BW sample**

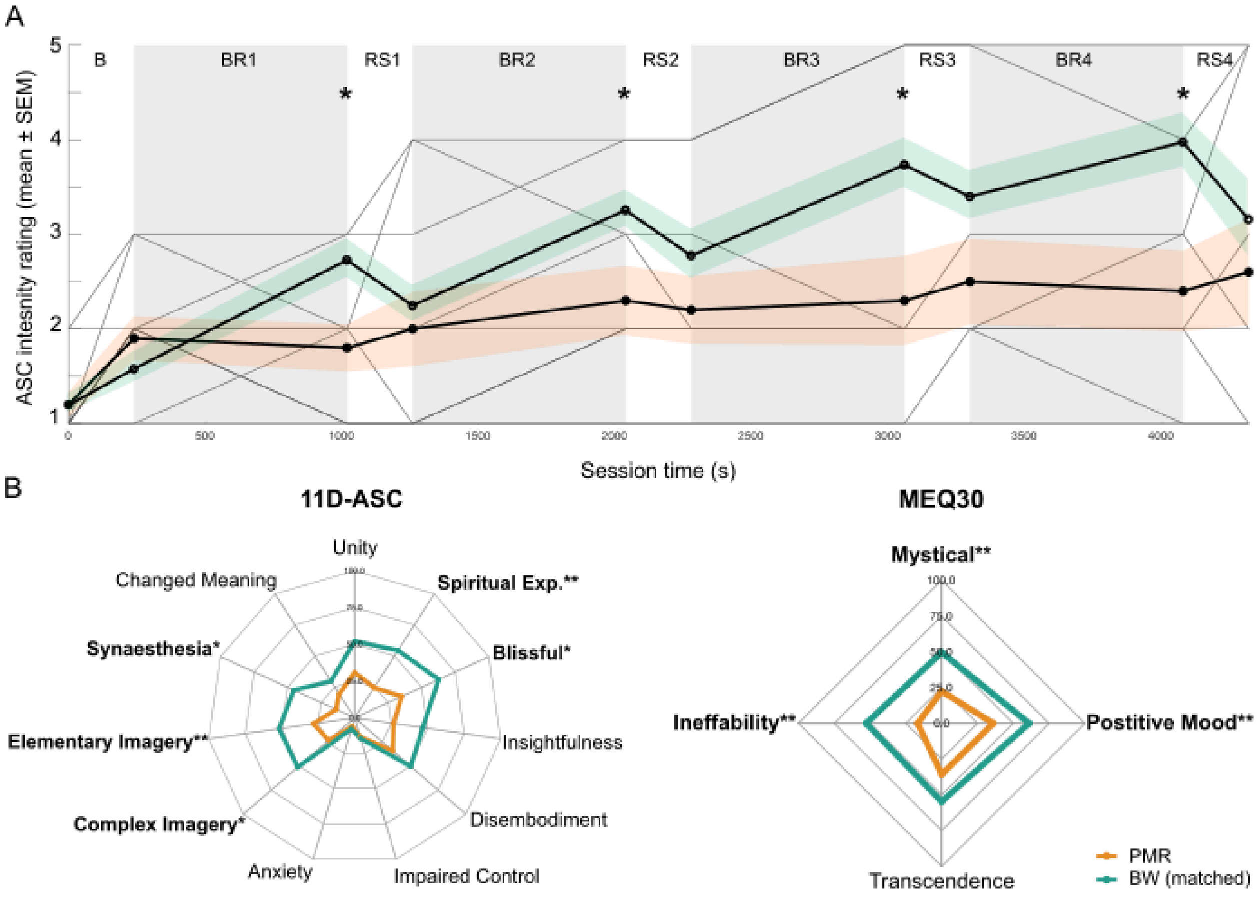

***Real time ASC intensity is lower in PMR (n = 10) than the matched breathwork sample (n = 10) after active trials, and PMR results in lower scores in some survey dimensions. A)*** *Real time ASC intensity across session timecourse in PMR (orange) and matched BW sample (green). Mean ASC intensity in PMR 2.12 (±1.04) is significantly lower than mean ASC intensity in matched BW sample 2.78 (±0.51) (paired sample t-test; n = 10; t(9) = −2.37, d = −0.75, p = 0.042). Matched breathwork sample has significantly higher intensity ratings than PMR at end of all 4 active epochs (paired samples t-test; n = 10; R1: d = 0.97. p = 0.0134; R2: d = 0.78, p = 0.0355; R3: d = 0.94, p = 0.0159; R4: d = 1.11, p = 0.0066; * = p < 0.05).* ***B)*** *11D-ASC and MEQ30 scores in PMR and matched BW sample. Significant 11D-ASC differences were observed for Spiritual Experience, Blissfulness, Complex Imagery, Elementary Imagery, and Synaesthesia. For MEQ30, significant differences were observed for Mystical, Positive Mood, and Ineffability (bold indicates p <0.05; ** = p < 0.01; see Supp. Table S11 for full statistics).*

**Supplemental Figure S6: Phenomenological scores for music control**

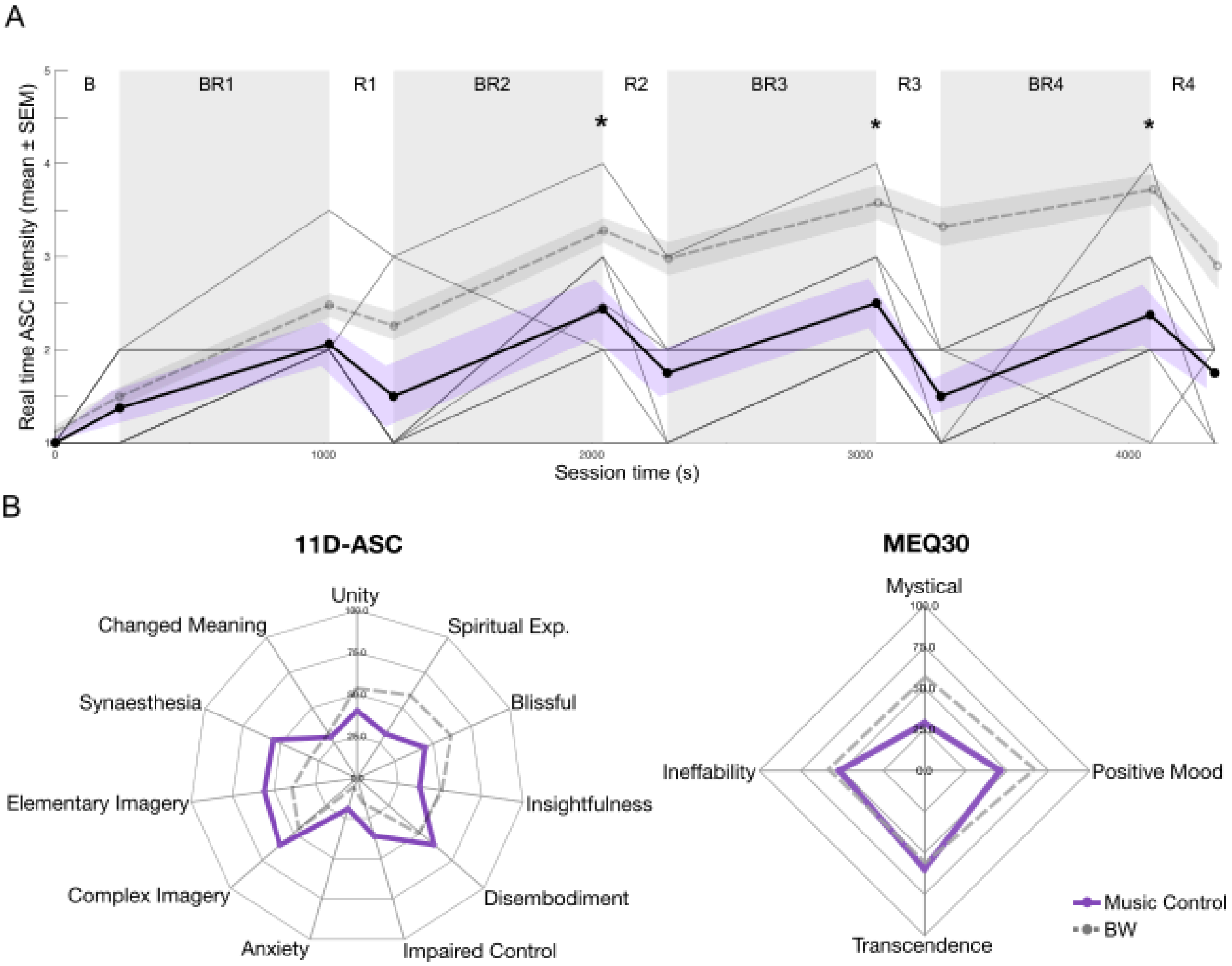

*Phenomenological and survey scores in music control (n = 8) and total breathwork sample (n = 25).* ***A)*** *Real time ASC intensity across session timecourse in music control (purple; n = 8) and total breathwork sample (dashed grey; n = 25). Mean real time ASC intensity score in music control 1.83 (±0.40) is significantly lower than mean real time intensity of 2.72 (±0.58) in the total breathwork sample (Welch's t-test, t(17.46) = −4.87, d = −1.78, p < 0.001). ASC intensity was lower in the music control at the end of BR2–BR4 (Mann–Whitney U owing to non-normal distributions, unpaired; BR2: r = −0.56, p = 0.0126; BR3: r = −0.61, p = 0.0091; BR4: r = −0.71, p = 0.0023; all remain significant after BH-FDR; * = p < 0.05).* ***B)*** *11D-ASC and MEQ30 scores in music control (purple; n = 8) and total breathwork sample (dashed grey; n = 25). In music control, mean 11D-ASC subscale scores: OBN 38.45 (±14.72), VR 50.21 (±19.51), DED*

*38.50 (±14.90). Mean MEQ30 scores: Transcendence 59.58 (±19.72), Positive Mood 45.83 (±16.01), Mystical Experience 28.83 (±24.62), Ineffability 51.67 (±25.39). No differences between music control and total breathwork sample in 11D-ASC or MEQ30 dimensions are significant after BH-FDR correction; see Supp. Table S18 for all dimensions.*

**Supplemental Figure S7: Post-session affect and well-being for PMR**

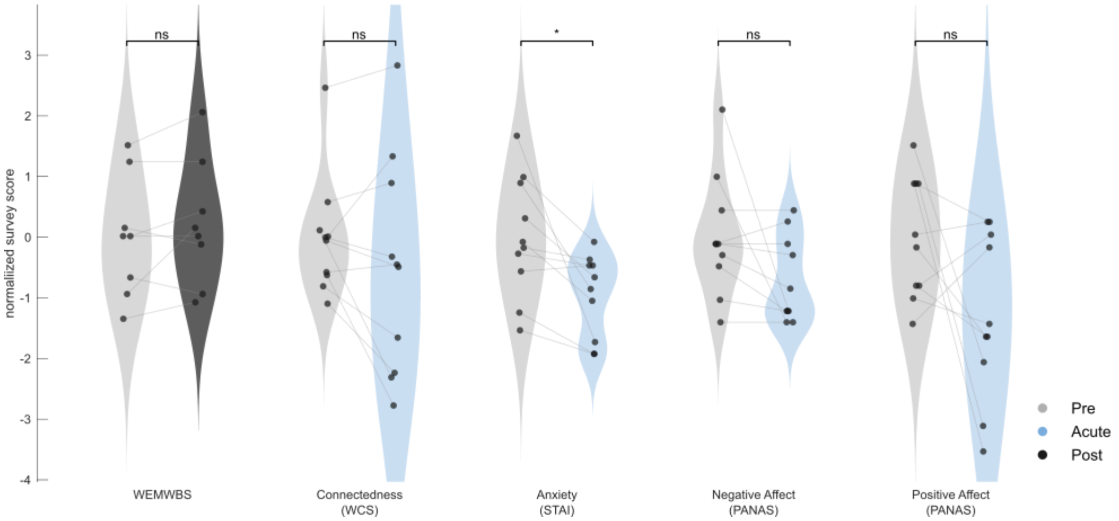

***State anxiety decreases following PMR session (n = 10), no other changes in affect acutely or one-week post PMR session.*** *One-week pre-session values (grey), with one-week post (dark grey) and acute (blue) values. State anxiety (STAI) decreased from pre to acute (39.80 (±10.29) to 30.00 (±7.01); paired samples t-test; n = 10; d = 0.94, p = 0.016; q = 0.064 after BH-FDR across the four acute comparisons). No significant pre-to-acute change was observed in connectedness (paired-samples t-tests; n = 10; WCS total; 66.31 (±10.74) to 60.75 (±19.35); d = 0.45, p = 0.192), negative affect (PANAS negative; 17.60 (±5.42) to 13.80 (±3.85); d = 0.585; p = 0.078), or positive affect (PANAS positive; 32.80 (±4.76) to 26.60 (±6.54); d = 0.64, p = 0.073). Wellbeing (WEMWBS) did not change between pre and one-week post (47.88 (±7.34) to 49.50 (±7.67); d = 0.48, p = 0.216; n = 8, as two participants did not complete the post-session timepoint). * = p< 0.05 (uncorrected).*

#### Supplemental Figure S8: Spectral EEG changes in PMR and matched BW sample

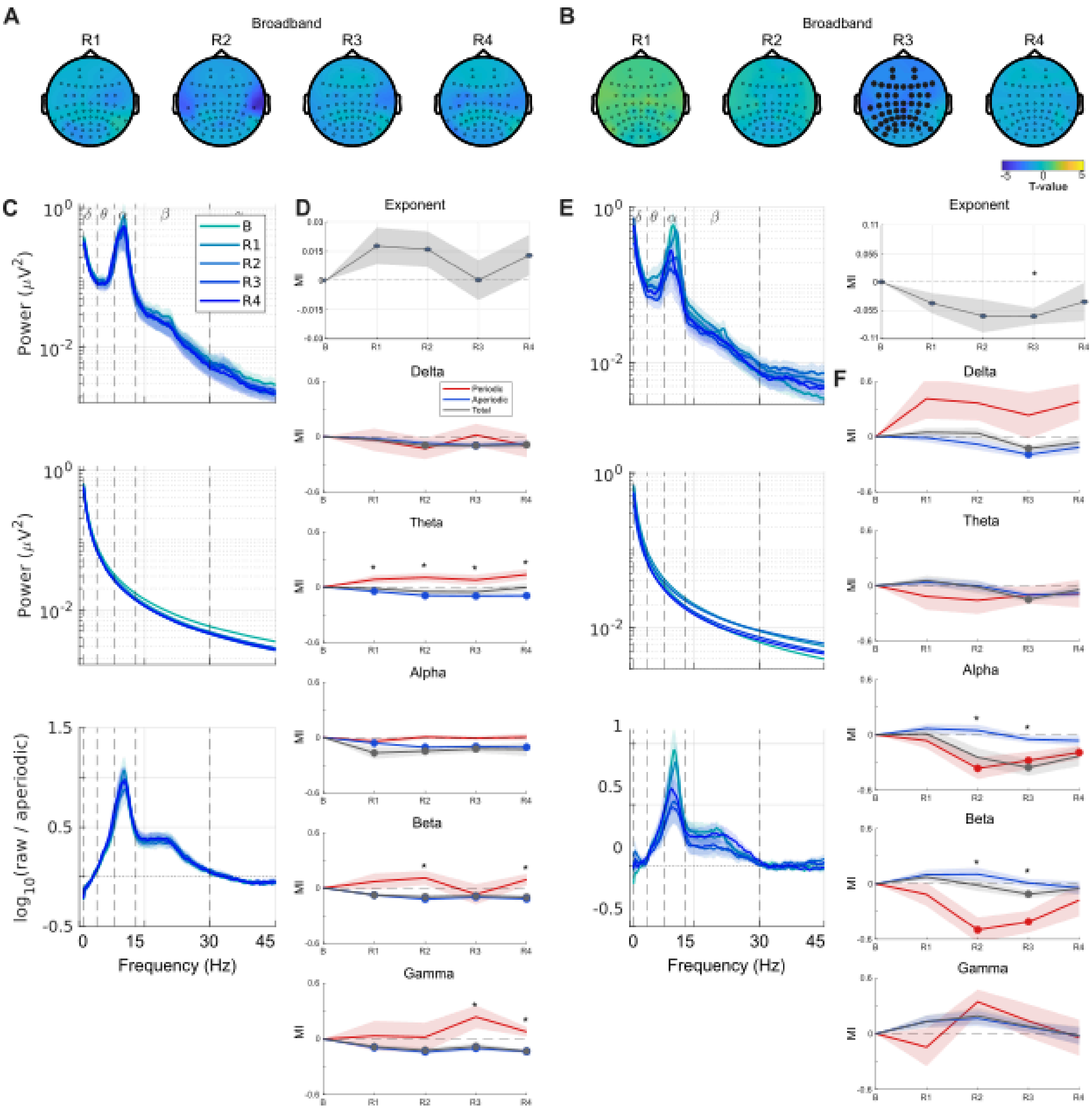

*Power spectrum changes in PMR (n = 10; left) and matched breathwork sample (n = 9; right). EEG data from one BW participant who additionally completed a PMR session was rejected on the basis of too many artefacts.* ***A)*** *Sensor t-maps show total broadband power across session timecourse; no significant clusters remain in PMR.* ***B)*** *Significant decrease in total broadband power observed at R3 in matched*

*breathwork sample (cluster-based permutation test, n = 9; p = 0.0039, significant after BH-FDR-correction).* ***C)*** *Total power spectrum density (top), aperiodic power spectrum density (middle), and periodic power spectrum density (bottom) across the session from baseline (light teal) to R4 (dark blue) in PMR.* ***D)*** *Modulation indices to baseline of aperiodic exponent, and canonical frequency bands (delta-gamma) in total (grey), aperiodic (blue), and periodic (red) power across the session timecourse in PMR (n = 10), with SEM shading. Baseline (B) is zero by definition. Effect size is the matched-pairs rank-biserial correlation (r; Kerby, 2014). Periodic vs. aperiodic direct comparison; * in figure; significance from baseline indicated by filled data points in figure. (See Supp. Table S12 for statistics).* ***E)*** *Total power spectrum density (top), aperiodic power spectrum density (middle), and periodic power spectrum density (bottom) in matched breathwork sample (n = 9). Same conventions as S5C.* ***F)*** *Modulation indices of aperiodic exponent and canonical frequency bands (delta-gamma)  total (grey), aperiodic (blue), and periodic (red) power across the session timecourse in matched breathwork sample (n = 9), with SEM shading; same conventions as S5D (See Supp. Table S13 for full statistics).*

#### Supplemental Figure S9: Spectral Power, Complexity, and Connectivity in Music Control

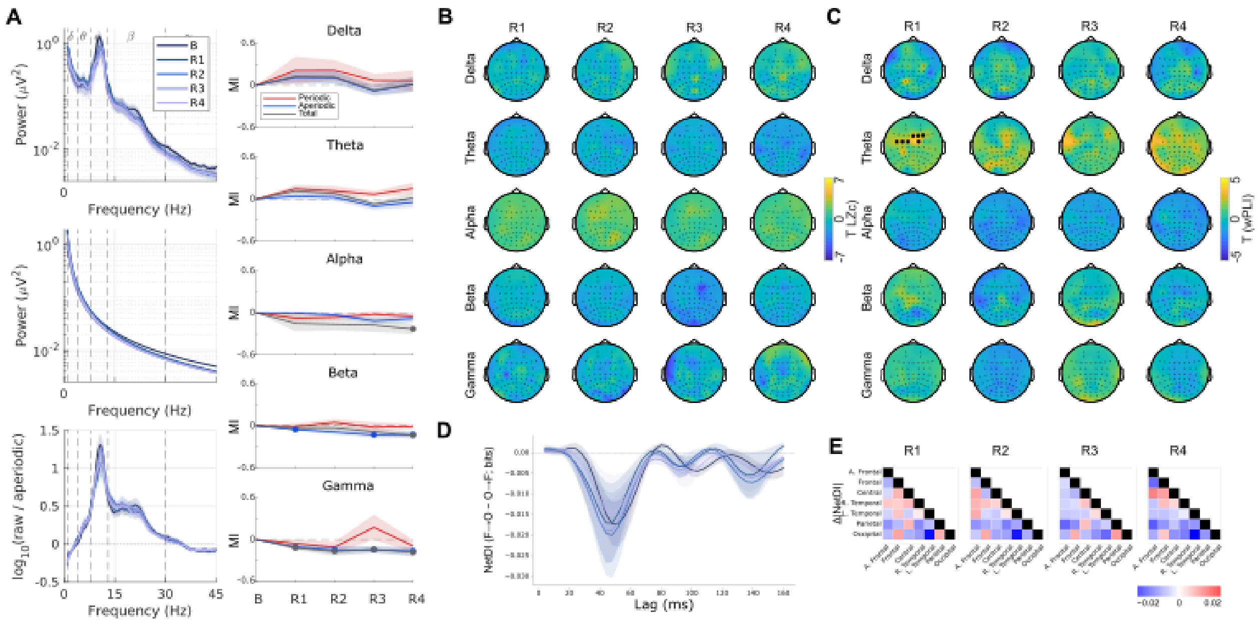

*Power spectrum, complexity, and connectivity in music control (n = 8).* ***A)*** *(Left) Power spectrum density in total power (top), aperiodic power (middle), and periodic power (bottom). Right) Modulation indices to baseline of total (grey), aperiodic (blue), and periodic (red) power across the session timecourse, with SEM shading. Baseline (B) is zero by definition. Filled symbols indicate signed-rank tests with q <0.05 (FDR-corrected) across R1–R4 within band (see Supp. Table S19 for full statistics).* ***B)*** *T(7) maps for broadband and delta–gamma voltage LZC at R1–R4 in music control (n = 8). Black markers identify electrodes in retained positive and negative spatial clusters, respectively, after two-sided cluster-permutation inference and BH-FDR across four resting maps within band and sign. Markers indicate cluster membership, not independent sensor significance. No clusters retained in LZc across the session timecourse (see Supp. Table S20 for full statistics).* ***C)*** *T(7) maps for broadband and delta–gamma voltage wPLI at R1–R4 in music control (n = 8). Same conventions at S13B. Theta connectivity (wPLI) showed a significant positive cluster at R1 versus baseline (cluster-based permutation test, p = 0.0078,BH-FDR-corrected; n = 8) but did not remain significant at R2–R4 (see Supp. Table S21 for full statistics).* ***D)*** *Mean ± SEM signed frontal–occipital netDI over 4–160 ms in music control. Positive values favor frontal-to-occipital lagged information. No clusters retained after correcting all recording comparisons.* ***E)*** *Group-selected peak Δ|NetDI| at R1–R4 in music control. Colors represent group-mean peak Δ|NetDI| magnitude (bits) across lags; red indicates stronger directional imbalance than baseline, blue indicates weaker. Matrices are symmetric; diagonals undefined. Stars denote lag-cluster tests*

*retained after BH-FDR correction across 21 ROI pairs × R1–R4. No significant lag-cluster tests were retained.*

### Supplemental Figure S10: Complexity and connectivity in PMR and matched BW sample

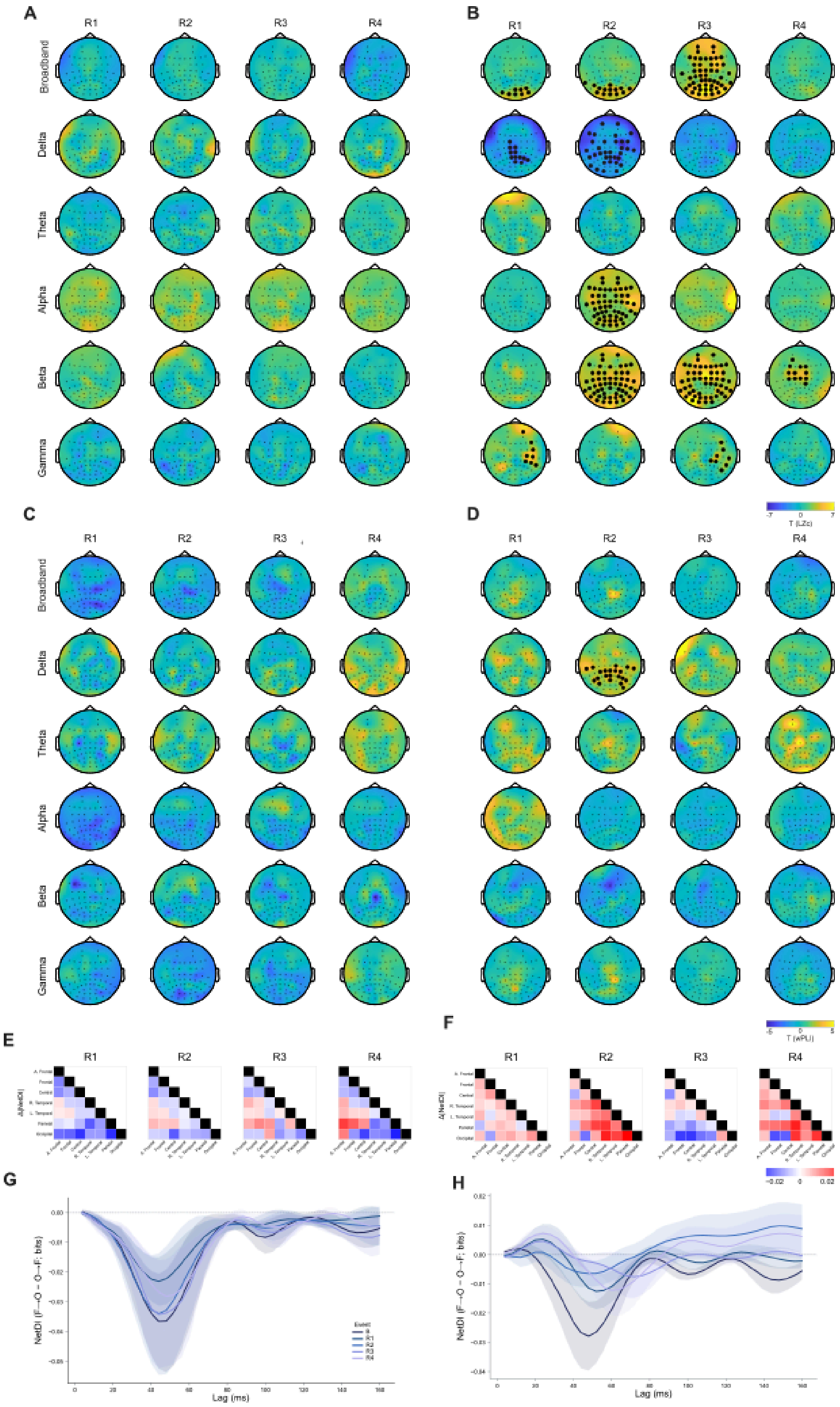

*Complexity (LZc) and connectivity (wPLI) metrics between PMR (n=10; left) and matched BW sample (n=9; right). EEG data from one BW participant who additionally completed a PMR session was rejected on the basis of too many artefacts.* ***A)*** *No changes in broadband or band-specific LZC in PMR condition (n = 10). T-sensor maps of LZc in PMR, for broadband and delta–gamma bands, at R1–R4. Paired event-minus-baseline t(9) maps. Black markers identify electrodes in retained positive and negative spatial clusters, respectively, after two-sided cluster-permutation inference and BH-FDR across four resting maps within band and sign. Markers indicate cluster membership, not independent sensor significance. No significant clusters were retained in PMR. See Supp. Table 14 for full statistics and for modulation index level statistics.* ***B)*** *T-sensor maps of LZc in matched breathwork sample (n = 9), for broadband and delta–gamma bands, at R1–R4; Same conventions as S10A except paired event-minus-baseline t(8) maps. Significant positive clusters were retained in broadband, alpha, beta, gamma, and a significant negative cluster was retained in delta (see Supp. Table S15 for full statistics and for participant level modulation index statistics).* ***C)*** *T-sensor maps of connectivity (wPLI) in PMR (n = 10), for broadband and delta–gamma bands, at R1–R4. Same conventions as S10A. No significant clusters were retained, see Supp. Table S16 for full statistics and participant level modulation index statistics.* ***D)*** *T-sensor maps of connectivity (wPLI) in matched breathwork sample (n = 9). Same conventions as S10C except paired event-minus-baseline t(8) maps. A significant positive cluster was retained in delta at R2 (see Sup. Table S17 for full statistics).* ***E)*** *Group-selected peak Δ|NetDI| at R1–R4 in PMR (n = 10), across seven regions of interest (21 pairs; see Supp. Table 6 for region definitions). Colors represent group-mean peak Δ|NetDI| magnitude (bits) across lags; red indicates stronger directional imbalance than baseline, blue indicates weaker. Matrices are symmetric; diagonals undefined. Stars denote lag-cluster tests retained after BH-FDR correction across 21 ROI pairs × R1–R4. No significant lag-cluster tests were retained.* ***F)*** *Group-selected peak Δ|NetDI| at R1–R4 in matched breathwork sample (n = 9). Same conventions as S10E. Lag curves were tested with two-sided sign-flip cluster-based permutation tests, No significant lag-curve tests were retained.* ***G)*** *Mean ± SEM signed Frontal–Occipital netDI over 4–160 ms in PMR (n = 10). Positive values favor frontal-to-occipital lagged information. No clusters retained after correcting all recording comparison (sign-flip cluster-based permutation tests, BH-FDR).* ***H)*** *Mean ± SEM signed frontal–occipital netDI over 4–160 ms in matched breathwork sample (n = 9). Same conventions as S10G. No clusters retained after correcting all recording comparisons.*

<u>**Supplemental Figure S11: All hierarchical netDI relationships between area clusters in BW**</u>

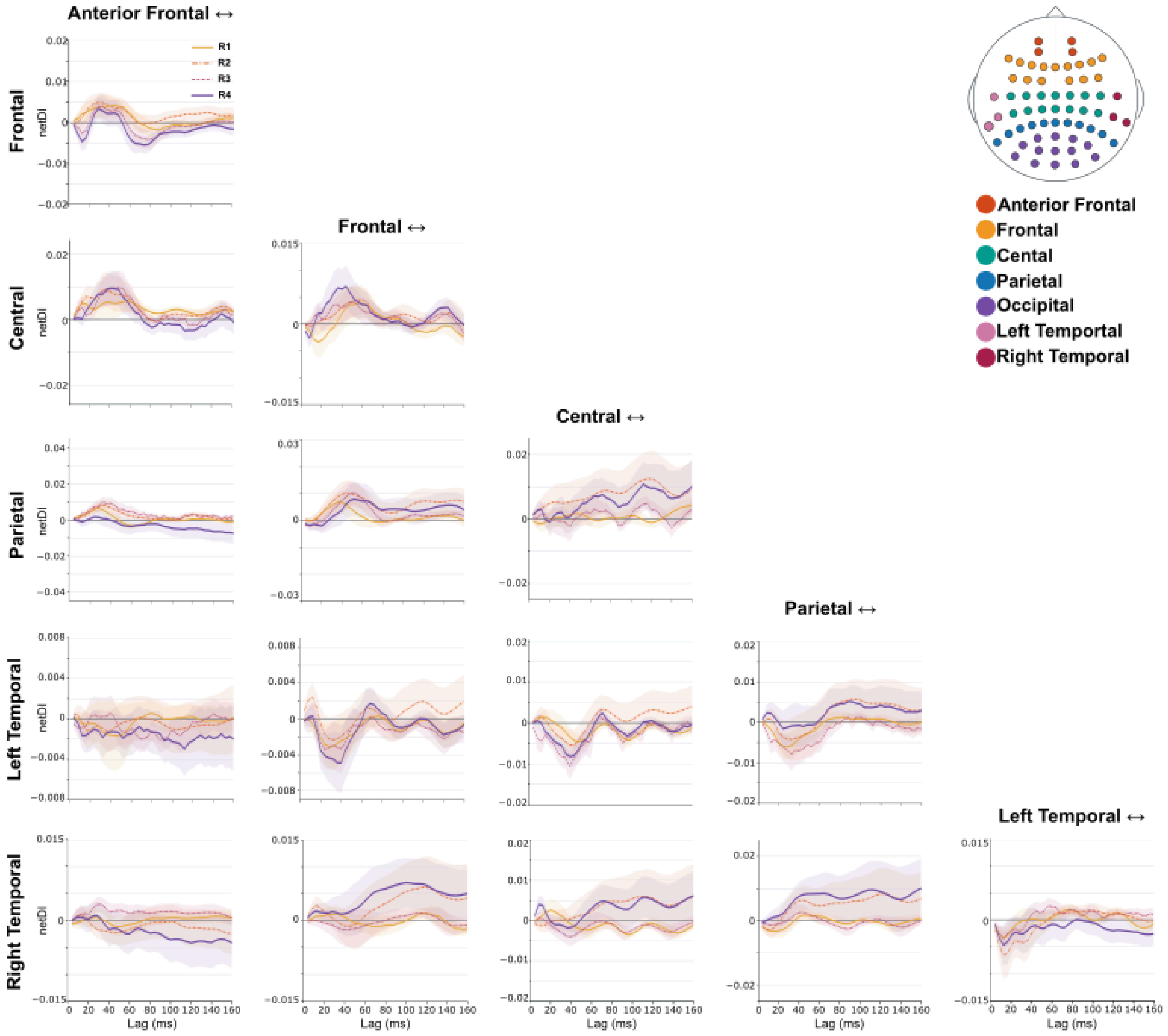

*Additional hierarchical netDI relationships between area clusters in BW (n = 25). NetDI = (Area A ⟶ Area B − Area B ⟶ Area A), where A is the area on the x-axis and B is the area on the y-axis; positive values favor A ⟶ B lagged information flow. Curves show mean ± SEM signed A ⟶ B netDI over 4–160 ms for each paired event minus ECPRE baseline. No clusters are retained after correcting across all ten recording comparisons. See main text for netDI in the occipital lobe.*

**Supplemental Figure S12: EEG associations with real time ASC intensity in PMR**

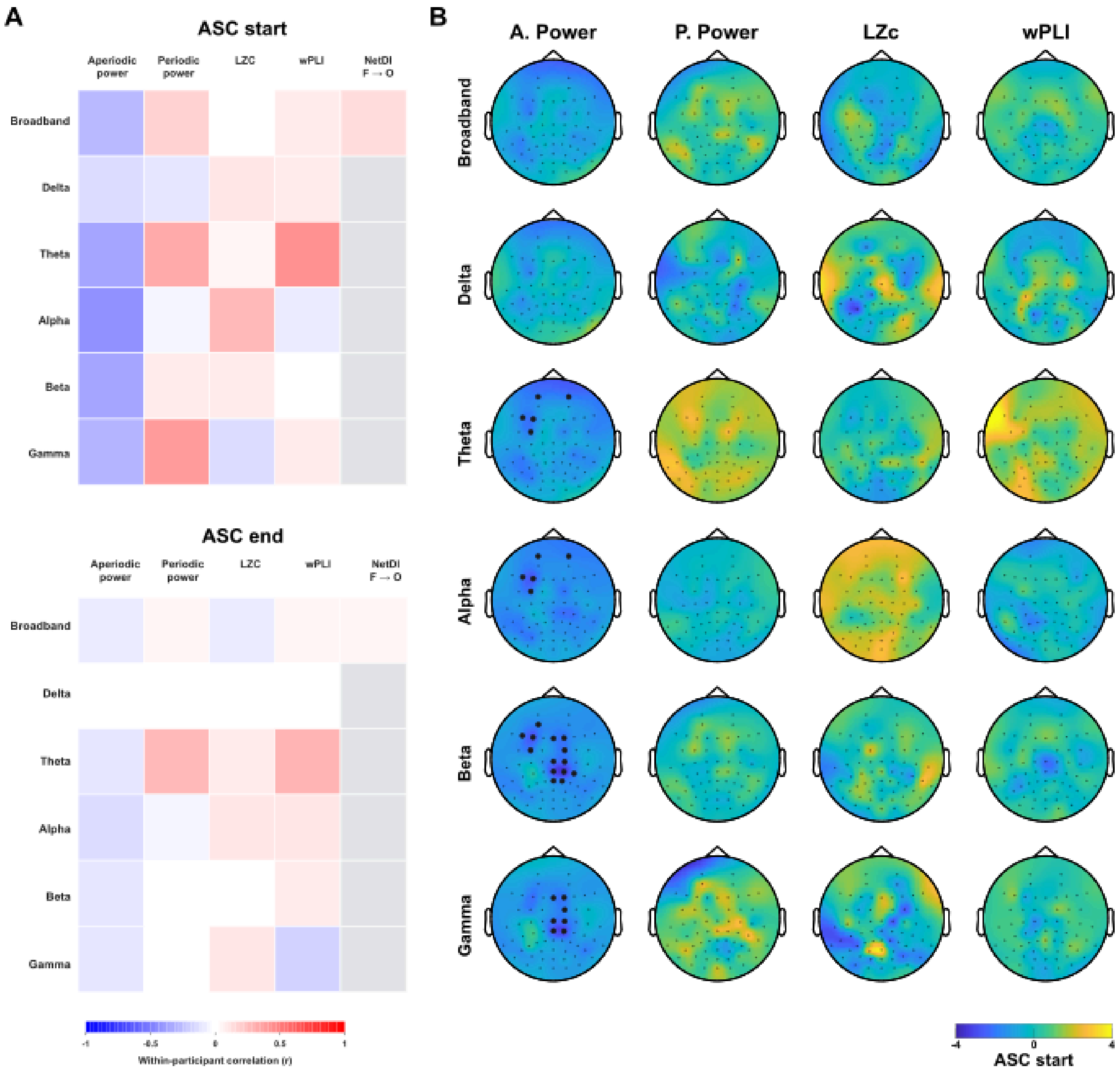

*Real time ASC intensity only correlates to aperiodic power in the theta-gamma range in PMR (n = 7). Three PMR participants were excluded from this analysis due to insufficient usable EEG data.* ***A)*** *Repeated-measures correlations (rmcorr; N orbs = 35; df = 27) between ASC start and ASC end and selected neural metrics (see figure), computed across baseline and R1-R4. No correlations survive BH-FDR.* ***B)*** *Sensor-level t-maps of the association between ASC start and total power, aperiodic power, periodic power, LZC and wPLI nodal strength in PMR. Black markers identify clusters retained after BH-FDR across six bands, separately by sign and metric. Aperiodic power showed negative clusters in theta (cluster-based permutation tests; n = 7; 5,000 permutations; participant-blocked; Σt = −11.90, 5*

*sensors, $p = 0.0098$, $q = 0.029$), alpha ($\Sigma t = -12.68$, 5 sensors, $p = 0.016$, $q = 0.030$), beta ($\Sigma t = -37.36$, 14 sensors, $p = 0.0064$, $q = 0.029$) and gamma ($\Sigma t = -19.46$, 7 sensors, $p = 0.020$, $q = 0.030$), and total power showed a negative beta cluster ($\Sigma t = -52.96$, 20 sensors, $p = 0.0048$, $q = 0.029$). No periodic-power, LZC or wPLI cluster survived correction.*

**Supplemental Figure S13: EEG associations with real time ASC intensity in music control**

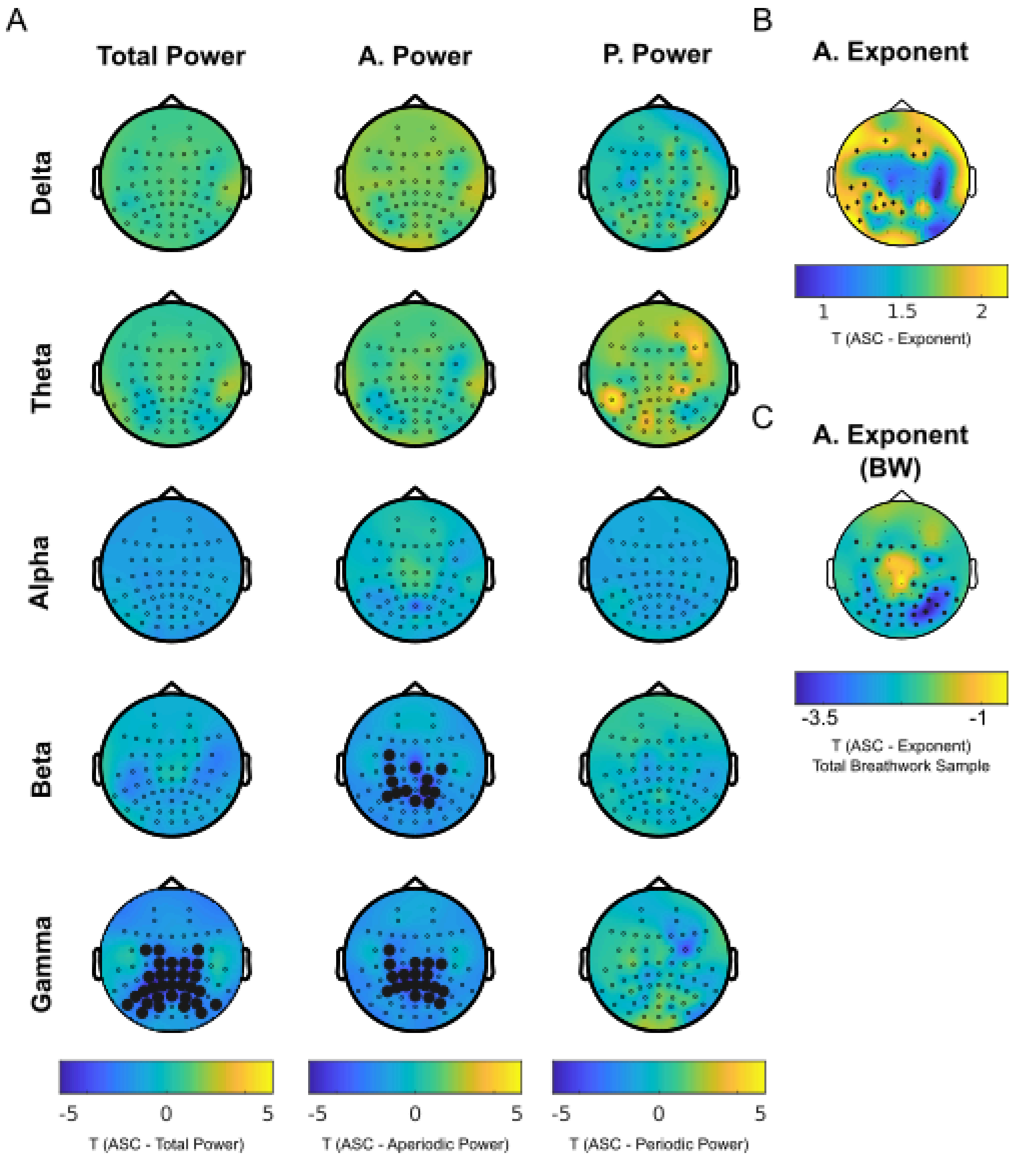

*Higher ASC intensity is associated with a steeper aperiodic exponent and lower aperiodic beta and gamma power in the music control (n = 8). Sensor-level t-maps of the within-participant association between ASC-start intensity and selected neural metrics. Black markers identify positive/negative clusters retained after participant-blocked permutations; 5,000 permutations with BH-FDR across the five bands, separately by sign and metric.* ***A)*** *Total power: negative gamma cluster (cluster permutation; n = 8; p = 0.0012, BH-FDR-significant); no other band survived correction (all p ≥0.13). Aperiodic power: negative clusters in beta (p = 0.0012) and gamma (p = 0.0008), both BH-FDR-significant; none in alpha (p = 0.35). Periodic power: no cluster survived correction (all p ≥0.093).* ***B)*** *Aperiodic exponent: one positive cluster*

*(p = 0.025; single map; uncorrected).* ***C)*** *Aperiodic exponent decrease correlates to ASC start in the total breathwork sample (within subject; n = 25; p = 0.0144).*

#### Supplemental Figure S14: EEG associations with ASC end and ASC delta

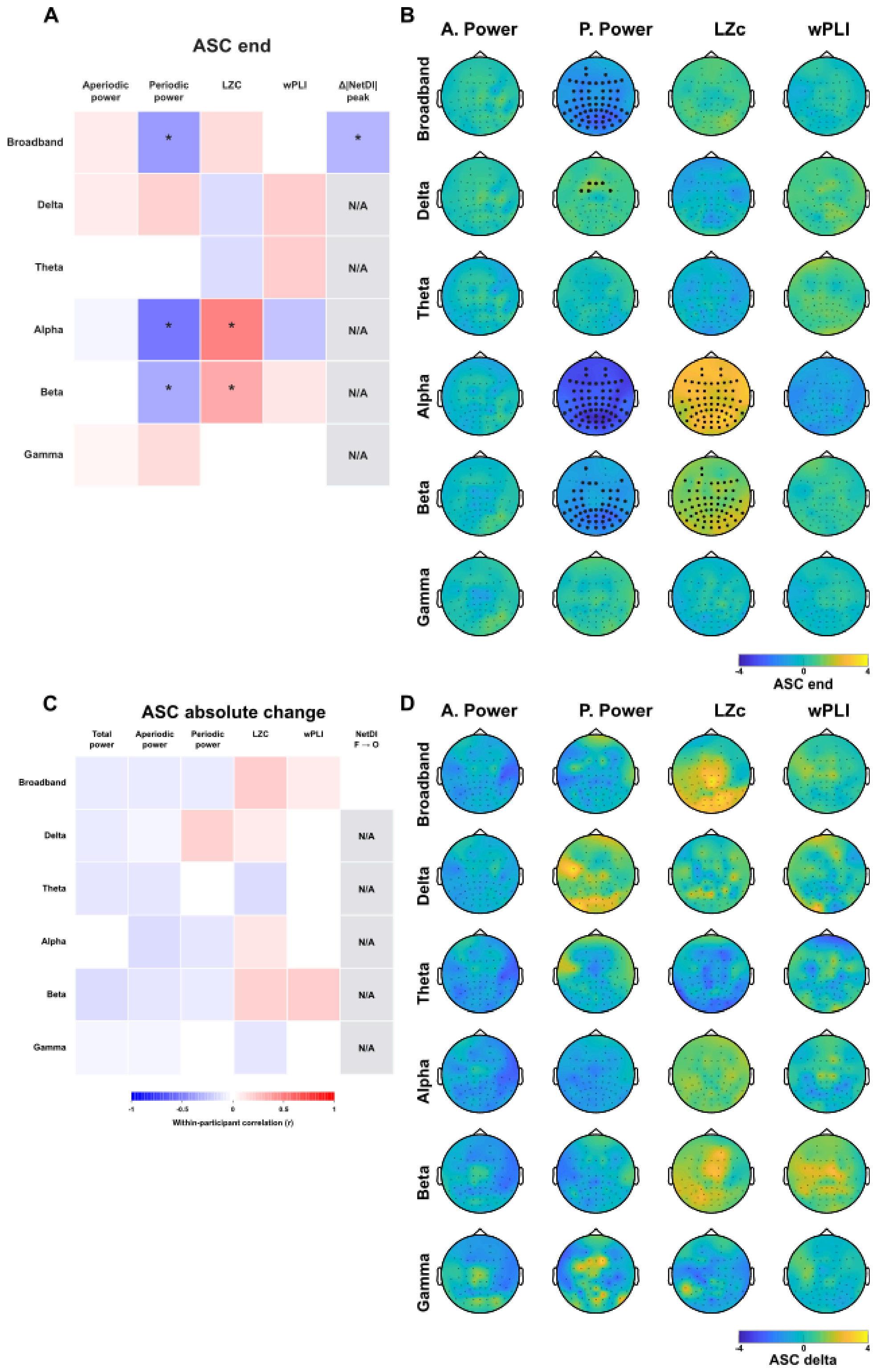

*ASC end shows a similar pattern of correlation to selected neural metrics as ASC start, however to a lower magnitude. ASC delta does not correlate to neural metrics.* ***A)*** *Repeated-measures correlations (rmcorr;*

*within-participant) between ASC end (reported ASC intensity at the end of the respective epoch) and neural metrics for the same epoch. Aperiodic power, periodic power, LZC and wPLI were computed across baseline and R1–R4 epochs (n = 25 participants × 5 time points = 125 observations, df = 99). Peak frontal–occipital Δ|NetDI| applies to epochs R1–R4, as Δ|NetDI| was computed relative to baseline (n = 25 participants × 4 time points = 100 observations, df = 74). Asterisks denote correlations significant at q < 0.05 after BH-FDR within one 25-test family per predictor. Higher ASC end was associated with lower total broadband power (rmcorr; r = −0.38, q = 0.0002), broadband periodic power (rmcorr; r = −0.39, q = 0.0002), total and periodic alpha power (rmcor; r = −0.58 and −0.53, both q < 0.001) and total and periodic beta power (rmcorr; r = −0.30, q = 0.0046; r = −0.31, q = 0.0029), and with higher alpha and beta LZC (rmcorr; r = 0.49, q < 0.001; r = 0.34, q = 0.0014) and frontal-to-occipital Δ|NetDI| (rmcorr; r = 0.40, q < 0.001)***. B)*** *Sensor-level t-statistics for the association between ASC end and aperiodic area, periodic area, LZC and wPLI nodal strength. Black markers identify significant clusters retained after participant-blocked permutation testing and BH-FDR across six bands, separately by sign and metric. Negative periodic-area clusters were observed in broadband (sensor-level cluster-based permutation test; 61 sensors, q = 0.0018), alpha (63 sensors, q = 0.0012), and beta (49 sensors, q = 0.0056), with a positive delta cluster spanning 6 sensors (q = 0.0204). Positive LZc clusters spanned all 63 alpha sensors (q = 0.0012) and 58 beta sensors (q = 0.0012). Total power also showed negative clusters in broadband (51 sensors, q = 0.0042), alpha (63 sensors, q = 0.0012), and beta (47 sensors, q = 0.0052). No aperiodic-power or wPLI cluster survived correction.* ***C)*** *No significant correlations between neural metrics and ASC delta (ASC end − ASC start; same test structure as panel S6A).* ***D)*** *No significant sensor-level clusters for ASC delta (same correction as panel as S6B).*

**Supplemental Figure S15: Fit dependencies (R2) of area under the curve (AUC) powers on exponent fit**

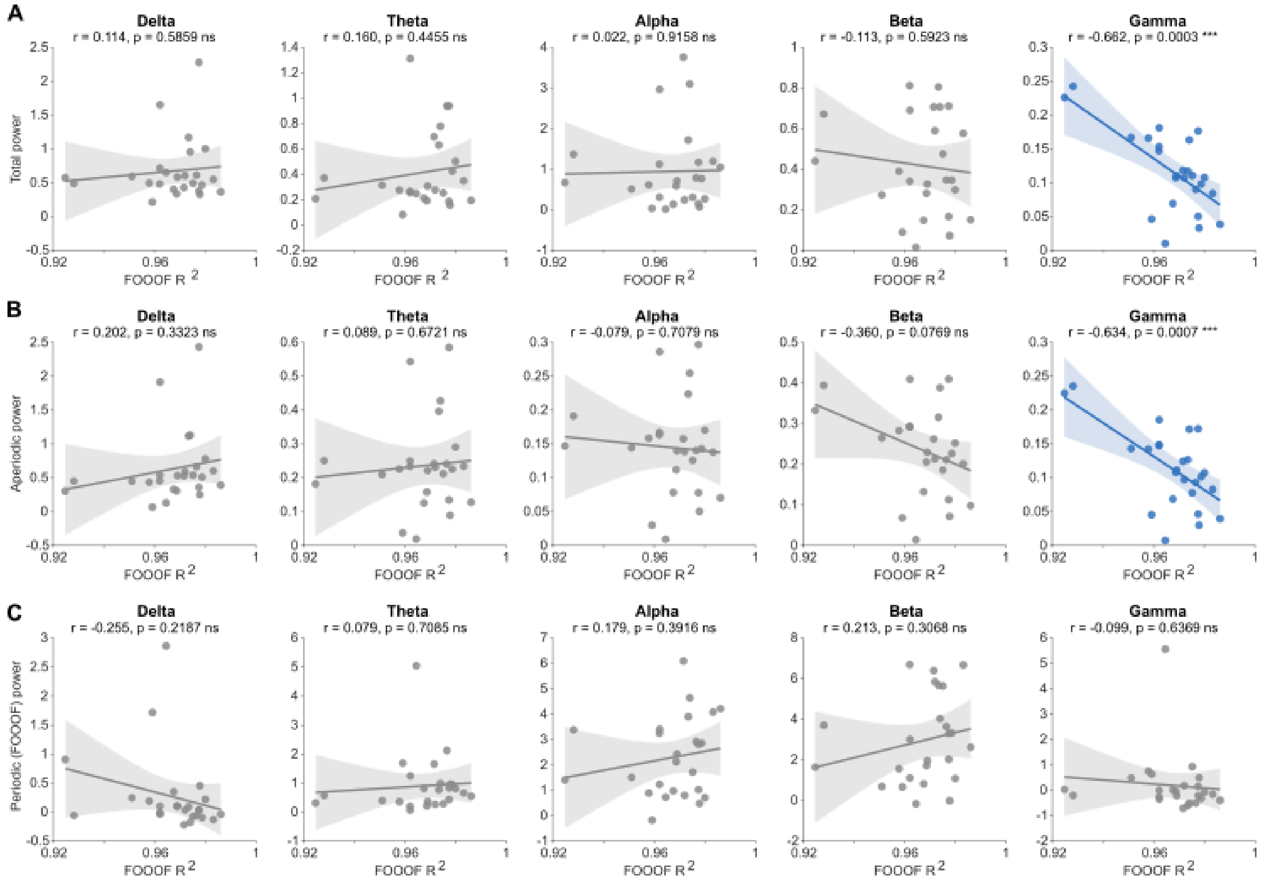

***A)** Total band power plotted against FOOOF model fit quality ($R^2$), across all five frequency bands. Blue indicates a statistically significant correlation ($p < 0.05$); grey indicates no significant relationship. Pearson's correlations; n = 25; Delta: r = 0.114, p = 0.586. Theta: r = 0.160, p = 0.445. Alpha: r = 0.022, p = 0.916. Beta: r = -0.113, p = 0.592. Gamma: r = -0.662, p <0.001.* ***B)** Aperiodic (1/f) band power plotted against FOOOF model fit quality ($R^2$), across all five frequency bands. Delta: r = 0.202, p = 0.332. Theta: r = 0.089, p = 0.672. Alpha: r = -0.079, p = 0.708. Beta: r = -0.360, p = 0.077. Gamma: r = -0.634, p <0.001.* ***C)** Periodic (FOOOF-corrected, oscillatory) band power plotted against FOOOF model fit quality ($R^2$), across all five frequency bands. Conventions as in Supplementary Figure Sx [Total]. Delta: r = -0.255, p = 0.219. Theta: r = 0.079, p = 0.708. Alpha: r = 0.179, p = 0.392. Beta: r = 0.213, p = 0.307. Gamma: r = -0.099, p = 0.637.*

**Supplemental Table S1: Demographic Information**

Total Breathwork Sample: n = 25

| Characteristic | n or value | % |
|---|---|---|
| **Age (years)** | | |
| Mean (SD) | 43.60 (8.16) | |
| Range | 30–64 | |
| **Gender** | | |
| Female | 18 | 72.0 |
| Male | 7 | 28.0 |
| **Country of residence** | | |
| Germany | 17 | 68.0 |
| United Kingdom | 4 | 16.0 |
| Netherlands | 3 | 12.0 |
| Poland | 1 | 4.0 |
| **Sessions completed** | | |
| Breathwork only | 16 | 64.0 |
| Breathwork + PMR | 9 | 36.0 |

**Supplemental Table S2: 11D-ASC and MEQ30 scores in breathwork sample**

| Dimension | M (SD) | n |
|---|---|---|
| Unity | 53.82 (26.18) | 25 |
| Spiritual Exp. | 59.13 (27.98) | 25 |
| Blissful | 61.65 (27.90) | 25 |
| Insightfulness | 50.88 (27.87) | 25 |
| Disembodiment | 48.81 (24.98) | 25 |
| Impaired Control | 14.69 (11.55) | 25 |
| Anxiety | 4.77 (7.80) | 25 |
| Complex Imagery | 44.77 (28.42) | 25 |
| Elementary Imagery | 37.97 (30.43) | 25 |
| Synaesthesia | 30.16 (28.40) | 25 |
| Changed Meaning | 31.37 (24.98) | 25 |
| Positive Mood | 64.67 (17.11) | 25 |
| Transcendence | 56.67 (24.65) | 25 |
| Ineffability | 57.60 (25.52) | 25 |
| Mystical | 55.84 (22.82) | 25 |

#### Supplemental Table S3: Post-Session Changes in Affect and Wellbeing

*All comparisons are made for the full breathwork sample (n=25). Values are reported as mean(SD); Δ = post-session minus pre-session score (mean and SD of the within-participant differences). Scores were collected 1 week before the session, acutely after the session, and at a 1-week follow-up. Normal = within-participant differences normally distributed (Lilliefors test, p ≥0.05; shown as >0.50 where the tabulated limit was reached). Paired t-tests (two-sided) are reported for all comparisons. Where differences were not normally distributed (Normal = No), the Wilcoxon signed-rank test is also reported (z, uncorrected p, matched-pairs rank-biserial r); — = not applicable (differences normal). d = Cohen's d for paired samples. p(FDR) = BH-FDR adjusted p-value for t-tests, computed separately within each panel; bold = p(FDR) <0.05. STAI = State-Trait Anxiety Inventory (state); PANAS = Positive and Negative Affect Schedule; WCS = Watts Connectedness Scale; WEMWBS = Warwick-Edinburgh Mental Wellbeing Scale (not collected acutely).*

*Panel A. Pre-session vs. acute post-session*

| **Measure** | **Pre-session** | **Acute post** | **Δ** | **Lilliefors p** | **Normal?** | **t(24)** | **p** | **p(FDR)** | ***d*** | **Wilcoxon z, p, r** |
|---|---|---|---|---|---|---|---|---|---|---|
| State anxiety (STAI) | 33.48 (10.05) | 26.88 (6.46) | −6.60 (11.07) | .302 | Yes | −2.98 | **.006** | **.013** | −0.60 | — |
| Negative affect (PANAS-negative) | 15.36 (4.91) | 12.12 (2.51) | −3.24 (4.39) | .352 | Yes | −3.69 | **.001** | **.005** | −0.74 | — |
| Positive affect (PANAS-positive) | 34.40 (6.30) | 33.76 (5.12) | −0.64 (6.30) | >0.50 | Yes | −0.51 | .616 | .616 | −0.10 | — |
| Total connectedness (WCS) | 74.93 (15.32) | 79.73 (12.79) | 4.80 (10.58) | .443 | Yes | 2.27 | **.033** | **.044** | 0.45 | — |

*Panel B. Pre-session vs. 1 week post-session*

| **Measure** | **Pre-session** | **1-week post** | **Δ** | **Lilliefors p** | **Normal?** | **t(24)** | **p** | **p(FDR)** | ***d*** | **Wilcoxon z, p, r** |
|---|---|---|---|---|---|---|---|---|---|---|
| State anxiety (STAI) | 33.48 (10.05) | 30.72 (7.97) | −2.76 (9.70) | .476 | Yes | −1.42 | .168 | .419 | −0.28 | — |
| Negative affect (PANAS-negative) | 15.36 (4.91) | 15.08 (4.45) | −0.28 (3.23) | .026 | **No** | −0.43 | .669 | .836 | −0.09 | −0.63,0.530, −.16 |
| Positive affect (PANAS-positive) | 34.40 (6.30) | 34.16 (5.47) | −0.24 (6.07) | .076 | Yes | −0.20 | .845 | .845 | −0.04 | — |
| Total connectedness (WCS) | 74.93 (15.32) | 78.11 (13.68) | 3.18 (10.06) | .123 | Yes | 1.58 | .127 | .419 | 0.32 | — |
| Wellbeing (WEMWBS) | 51.64 (6.39) | 52.12 (6.21) | 0.48 (5.28) | .220 | Yes | 0.45 | .654 | .836 | 0.09 | — |

**Supplemental Table S4: Modulation Index and Sensor-Level Cluster Statistics for Power Spectra**

*Modulation Index (MI) p and r are participant-level modulation-index statistics: matched-pairs rank-biserial correlations (Kerby, 2014) from Wilcoxon signed-rank or paired t-tests contrasting each resting epoch (R1–R4) against baseline, one 4-round BH-FDR family correction per band/metric; p is bolded where significant after BH-FDR correction, and r is signed (negative = decrease relative to baseline). Cluster p is the sensor-level statistic: the p-value of the strongest positive- or negative-mass cluster (sign shown as + or −) from a two-sided cluster-based permutation test, participant-blocked (n = 25), BH-FDR-corrected within each band/metric across the four rounds; bolded values are significant after BH-FDR correction, and an em dash (—) indicates that no cluster formed in either direction. Markers on associated topographic maps indicate cluster membership, not independent sensor-level significance.*

| Band | Metric | Round | MI p | MI r | Cluster p |
|---|---|---|---|---|---|
| **Exponent** | — | R1 | **.0119** | **−0.575** | **−0.0080** |
| | — | R2 | **.0025** | **−0.692** | **−0.0032** |
| | — | R3 | **.0173** | **−0.545** | **−0.0252** |
| | — | R4 | .0827 | −0.397 | −0.0776 |
| **Broadband** | Total | R1 | .4593 | 0.169 | +0.2436 |
| | Total | R2 | .1036 | −0.372 | **−0.0252** |
| | Total | R3 | **.0074** | **−0.612** | **−0.0252** |
| | Total | R4 | **.0007** | **−0.778** | **−0.0004** |
| | Aperiodic | R1 | .1658 | 0.317 | +0.1528 |
| | Aperiodic | R2 | .7366 | −0.077 | — |
| | Aperiodic | R3 | .2418 | −0.268 | — |
| | Aperiodic | R4 | .1094 | −0.366 | — |
| | Periodic | R1 | .2528 | −0.262 | −0.0640 |
| | Periodic | R2 | **.0023** | **−0.698** | **−0.0016** |
| | Periodic | R3 | **.0001** | **−0.877** | **−0.0004** |
| | Periodic | R4 | **.0087** | **−0.600** | **−0.0076** |
| **Delta** | Total | R1 | .0827 | 0.397 | +0.1508 |
| | Total | R2 | .5449 | 0.138 | — |
| | Total | R3 | .6377 | −0.108 | — |
| | Total | R4 | .3819 | −0.200 | — |

| | | | | | |
|---|---|---|---|---|---|
| | Aperiodic | R1 | .8401 | −0.046 | — |
| | Aperiodic | R2 | **.0230** | **−0.520** | −0.1764 |
| | Aperiodic | R3 | .0980 | −0.378 | — |
| | Aperiodic | R4 | **.0230** | **−0.520** | — |
| | Periodic | R1 | **.0012** | **0.742** | **+0.0068** |
| | Periodic | R2 | **.0019** | **0.708** | **+0.0004** |
| | Periodic | R3 | **.0019** | **0.708** | **+0.0008** |
| | Periodic | R4 | **.0005** | **0.791** | **+0.0008** |
| **Theta** | Total | R1 | .0980 | 0.378 | +0.1496 |
| | Total | R2 | .3395 | −0.218 | — |
| | Total | R3 | .0875 | −0.391 | — |
| | Total | R4 | .1036 | −0.372 | −0.3547 |
| | Aperiodic | R1 | .1218 | 0.354 | +0.0616 |
| | Aperiodic | R2 | .7366 | −0.077 | — |
| | Aperiodic | R3 | .1829 | −0.305 | — |
| | Aperiodic | R4 | **.0186** | **−0.538** | −0.1592 |
| | Periodic | R1 | .3130 | −0.231 | — |
| | Periodic | R2 | .0370 | −0.477 | −0.2072 |
| | Periodic | R3 | .1035 | −0.372 | −0.3063 |
| | Periodic | R4 | .6766 | −0.095 | +0.1684 |
| **Alpha** | Total | R1 | .2641 | −0.255 | **−0.0356** |
| | Total | R2 | **.0004** | **−0.815** | **−0.0004** |
| | Total | R3 | **<.0001** | **−0.969** | **−0.0004** |
| | Total | R4 | **.0001** | **−0.908** | **−0.0004** |
| | Aperiodic | R1 | **.0094** | **0.594** | +0.0156 |
| | Aperiodic | R2 | .2879 | 0.243 | +0.2763 |
| | Aperiodic | R3 | .9678 | −0.009 | — |
| | Aperiodic | R4 | .1353 | −0.342 | −0.4075 |
| | Periodic | R1 | .1094 | −0.366 | **−0.0192** |
| | Periodic | R2 | **.0006** | **−0.782** | **−0.0004** |
| | Periodic | R3 | **<.0001** | **−0.945** | **−0.0004** |
| | Periodic | R4 | **<.0001** | **−0.951** | **−0.0004** |
| **Beta** | Total | R1 | **.0058** | **0.631** | **+0.0096** |
| | Total | R2 | .7570 | −0.071 | −0.2843 |
| | Total | R3 | .0511 | −0.446 | **−0.0212** |
| | Total | R4 | **.0119** | **−0.575** | **−0.0056** |
| | Aperiodic | R1 | **.0021** | **0.705** | **+0.0016** |

| | | | | | |
|---|---|---|---|---|---|
| | Aperiodic | R2 | .0396 | 0.471 | +0.0556 |
| | Aperiodic | R3 | .0827 | 0.397 | +0.0860 |
| | Aperiodic | R4 | .6964 | −0.089 | — |
| | Periodic | R1 | **.0347** | **−0.483** | −0.1012 |
| | Periodic | R2 | **.0004** | **−0.809** | **−0.0016** |
| | Periodic | R3 | **.0001** | **−0.883** | **−0.0004** |
| | Periodic | R4 | **.0045** | **−0.649** | **−0.0096** |
| **Gamma** | Total | R1 | **.0017** | **0.717** | **+0.0056** |
| | Total | R2 | **.0068** | **0.618** | **+0.0076** |
| | Total | R3 | **.0214** | **0.526** | **+0.0332** |
| | Total | R4 | .5998 | 0.120 | — |
| | Aperiodic | R1 | **.0014** | **0.729** | **+0.0024** |
| | Aperiodic | R2 | **.0094** | **0.594** | **+0.0164** |
| | Aperiodic | R3 | **.0264** | **0.508** | +0.0444 |
| | Aperiodic | R4 | .8824 | 0.034 | — |
| | Periodic | R1 | .2010 | 0.292 | +0.2747 |
| | Periodic | R2 | **.0019** | **0.711** | **+0.0024** |
| | Periodic | R3 | **.0062** | **0.625** | **+0.0056** |
| | Periodic | R4 | **.0245** | **0.514** | **+0.0032** |

#### Supplemental Table S5: 11D-ASC and MEQ30 Scores in PMR and matched HVB Sample

*Bold rows indicate p <0.05. p-values from paired samples t-tests, except [U] Wilcoxon signed-rank test. n = 8 for Complex Imagery, Elementary Imagery, and Synaesthesia in the PMR sample; two participants did not complete the Visionary Restructuralization (Complex Imagery, Elementary Imagery, Synaesthesia) section of the survey; only paired comparisons made in this case.*

| | PMR | | Matched Breathwork | | | |
|---|---|---|---|---|---|---|
| **Dimension** | **M (SD)** | **n** | **M (SD)** | **n** | ***p*** | ***d*** |
| Unity | 30.86 (30.08) | 10 | 52.06 (16.54) | 10 | .067 | 0.658 |
| **Spiritual Exp.** | **24.43 (24.64)** | **10** | **54.47 (21.48)** | **10** | **.0068** | **1.106** |
| **Blissful** | **35.07 (26.37)** | **10** | **62.67 (22.79)** | **10** | **.0287** | **0.822** |
| Insightfulness | 26.73 (18.96) | 10 | 46.77 (24.92) | 10 | .121 | 0.542 |
| Disembodiment | 34.23 (23.93) | 10 | 50.37 (22.49) | 10 | .209 | 0.428 |
| Impaired Control | 13.41 (12.16) | 10 | 14.57 (8.25) | 10 | .759 | 0.1 |
| Anxiety | 6.12 (5.39) | 10 | 7.87 (8.91) | 10 | .4870 | 0.229 |
| **Complex Imagery** | **23.29 (21.16)** | **8** | **56.86 (23.29)** | **10** | **.0066** | **1.349** |
| **Elementary Imagery** | **28.17 (30.56)** | **8** | **52.83 (33.015)** | **10** | **.0024** | **1.642** |
| **Synaesthesia** | **13.83 (26.05)** | **8** | **44.79 (30.87)** | **10** | **.047[U]** | **0.922** |
| Changed Meaning | 19.27 (24.32) | 10 | 29.60 (19.97) | 10 | .0763 | 0.633 |
| ***MEQ Positive Mood*** | **36.00 (16.54)** | **10** | **61.00 (14.32)** | **10** | **.0026** | **1.302** |
| *MEQ Transcendence* | 35.67 (27.36) | 10 | 54.67 (24.50) | 10 | .0521 | 0.708 |
| ***MEQ Ineffability*** | **16.67 (16.10)** | **10** | **52.67 (22.32)** | **10** | **<0.001** | **1.959** |
| ***MEQ Mystical*** | **22.40 (19.13)** | **10** | **49.60 (18.17)** | **10** | **<0.001** | **1.631** |

#### Supplemental Table S6: 11D-ASC and MEQ30 Scores in the Music Control compared to HVB

*Welch's t-test independent sample un-paired t-test is shown for all dimensions (effect size: Cohen's d, mean difference / pooled SD). † Either group deviated from normality (Lilliefors test, p < 0.05), so the Mann–Whitney U test (effect size: rank-biserial correlation r) is also shown; — = not applicable (both groups normally distributed). Negative effect sizes indicate lower scores in the music control. Bold = uncorrected p < 0.05. q = BH-FDR-adjusted p across the 15 dimensions, computed separately for each test; no dimension was significant at q < 0.05 for either test.*

| | | | Welch's t-test | | | | Mann–Whitney U test | | | |
|---|---|---|---|---|---|---|---|---|---|---|
| **Dimension** | **Music M (SD)** | **Total BW M (SD)** | **t** | **d** | **p** | **q** | **U** | **r** | **p** | **q** |
| Unity | 40.30 (20.27) | 53.82 (26.18) | −1.52 | −0.58 | 0.148 | 0.222 | — | — | — | — |
| Spiritual Exp. | 31.38 (21.58) | 59.13 (27.98) | −2.93 | −1.11 | **0.010** | 0.070 | — | — | — | — |
| Blissful | 44.42 (21.17) | 61.65 (27.90) | −1.85 | −0.70 | 0.084 | 0.162 | — | — | — | — |
| Insightfulness | 37.71 (14.71) | 50.88 (27.87) | −1.73 | −0.59 | 0.097 | 0.162 | — | — | — | — |
| Disembodiment† | 60.63 (19.39) | 48.81 (24.98) | 1.39 | 0.53 | 0.184 | 0.230 | 131.5 | 0.32 | 0.193 | 0.263 |
| Impaired Control | 35.82 (23.13) | 14.69 (11.55) | 2.49 | 1.16 | **0.037** | 0.093 | — | — | — | — |
| Anxiety† | 19.04 (11.79) | 4.77 (7.80) | 3.21 | 1.43 | **0.011** | 0.070 | 164 | 0.64 | **0.0076** | 0.061 |
| Complex Imagery† | 60.88 (19.66) | 44.77 (28.42) | 1.79 | 0.66 | 0.090 | 0.162 | 136 | 0.36 | 0.136 | 0.245 |
| Elementary Imagery† | 55.92 (29.66) | 37.97 (30.43) | 1.48 | 0.60 | 0.164 | 0.224 | 136.5 | 0.36 | 0.130 | 0.245 |
| Synaesthesia† | 55.00 (24.73) | 30.16 (28.40) | 2.38 | 0.93 | **0.033** | 0.093 | 154.5 | 0.55 | **0.023** | 0.061 |
| Changed Meaning | 29.04 (22.21) | 31.37 (24.98) | −0.25 | −0.10 | 0.806 | 0.806 | — | — | — | — |

| | | | | | | | | | | |
|---|---|---|---|---|---|---|---|---|---|---|
| MEQ Positive Mood | 45.83 (16.01) | 64.67 (17.11) | −2.85 | −1.14 | **0.014** | 0.070 | — | — | — | — |
| MEQ Transcendence | 59.58 (19.71) | 56.67 (24.65) | 0.34 | 0.13 | 0.737 | 0.790 | — | — | — | — |
| MEQ Ineffability | 51.67 (25.39) | 57.60 (25.52) | −0.57 | −0.23 | 0.576 | 0.665 | — | — | — | — |
| MEQ Mystical | 28.83 (24.62) | 55.84 (22.82) | −2.75 | −1.14 | **0.019** | 0.070 | — | — | — | — |

#### Supplemental Table S7: Power-Spectrum Modulation Index for PMR and Matched BW Sample

*Effect size r = matched-pairs rank-biserial correlation (Kerby, 2014). Bolded values are significant after BH-FDR correction (q <0.05), applied within group/band across the four within-band tests per comparison type. PMR n = 10.*

##### Panel A: PMR

| Band | Round | Total vs. Baseline | | Aperiodic vs. Baseline | | Periodic vs. Baseline | | Aperiodic vs. Periodic (Direct) | |
|---|---|---|---|---|---|---|---|---|---|
| | | r | p | r | p | r | p | r | p |
| Delta | R1 | -0.345 | 0.3750 | -0.273 | 0.4922 | 0.091 | 0.8457 | 0.164 | 0.6953 |
| | R2 | **-0.782** | **0.0273** | -0.673 | 0.0645 | -0.382 | 0.3223 | -0.127 | 0.7695 |
| | R3 | **-0.855** | **0.0137** | **-0.891** | **0.0098** | 0.491 | 0.1934 | 0.636 | 0.0840 |
| | R4 | **-0.745** | **0.0371** | -0.600 | 0.1055 | -0.164 | 0.6953 | 0.236 | 0.5566 |
| Theta | R1 | -0.309 | 0.4316 | **-0.709** | **0.0488** | 0.745 | 0.0371 | **0.745** | **0.0371** |
| | R2 | -0.636 | 0.0840 | **-0.964** | **0.0039** | 0.600 | 0.1055 | **0.964** | **0.0039** |
| | R3 | -0.527 | 0.1602 | **-1.000** | **0.0020** | 0.382 | 0.3223 | **0.927** | **0.0059** |
| | R4 | -0.164 | 0.6953 | **-1.000** | **0.0020** | 0.600 | 0.1055 | **0.855** | **0.0137** |
| Alpha | R1 | **-0.782** | **0.0273** | **-0.964** | **0.0039** | -0.236 | 0.5566 | 0.236 | 0.5566 |
| | R2 | **-0.855** | **0.0137** | **-1.000** | **0.0020** | 0.164 | 0.6953 | 0.818 | 0.0195 |
| | R3 | **-0.818** | **0.0195** | **-1.000** | **0.0020** | -0.091 | 0.8457 | 0.745 | 0.0371 |
| | R4 | -0.673 | 0.0645 | **-1.000** | **0.0020** | 0.055 | 0.9219 | 0.673 | 0.0645 |
| Beta | R1 | **-1.000** | **0.0020** | **-1.000** | **0.0020** | 0.455 | 0.2324 | 0.673 | 0.0645 |
| | R2 | **-1.000** | **0.0020** | **-1.000** | **0.0020** | 0.491 | 0.1934 | **0.927** | **0.0059** |
| | R3 | **-1.000** | **0.0020** | **-0.964** | **0.0039** | 0.055 | 0.9219 | 0.600 | 0.1055 |
| | R4 | **-0.964** | **0.0039** | **-1.000** | **0.0020** | 0.382 | 0.3223 | **0.964** | **0.0039** |
| Gamma | R1 | **-0.855** | **0.0137** | **-1.000** | **0.0020** | 0.145 | 0.7188 | 0.418 | 0.2754 |
| | R2 | **-0.927** | **0.0059** | **-1.000** | **0.0020** | 0.109 | 0.7949 | 0.673 | 0.0645 |

| | | | | | | | | | |
|---|---|---|---|---|---|---|---|---|---|
| | R3 | **-0.709** | **0.0488** | **-0.927** | **0.0059** | 0.527 | 0.1602 | **1.000** | **0.0020** |
| | R4 | **-1.000** | **0.0020** | **-1.000** | **0.0020** | 0.491 | 0.1934 | **0.964** | **0.0039** |

**Panel B: Matched BW Sample**

*Same conventions as Table S7a. PMR n = 10. Matched breathwork sample n = 9 (one participant excluded due to noisy EEG data).*

| Band | Round | Total vs. Baseline | | Aperiodic vs. Baseline | | Periodic vs. Baseline | | Aperiodic vs. Periodic (Direct) | |
|---|---|---|---|---|---|---|---|---|---|
| | | r | p | r | p | r | p | r | p |
| Delta | R1 | 0.422 | 0.3008 | -0.200 | 0.6523 | 0.600 | 0.1289 | 0.689 | 0.0742 |
| | R2 | 0.111 | 0.8203 | -0.644 | 0.0977 | 0.600 | 0.1250 | 0.689 | 0.0742 |
| | R3 | **-0.956** | **0.0078** | **-1.000** | **0.0039** | 0.333 | 0.4219 | 0.556 | 0.1641 |
| | R4 | -0.333 | 0.4258 | -0.556 | 0.1641 | 0.644 | 0.0977 | 0.778 | 0.0391 |
| Theta | R1 | 0.333 | 0.4258 | 0.244 | 0.5703 | -0.156 | 0.7344 | -0.289 | 0.4961 |
| | R2 | -0.156 | 0.7344 | -0.067 | 0.9102 | -0.333 | 0.4258 | -0.333 | 0.4258 |
| | R3 | **-1.000** | **0.0039** | -0.867 | 0.0195 | -0.289 | 0.4961 | 0.067 | 0.9102 |
| | R4 | -0.333 | 0.4258 | -0.644 | 0.0977 | -0.067 | 0.9102 | 0.156 | 0.7344 |
| Alpha | R1 | -0.067 | 0.9102 | 0.556 | 0.1641 | -0.111 | 0.8203 | -0.689 | 0.0742 |
| | R2 | -0.778 | 0.0391 | 0.200 | 0.6523 | **-1.000** | **0.0039** | **-1.000** | **0.0039** |
| | R3 | **-0.956** | **0.0078** | -0.511 | 0.2031 | **-1.000** | **0.0039** | **-1.000** | **0.0039** |
| | R4 | -0.689 | 0.0742 | -0.511 | 0.2031 | **-0.867** | **0.0195** | -0.556 | 0.1641 |
| Beta | R1 | 0.600 | 0.1289 | 0.689 | 0.0742 | -0.467 | 0.2500 | -0.689 | 0.0742 |
| | R2 | -0.156 | 0.7344 | 0.511 | 0.2031 | **-0.956** | **0.0078** | **-0.956** | **0.0078** |
| | R3 | **-1.000** | **0.0039** | 0.111 | 0.8203 | **-0.867** | **0.0195** | **-0.867** | **0.0195** |

| | | | | | | | | | |
|---|---|---|---|---|---|---|---|---|---|
| | R4 | -0.378 | 0.3594 | -0.467 | 0.2500 | -0.244 | 0.5703 | -0.067 | 0.9102 |
| Gamma | R1 | 0.556 | 0.1641 | 0.689 | 0.0742 | -0.244 | 0.5469 | -0.556 | 0.1641 |
| | R2 | 0.689 | 0.0742 | 0.644 | 0.0977 | 0.733 | 0.0547 | 0.511 | 0.2031 |
| | R3 | 0.378 | 0.3594 | 0.289 | 0.4961 | 0.533 | 0.1680 | 0.289 | 0.4961 |
| | R4 | -0.244 | 0.5703 | -0.244 | 0.5703 | -0.089 | 0.8516 | 0.067 | 0.9102 |

**Supplemental Table S8: Power-Spectrum Modulation Index for Music Control**

*Effect size r = matched-pairs rank-biserial correlation (Kerby, 2014). Bolded values are significant after BH-FDR correction (q <0.05), applied within group/band across the four within-band tests per comparison type. Music control (n = 8).*

| Band | Round | Total vs. Baseline | | Aperiodic vs. Baseline | | Periodic vs. Baseline | | Aperiodic vs. Periodic (Direct) | |
|---|---|---|---|---|---|---|---|---|---|
| | | r | p | r | p | r | p | r | p |
| Delta | R1 | 0.333 | 0.4609 | 0.389 | 0.3828 | 0.389 | 0.3828 | 0.333 | 0.4609 |
| | R2 | 0.444 | 0.3125 | 0.389 | 0.3828 | 0.500 | 0.2500 | 0.500 | 0.2500 |
| | R3 | -0.556 | 0.1953 | -0.611 | 0.1484 | 0.000 | 1.0000 | 0.833 | 0.0391 |
| | R4 | 0.111 | 0.8438 | 0.056 | 0.9453 | 0.056 | 0.9453 | 0.278 | 0.5469 |
| Theta | R1 | 0.667 | 0.1094 | 0.167 | 0.7422 | 0.889 | 0.0234 | 0.500 | 0.2500 |
| | R2 | 0.500 | 0.2500 | 0.167 | 0.7422 | 0.556 | 0.1953 | 0.389 | 0.3828 |
| | R3 | -0.722 | 0.0781 | -0.611 | 0.1484 | 0.444 | 0.3125 | 0.611 | 0.1484 |
| | R4 | -0.056 | 0.9453 | -0.167 | 0.7422 | 0.667 | 0.1094 | 0.611 | 0.1484 |
| Alpha | R1 | -0.556 | 0.1953 | -0.222 | 0.6406 | -0.556 | 0.1953 | -0.222 | 0.6406 |
| | R2 | -0.722 | 0.0781 | -0.389 | 0.3828 | -0.444 | 0.3125 | -0.111 | 0.8438 |
| | R3 | -0.611 | 0.1484 | -0.889 | 0.0234 | 0.111 | 0.8438 | 0.556 | 0.1953 |
| | R4 | **-1.000** | **0.0078** | -0.500 | 0.2500 | -0.389 | 0.3828 | 0.111 | 0.8438 |
| Beta | R1 | -0.500 | 0.2500 | **-0.944** | **0.0156** | -0.111 | 0.8438 | 0.389 | 0.3828 |
| | R2 | -0.389 | 0.3828 | -0.722 | 0.0781 | 0.389 | 0.3828 | 0.611 | 0.1484 |
| | R3 | -0.556 | 0.1953 | **-1.000** | **0.0078** | 0.111 | 0.8438 | 0.667 | 0.1094 |
| | R4 | **-1.000** | **0.0078** | **-0.944** | **0.0156** | -0.056 | 0.9453 | 0.556 | 0.1953 |
| Gamma | R1 | **-0.944** | **0.0156** | **-1.000** | **0.0078** | -0.222 | 0.6406 | 0.556 | 0.1953 |
| | R2 | **-0.833** | **0.0391** | **-0.833** | **0.0391** | -0.500 | 0.2500 | 0.111 | 0.8438 |
| | R3 | **-0.944** | **0.0156** | **-0.944** | **0.0156** | 0.278 | 0.5469 | 0.556 | 0.1953 |
| | R4 | **-0.889** | **0.0234** | **-0.889** | **0.0234** | -0.278 | 0.5469 | 0.444 | 0.3125 |

#### Supplemental Table S9: LZC Modulation Indices for HVB

*Baseline mean LZC and Round mean LZC are group-mean values (5-s epochs, median-threshold binarization); % Δ is the percent change of the group mean from baseline. MI p and MI r are participant-level modulation-index statistics (matched-pairs rank-biserial correlation; Wilcoxon signed-rank, n = 25), one 4-round BH-FDR family per band; p is bolded where significant after BH-FDR correction, and r is signed (negative = decrease relative to baseline). Cluster p is the sensor-level statistic: the p-value of the strongest positive- or negative-mass cluster (sign shown as + or −) from a two-sided cluster-based permutation test, participant-blocked (n = 25), BH-FDR-corrected within each band across the four rounds; bolded values are significant after-BH-FDR correction. Markers on associated topographic maps indicate cluster membership, not independent sensor-level significance.*

| Band | Round | Baseline mean LZc | Round mean LZc | % Δ | MI p | MI r | Cluster p |
|---|---|---|---|---|---|---|---|
| **Broadband** | R1 | 0.2732 | 0.2853 | +4.46% | **.0042** | **0.655** | **+0.0020** |
| | R2 | | 0.2922 | +6.99% | **.0004** | **0.815** | **+0.0004** |
| | R3 | | 0.2918 | +6.82% | **.0058** | **0.631** | **+0.0104** |
| | R4 | | 0.2883 | +5.53% | **.0186** | **0.538** | **+0.0200** |
| **Delta** | R1 | 0.0761 | 0.0750 | −1.48% | **.0019** | **−0.711** | **−0.0016** |
| | R2 | | 0.0746 | −2.01% | **<.0001** | **−0.895** | **−0.0004** |
| | R3 | | 0.0750 | −1.38% | **.0021** | **−0.705** | **−0.0004** |
| | R4 | | 0.0757 | −0.52% | .4926 | −0.157 | −0.4619 |
| **Theta** | R1 | 0.1530 | 0.1534 | +0.27% | .3130 | 0.231 | +0.2040 |
| | R2 | | 0.1528 | −0.17% | .7982 | 0.058 | −0.2104 |
| | R3 | | 0.1522 | −0.54% | .3533 | −0.212 | −0.3571 |

| | | | | | | | |
|---|---|---|---|---|---|---|---|
| | R4 | | 0.1522 | −0.51% | .3002 | −0.237 | −0.1552 |
| **Alpha** | R1 | 0.1885 | 0.1917 | +1.68% | **.0074** | **0.612** | **+0.0072** |
| | R2 | | 0.1945 | +3.17% | **.0009** | **0.760** | **+0.0004** |
| | R3 | | 0.1960 | +3.95% | **.0001** | **0.883** | **+0.0004** |
| | R4 | | 0.1953 | +3.62% | **.0004** | **0.809** | **+0.0008** |
| **Beta** | R1 | 0.3387 | 0.3410 | +0.68% | **.0422** | **0.465** | **+0.0256** |
| | R2 | | 0.3440 | +1.56% | **.0004** | **0.803** | **+0.0008** |
| | R3 | | 0.3445 | +1.71% | **.0003** | **0.822** | **+0.0004** |
| | R4 | | 0.3445 | +1.72% | **.0002** | **0.858** | **+0.0004** |
| **Gamma** | R1 | 0.3559 | 0.3566 | +0.20% | .0980 | 0.378 | +0.1516 |
| | R2 | | 0.3558 | −0.03% | .8401 | 0.046 | −0.7443 |
| | R3 | | 0.3564 | +0.13% | .5098 | 0.151 | +0.2308 |
| | R4 | | 0.3555 | −0.13% | .3674 | −0.206 | −0.6423 |

#### Supplemental Table S10: LZC in the Music Control

*MI r and MI p are participant-level modulation-index statistics (Wilcoxon signed-rank test, n = 8; r = matched-pairs rank-biserial correlation, positive = increase and negative = decrease relative to baseline). Cluster p is the p-value of the strongest sensor-level cluster per time point (when both a positive and a negative cluster formed, the one with the lower p) from a two-sided cluster-based permutation test on paired differences (cluster-forming threshold p < 0.05; n = 8); — = no candidate cluster formed. All p-values are uncorrected; no result was significant at q < 0.05 after BH-FDR correction across the four time points (MI: within each band; cluster: within each band and direction, absent clusters entered as p = 1).*

| Band | Time point | MI r | MI p | Cluster p |
|---|---|---|---|---|
| Broadband | R1 | −0.83 | 0.0391 | 0.0234 |
| | R2 | −0.67 | 0.1094 | 0.1250 |
| | R3 | −0.33 | 0.4609 | 0.5156 |
| | R4 | −0.78 | 0.0547 | 0.0313 |
| Delta | R1 | −0.11 | 0.8438 | 0.6797 |
| | R2 | −0.17 | 0.7422 | — |
| | R3 | −0.11 | 0.8438 | 0.7109 |
| | R4 | 0.22 | 0.6406 | 0.6250 |
| Theta | R1 | −0.17 | 0.7422 | — |
| | R2 | −0.56 | 0.1953 | 0.4844 |
| | R3 | −0.06 | 0.9453 | — |
| | R4 | −0.67 | 0.1094 | 0.1406 |
| Alpha | R1 | 0.67 | 0.1094 | 0.1016 |
| | R2 | 0.72 | 0.0781 | 0.0703 |
| | R3 | 0.44 | 0.3125 | 0.1797 |
| | R4 | 0.44 | 0.3125 | 0.2109 |
| Beta | R1 | −0.17 | 0.7422 | — |
| | R2 | −0.22 | 0.6406 | — |
| | R3 | −0.94 | 0.0156 | 0.0547 |
| | R4 | −0.50 | 0.2500 | 0.5000 |
| Gamma | R1 | −0.50 | 0.2500 | 0.3359 |
| | R2 | −0.61 | 0.1484 | 0.0234 |
| | R3 | −0.39 | 0.3828 | 0.1172 |
| | R4 | −0.17 | 0.7422 | 0.2266 |

#### Supplemental Table S11: LZC in PMR and Matched HVB sample

MI r and MI p are participant-level modulation-index statistics (Wilcoxon signed-rank test, n = 9; r = matched-pairs rank-biserial correlation, positive = increase relative to baseline). Cluster p is the p-value of the strongest sensor-level cluster per time point from a two-sided cluster-based permutation test on paired differences (cluster-forming threshold $p < 0.05$; n = 9); — = no candidate cluster formed. All p-values are uncorrected; bold = significant after BH-FDR correction across the four time points within each band (MI: $q < 0.05$; cluster: within each band and direction, absent clusters entered as p = 1). Broadband = 1–45 Hz.

##### Panel A: PMR

| Band | Time point | MI r | MI p | Cluster p |
|---|---|---|---|---|
| Broadband | R1 | 0.16 | 0.7344 | **0.0313** |
| | R2 | 0.82 | 0.0273 | **0.0078** |
| | R3 | **1.00** | **0.0039** | **0.0039** |
| | R4 | 0.51 | 0.2031 | 0.1445 |
| Delta | R1 | −0.82 | 0.0273 | **0.0078** |
| | R2 | **−0.96** | **0.0078** | **0.0039** |
| | R3 | −0.73 | 0.0547 | 0.0859 |
| | R4 | −0.38 | 0.3594 | 0.3555 |
| Theta | R1 | 0.56 | 0.1641 | 0.0508 |
| | R2 | 0.16 | 0.7344 | 0.5703 |
| | R3 | 0.24 | 0.5703 | — |
| | R4 | 0.47 | 0.2500 | 0.1719 |
| Alpha | R1 | 0.38 | 0.3594 | — |
| | R2 | 0.87 | 0.0195 | **0.0117** |
| | R3 | 0.82 | 0.0273 | 0.0469 |
| | R4 | 0.51 | 0.2031 | 0.1992 |
| Beta | R1 | **0.78** | **0.0391** | 0.0664 |
| | R2 | **1.00** | **0.0039** | **0.0039** |
| | R3 | **1.00** | **0.0039** | **0.0039** |
| | R4 | **0.82** | **0.0273** | **0.0234** |
| Gamma | R1 | **1.00** | **0.0039** | **0.0078** |
| | R2 | 0.47 | 0.2500 | 0.1680 |
| | R3 | 0.69 | 0.0742 | **0.0078** |
| | R4 | 0.02 | 1.0000 | — |

**Panel B: LZC Matched BW Sample**

| Band | Time point | MI r | MI p | Cluster p |
|---|---|---|---|---|
| Broadband | R1 | 0.24 | 0.5703 | 0.1250 |
| | R2 | 0.16 | 0.7344 | 0.1875 |
| | R3 | −0.16 | 0.7344 | — |
| | R4 | 0.07 | 0.9102 | — |
| Delta | R1 | 0.60 | 0.1289 | 0.0703 |
| | R2 | **1.00** | **0.0039** | **0.0039** |
| | R3 | 0.60 | 0.1289 | 0.1992 |
| | R4 | 0.47 | 0.2500 | 0.3516 |
| Theta | R1 | 0.78 | 0.0391 | 0.0430 |
| | R2 | 0.51 | 0.2031 | 0.1250 |
| | R3 | 0.33 | 0.4258 | 0.5234 |
| | R4 | 0.73 | 0.0547 | 0.0586 |
| Alpha | R1 | 0.73 | 0.0547 | 0.0781 |
| | R2 | −0.20 | 0.6523 | — |
| | R3 | −0.20 | 0.6523 | — |
| | R4 | −0.24 | 0.5703 | — |
| Beta | R1 | −0.11 | 0.8203 | 0.5039 |
| | R2 | −0.56 | 0.1641 | 0.1406 |
| | R3 | −0.38 | 0.3594 | 0.3984 |
| | R4 | 0.11 | 0.8203 | — |
| Gamma | R1 | 0.20 | 0.6523 | 0.2695 |
| | R2 | 0.33 | 0.4258 | 0.1680 |
| | R3 | 0.29 | 0.4961 | — |
| | R4 | −0.11 | 0.8203 | — |

#### Supplemental Table S12: Weighted Phase Lag Index for HVB

*Modulation Index (MI) p and r are participant-level modulation-index statistics (matched-pairs rank-biserial correlation; Wilcoxon signed-rank, n = 25), one 4-round BH-FDR family per band; p is bolded where significant after BH-FDR correction. Cluster p is the sensor-level statistic: the p-value of the strongest positive- or negative-mass cluster (sign shown as + or −) from a two-sided cluster-based permutation test comparing each round's nodal wPLI strength against baseline, participant-blocked (n = 25), BH-FDR-corrected within each band across the four rounds; bolded values are significant after BH-FDR correction. An em-dash indicates no cluster of either sign was retained at that band/round.*

| Band | Round | MI p | MI r | Cluster p |
|---|---|---|---|---|
| Broadband | R1 | .1919 | 0.298 | +.0992 |
| | R2 | .9036 | 0.028 | — |
| | R3 | .9464 | 0.015 | — |
| | R4 | .7366 | −0.077 | — |
| Delta | R1 | **.0303** | **0.495** | **+.0276** |
| | R2 | **.0080** | **0.606** | **+.0064** |
| | R3 | **.0019** | **0.711** | **+.0008** |
| | R4 | **.0058** | **0.631** | **+.0024** |
| Theta | R1 | **.0003** | **0.828** | **+.0004** |
| | R2 | **.0160** | **0.551** | **+.0044** |
| | R3 | **.0264** | **0.508** | **+.0052** |
| | R4 | **.0019** | **0.711** | **+.0008** |
| Alpha | R1 | .6377 | 0.108 | +.5387 |
| | R2 | **.0370** | **−0.477** | −.0440 |
| | R3 | **.0173** | **−0.545** | **−.0120** |
| | R4 | **.0032** | **−0.674** | **−.0044** |
| Beta | R1 | .7570 | 0.071 | — |
| | R2 | .8612 | 0.040 | +.4723 |
| | R3 | .6571 | 0.102 | — |
| | R4 | .3395 | 0.218 | +.3803 |
| Gamma | R1 | .1500 | 0.329 | +.1412 |
| | R2 | .7570 | 0.071 | — |
| | R3 | .5998 | 0.120 | — |
| | R4 | .5449 | −0.138 | — |

#### Supplemental Table S13: Electrode Assignments for Seven Scalp Regions of Interest

*Regions are sensor-space electrode groupings and should not be interpreted as source-localised cortical regions. These groupings were used for the regional wPLI and directed-information analyses.*

| Region | n | Electrodes |
|---|---|---|
| Anterior frontal | 4 | Fp1, Fp2, AF3, AF4 |
| Frontal | 9 | F1, F2, F3, F4, F5, F6, F7, F8, Fz |
| Central | 16 | C1, C2, C3, C4, C5, C6, Cz, FC1, FC2, FC3, FC4, CP1, CP2, CP3, CP4, CPz |
| Left temporal | 6 | T7, P7, FC5, CP5, TP7, TP9 |
| Right temporal | 6 | T8, P8, FC6, CP6, TP8, TP10 |
| Parietal | 9 | P1, P2, P3, P4, P5, P6, P9, P10, Pz |
| Occipital | 13 | PO3, PO4, PO7, PO8, PO9, PO10, POz, O1, O2, O9, O10, Oz, Iz |

**Supplemental Table S14: Pair-wise cross-region Weighted Phase-Lag Index for HVB**

*Only BH-FDR-Significant Edges by Band and Round Shown here (Total Breathwork Sample, n = 25). Of 504 tests (21 unique region-pair edges × 6 bands × 4 rounds), the 63 shown here survived BH-FDR correction within their own 21-edge band-by-round panel; no beta, gamma, or broadband panel retained any edge. Region abbreviations: AF = anterior-frontal, F = frontal, C = central, P = parietal, O = occipital, LT = left-temporal, RT = right-temporal. Baseline wPLI and Event wPLI are group-mean values; MI r is the participant-level modulation index (matched-pairs rank-biserial correlation; Wilcoxon signed-rank, n = 25), signed (negative = decrease relative to baseline); MI p is its associated p-value; q is theBH-FDR-corrected p-value within the 21-edge panel for that band and round. All rows shown are significant at q <0.05; MI p and q values are bolded to indicate significance.*

| Band | Round | Edge | BaselinewPLI | Event wPLI | MI r | MI p | q |
|---|---|---|---|---|---|---|---|
| **Delta** | R2 | F–P | 0.209 | 0.235 | 0.797 | .0005 | .0052 |
| | R2 | P–RT | 0.215 | 0.249 | 0.815 | .0004 | .0052 |
| | R3 | AF–C | 0.215 | 0.234 | 0.538 | .0186 | .0487 |
| | R3 | AF–P | 0.214 | 0.233 | 0.526 | .0214 | .0500 |
| | R3 | F–C | 0.211 | 0.235 | 0.686 | .0027 | .0142 |
| | R3 | F–P | 0.209 | 0.235 | 0.797 | .0005 | .0104 |
| | R3 | F–O | 0.211 | 0.234 | 0.723 | .0016 | .0142 |
| | R3 | C–P | 0.214 | 0.232 | 0.563 | .0138 | .0447 |
| | R3 | C–O | 0.216 | 0.233 | 0.557 | .0149 | .0447 |
| | R3 | C–LT | 0.215 | 0.238 | 0.705 | .0021 | .0142 |
| | R3 | P–RT | 0.215 | 0.240 | 0.612 | .0074 | .0312 |
| **Theta** | R1 | AF–F | 0.223 | 0.251 | 0.582 | .0110 | .0289 |
| | R1 | AF–C | 0.209 | 0.227 | 0.649 | .0045 | .0205 |
| | R1 | AF–P | 0.199 | 0.222 | 0.625 | .0063 | .0205 |
| | R1 | AF–O | 0.199 | 0.220 | 0.625 | .0063 | .0205 |
| | R1 | F–C | 0.212 | 0.234 | 0.742 | .0012 | .0125 |
| | R1 | F–P | 0.205 | 0.223 | 0.637 | .0054 | .0205 |
| | R1 | F–O | 0.205 | 0.220 | 0.557 | .0149 | .0347 |
| | R1 | C–P | 0.208 | 0.236 | 0.618 | .0068 | .0205 |
| | R1 | C–O | 0.206 | 0.235 | 0.791 | .0005 | .0114 |
| | R4 | AF–C | 0.209 | 0.227 | 0.643 | .0049 | .0244 |
| | R4 | AF–O | 0.199 | 0.213 | 0.631 | .0058 | .0244 |
| | R4 | F–O | 0.205 | 0.219 | 0.785 | .0006 | .0077 |
| | R4 | C–P | 0.208 | 0.233 | 0.705 | .0021 | .0144 |
| | R4 | C–O | 0.206 | 0.233 | 0.772 | .0007 | .0077 |

| | | | | | | | |
|---|---|---|---|---|---|---|---|
| | R4 | P–LT | 0.220 | 0.245 | 0.575 | .0119 | .0356 |
| | R4 | O–LT | 0.222 | 0.238 | 0.588 | .0102 | .0356 |
| **Alpha** | R2 | AF–P | 0.359 | 0.322 | −0.631 | .0058 | .0494 |
| | R2 | AF–RT | 0.347 | 0.290 | −0.600 | .0087 | .0494 |
| | R2 | C–LT | 0.345 | 0.294 | −0.705 | .0021 | .0433 |
| | R2 | O–LT | 0.378 | 0.332 | −0.594 | .0094 | .0494 |
| | R3 | AF–C | 0.340 | 0.303 | −0.532 | .0199 | .0322 |
| | R3 | AF–P | 0.359 | 0.308 | −0.637 | .0054 | .0161 |
| | R3 | AF–O | 0.349 | 0.296 | −0.625 | .0063 | .0166 |
| | R3 | AF–RT | 0.347 | 0.286 | −0.711 | .0019 | .0119 |
| | R3 | F–P | 0.351 | 0.308 | −0.502 | .0283 | .0372 |
| | R3 | F–O | 0.348 | 0.297 | −0.575 | .0119 | .0227 |
| | R3 | F–RT | 0.331 | 0.289 | −0.502 | .0283 | .0372 |
| | R3 | C–P | 0.388 | 0.335 | −0.545 | .0173 | .0302 |
| | R3 | C–O | 0.387 | 0.324 | −0.680 | .0029 | .0123 |
| | R3 | C–LT | 0.345 | 0.288 | −0.600 | .0087 | .0203 |
| | R3 | C–RT | 0.355 | 0.300 | −0.717 | .0017 | .0119 |
| | R3 | P–O | 0.422 | 0.362 | −0.520 | .0230 | .0345 |
| | R3 | P–LT | 0.374 | 0.310 | −0.668 | .0035 | .0123 |
| | R3 | P–RT | 0.376 | 0.323 | −0.582 | .0110 | .0227 |
| | R3 | O–LT | 0.378 | 0.311 | −0.698 | .0023 | .0119 |
| | R3 | O–RT | 0.382 | 0.304 | −0.809 | .0004 | .0085 |
| | R4 | AF–C | 0.340 | 0.304 | −0.538 | .0186 | .0300 |
| | R4 | AF–P | 0.359 | 0.312 | −0.705 | .0021 | .0175 |
| | R4 | AF–O | 0.349 | 0.297 | −0.668 | .0035 | .0175 |
| | R4 | AF–RT | 0.347 | 0.302 | −0.545 | .0173 | .0300 |
| | R4 | F–P | 0.351 | 0.312 | −0.588 | .0102 | .0289 |
| | R4 | F–O | 0.348 | 0.302 | −0.625 | .0063 | .0221 |
| | R4 | F–RT | 0.331 | 0.301 | −0.489 | .0324 | .0426 |
| | R4 | C–P | 0.388 | 0.335 | −0.582 | .0110 | .0289 |
| | R4 | C–O | 0.387 | 0.329 | −0.668 | .0035 | .0175 |
| | R4 | C–LT | 0.345 | 0.302 | −0.551 | .0160 | .0300 |
| | R4 | C–RT | 0.355 | 0.313 | −0.551 | .0160 | .0300 |
| | R4 | P–O | 0.422 | 0.358 | −0.655 | .0042 | .0175 |
| | R4 | P–LT | 0.374 | 0.324 | −0.508 | .0264 | .0396 |
| | R4 | P–RT | 0.376 | 0.331 | −0.495 | .0303 | .0424 |
| | R4 | O–LT | 0.378 | 0.321 | −0.569 | .0128 | .0299 |
| | R4 | O–RT | 0.382 | 0.319 | −0.674 | .0032 | .0175 |

#### Supplemental Table S15: Weighted Phase-Lag Index in PMR

MI r and MI p are participant-level modulation-index statistics (Wilcoxon signed-rank test, n = 10; r = matched-pairs rank-biserial correlation, positive = increase relative to baseline). MI p is uncorrected and is bolded together with MI r where significant after BH-FDR correction across the four time points within each band (q < 0.05). Cluster columns show the strongest sensor-level cluster per time point from a two-sided cluster-based permutation test on paired differences (cluster-forming threshold p < 0.05; cluster statistic = summed t, Σt, where the sign of Σt gives direction; df = 9; n = 10; all 1,024 sign-flip permutations). — = no candidate cluster formed; where both a positive and a negative cluster formed (alpha R3), the cluster with the lower p is shown. Cluster p is uncorrected; no cluster or MI result was significant at q < 0.05 (BH-FDR across the four time points within each band and direction; broadband R1 q = 0.102, alpha R1 q = 0.070). Broadband = 1–45 Hz.

| Band | Time point | MI r | MI p | Cluster Σt | Electrodes (n) | Cluster p |
|---|---|---|---|---|---|---|
| Broadband | R1 | −0.85 | 0.0137 | −26.77 | 9 | 0.0254 |
| | R2 | −0.42 | 0.2754 | −11.92 | 4 | 0.0898 |
| | R3 | −0.20 | 0.6250 | −8.03 | 3 | 0.2324 |
| | R4 | 0.27 | 0.4922 | — | — | — |
| Delta | R1 | 0.09 | 0.8457 | 2.32 | 1 | 1.0000 |
| | R2 | 0.05 | 0.9219 | −2.64 | 1 | 0.8906 |
| | R3 | 0.24 | 0.5566 | 2.60 | 1 | 0.8477 |
| | R4 | 0.82 | 0.0195 | 10.68 | 4 | 0.1523 |
| Theta | R1 | −0.02 | 1.0000 | −4.05 | 1 | 0.5391 |
| | R2 | 0.31 | 0.4316 | 2.42 | 1 | 0.8047 |
| | R3 | 0.16 | 0.6953 | −5.40 | 2 | 0.3848 |
| | R4 | 0.60 | 0.1055 | 10.48 | 4 | 0.1582 |
| Alpha | R1 | −0.78 | 0.0273 | −70.61 | 25 | 0.0176 |
| | R2 | −0.60 | 0.1055 | −3.03 | 1 | 0.3574 |
| | R3 | −0.05 | 0.9219 | 2.63 | 1 | 0.5996 |
| | R4 | −0.31 | 0.4316 | −2.72 | 1 | 0.3398 |
| Beta | R1 | −0.16 | 0.6953 | −7.63 | 2 | 0.2188 |
| | R2 | −0.13 | 0.7695 | −2.76 | 1 | 0.6504 |
| | R3 | −0.05 | 0.9219 | −5.78 | 2 | 0.2539 |
| | R4 | 0.20 | 0.6250 | −7.41 | 2 | 0.2246 |
| Gamma | R1 | −0.53 | 0.1602 | −3.27 | 1 | 0.5293 |

| | | | | | |
|---|---|---|---|---|---|
| R2 | −0.64 | 0.0840 | −12.71 | 4 | 0.0957 |
| R3 | −0.42 | 0.2754 | −2.63 | 1 | 0.7129 |
| R4 | 0.05 | 0.9219 | 2.30 | 1 | 0.6523 |

#### Supplemental Table S16: Weighted Phase-Lag Index in the Music Control

*MI r and MI p are participant-level modulation-index statistics (Wilcoxon signed-rank test, n = 8; r = matched-pairs rank-biserial correlation, positive = increase relative to baseline). Cluster p is the p-value of the strongest sensor-level cluster per time point from a two-sided cluster-based permutation test on paired differences (cluster-forming threshold p < 0.05; cluster statistic = summed t on nodal wPLI strength; n = 8); — = no candidate cluster formed; where both a positive and a negative cluster formed, the one with the lower p is shown. All p-values are uncorrected; bold = significant after BH-FDR correction across the four time points within each band (MI: q < 0.05; cluster: within each band and direction, absent clusters entered as p = 1).*

| Band | Time point | MI r | MI p | Cluster p |
|---|---|---|---|---|
| Broadband | R1 | 0.17 | 0.7422 | — |
| | R2 | −0.61 | 0.1484 | — |
| | R3 | 0.33 | 0.4609 | — |
| | R4 | −0.17 | 0.7422 | — |
| Delta | R1 | −0.06 | 0.9453 | 0.5547 |
| | R2 | 0.06 | 0.9453 | — |
| | R3 | 0.22 | 0.6406 | — |
| | R4 | 0.11 | 0.8438 | 0.7109 |
| Theta | R1 | **1.00** | **0.0078** | **0.0078** |
| | R2 | 0.78 | 0.0547 | 0.1172 |
| | R3 | 0.50 | 0.2500 | 0.3047 |
| | R4 | **0.89** | **0.0234** | 0.1250 |
| Alpha | R1 | −0.50 | 0.2500 | 0.3516 |
| | R2 | −0.61 | 0.1484 | 0.3750 |
| | R3 | −0.33 | 0.4609 | — |
| | R4 | −0.44 | 0.3125 | — |
| Beta | R1 | 0.11 | 0.8438 | 0.3359 |
| | R2 | −0.39 | 0.3828 | 0.3828 |
| | R3 | 0.22 | 0.6406 | — |
| | R4 | −0.56 | 0.1953 | — |
| Gamma | R1 | 0.00 | 1.0000 | — |
| | R2 | −0.39 | 0.3828 | — |
| | R3 | 0.28 | 0.5469 | — |
| | R4 | −0.06 | 0.9453 | — |

#### Supplemental Table S17: Net Directed Information for Frontal–Occipital Regions

*Signed group-mean lag curves (NetDI_event − NetDI_B, or the indicated epoch-to-epoch contrast) were compared via two-sided participant sign-flip cluster-based permutation (5,000 permutations, cluster-forming p <0.05, seed 42) across 40 lags (4–160 ms). One row is shown per comparison: the single strongest cluster entered into the 10-comparison BH-FDR family for this ROI pair (secondary, non-selected clusters from the same comparison are omitted). Mean Δ is the cluster's mean signed difference in directed information (bits); positive values indicate a shift toward frontal→occipital information flow. R1 vs R2 retained no cluster of either sign. Bolded values are significant afterBH-FDR correction (q <0.05).*

| Comparison | Lag (ms) | Cluster mass | Mean Δ (bits) | Direction | Cluster p | FDR q | Sig. |
|---|---|---|---|---|---|---|---|
| B vs R1 | 16–68 | 47.03 | **.0116** | positive | **.0002** | **.0020** | **Yes** |
| B vs R2 | 24–68 | 40.51 | **.0144** | positive | **.0016** | **.0040** | **Yes** |
| B vs R3 | 24–64 | 40.24 | **.0169** | positive | **.0016** | **.0040** | **Yes** |
| B vs R4 | 24–64 | 39.10 | **.0172** | positive | **.0006** | **.0030** | **Yes** |
| R1 vs R2 | — | — | — | none | 1.0000 | 1.000 | No |
| R1 vs R3 | 44–56 | 11.20 | .0065 | positive | .0858 | .1470 | No |
| R1 vs R4 | 44–52 | 7.09 | .0061 | positive | .1050 | .1500 | No |
| R2 vs R3 | 48–52 | 4.28 | .0041 | positive | .2024 | .2530 | No |
| R2 vs R4 | 12–12 | 2.27 | .0028 | positive | .3234 | .3593 | No |
| R3 vs R4 | 8–16 | 7.45 | .0020 | positive | .0882 | .1470 | No |

**Supplemental Table S18: Peak-Lag Net Directed Information for Frontal–Occipital Regions**

*For each round, the participant-paired signed ΔNetDI lag curve (event minus baseline) was evaluated at the lag of largest absolute group-mean excursion (peak selection by |Δ|); exact ties selected the earliest lag. Mean peak ΔNetDI ± SEM is the group-mean signed value (bits) at that lag. Cluster window and cluster p are from the lag-resolved two-sided cluster-permutation map (same procedure as Table S17). Within-pair q is the BH-FDR correction across R1–R4 for this ROI pair; global q is the correction across all 21 ROI pairs × R1–R4 (84 maps). All four rounds are significant at both correction levels (n = 25 participants).*

| Round | Peak lag (ms) | Mean peak ΔNetDI (bits) | Cluster window (ms) | Cluster p | Within-pair q | Global q |
|---|---|---|---|---|---|---|
| R1 | 44 | .0186 ±0.0051 | 16–68 | **.0002** | **.0008** | **.0168** |
| R2 | 48 | .0216 ±0.0060 | 24–68 | **.0016** | **.0016** | **.0336** |
| R3 | 48 | .0256 ±0.0063 | 24–64 | **.0016** | **.0016** | **.0336** |
| R4 | 48 | .0249 ±0.0067 | 24–64 | **.0006** | **.0012** | **.0252** |

**Supplemental Table S19: Absolute Net Directed Information from Occipital to Four Other Regions**

*(Total Breathwork Sample, n = 25). For each connection, the participant-level change was mean(|NetDI| over R1–R4) − |NetDI| at baseline, tested against zero across 40 lags (4–160 ms) with a two-sided participant sign-flip cluster-based permutation test (5,000 permutations, cluster-forming p < 0.05, seed 42). Both the strongest negative and the strongest positive cluster are shown for each connection (none = no cluster exceeded the cluster-forming threshold). FDR q is given for the cluster with the lower map-level p, which was the one entered into the BH-FDR family across the four occipital connections; — = not entered into the family or not applicable. Mean Δ is the cluster's mean difference in |NetDI| (bits); negative values indicate a reduction in the magnitude of directed information (weaker directional asymmetry), positive values an increase. Bold = significant after BH-FDR (q < 0.05). Cluster mass = sum of t-values across the lags in the cluster.*

| Connection | Cluster sign | Lag (ms) | Cluster mass | Mean Δ (bits) | Cluster p | FDR q | Sig. |
|---|---|---|---|---|---|---|---|
| Occipital–Frontal | negative | 32–64 | −25.53 | −0.0142 | **0.0068** | **0.0272** | **Yes** |
| | positive | 8–12 | 6.16 | 0.0013 | 0.2062 | — | — |
| Occipital–Anterior frontal | negative | 28–52 | −16.25 | −0.0094 | 0.0480 | 0.0725 | No |
| | positive | none | — | — | — | — | — |
| Occipital–Central | negative | 152–156 | −4.53 | −0.0051 | 0.2610 | — | — |
| | positive | 4–16 | 12.89 | 0.0034 | 0.0544 | 0.0725 | No |
| Occipital–Parietal | negative | none | — | — | — | — | — |
| | positive | 12–16 | 4.67 | 0.0065 | 0.1376 | 0.1376 | No |

**<u>Supplemental Table S20: Associations Between Neural Metrics and Real-Time ASC Ratings</u>**

*(Total Breathwork Sample, n = 25). Two-sided parametric repeated-measures correlations (rmcorr; Bakdash & Marusich, 2017) between neural metrics and the real-time ASC rating given at the start (Panel A) or end (Panel B) of the same resting-state epoch, across baseline and R1–R4 (n = 25 participants × 5 time points = 125 observations, df = 99). For frontal–occipital Δ|NetDI| (|NetDI| at the group peak lag minus baseline), only R1–R4 were included (100 observations, df = 74). p-values are uncorrected. q = BH-FDR-adjusted p across the 25 tests within each panel (4 neural metrics × 6 frequency bands + 1 NetDI measure; total power excluded). Bold = q < 0.05. Broadband = 1–45 Hz. LZC = Lempel–Ziv complexity; wPLI = weighted phase-lag index; NetDI = net directed information.*

*Panel A. ASC rating at the start of each resting-state epoch*

| Neural metric | Band | r | t | df | p | q (BH-FDR) |
|---|---|---|---|---|---|---|
| Aperiodic power | Broadband | 0.042 | 0.42 | 99 | 0.679 | 0.800 |
| | Delta | 0.034 | 0.34 | 99 | 0.736 | 0.800 |
| | Theta | −0.034 | −0.34 | 99 | 0.736 | 0.800 |
| | Alpha | −0.023 | −0.23 | 99 | 0.820 | 0.855 |
| | Beta | 0.059 | 0.59 | 99 | 0.557 | 0.774 |
| | Gamma | 0.130 | 1.31 | 99 | 0.194 | 0.323 |
| Periodic power | Broadband | **−0.476** | −5.38 | 99 | **< 0.001** | **< 0.001** |
| | Delta | **0.254** | 2.61 | 99 | **0.010** | **0.026** |
| | Theta | −0.111 | −1.11 | 99 | 0.268 | 0.419 |
| | Alpha | **−0.607** | −7.59 | 99 | **< 0.001** | **< 0.001** |
| | Beta | **−0.408** | −4.44 | 99 | **< 0.001** | **< 0.001** |
| | Gamma | 0.212 | 2.16 | 99 | 0.033 | 0.064 |
| LZC | Broadband | **0.338** | 3.58 | 99 | **< 0.001** | **0.002** |
| | Delta | **−0.240** | −2.46 | 99 | **0.016** | **0.035** |
| | Theta | −0.167 | −1.69 | 99 | 0.094 | 0.169 |
| | Alpha | **0.550** | 6.55 | 99 | **< 0.001** | **< 0.001** |
| | Beta | **0.525** | 6.15 | 99 | **< 0.001** | **< 0.001** |
| | Gamma | 0.015 | 0.15 | 99 | 0.884 | 0.884 |
| wPLI nodal strength | Broadband | −0.034 | −0.34 | 99 | 0.733 | 0.800 |
| | Delta | **0.294** | 3.06 | 99 | **0.003** | **0.008** |
| | Theta | 0.222 | 2.27 | 99 | 0.025 | 0.053 |
| | Alpha | **−0.294** | −3.07 | 99 | **0.003** | **0.008** |

| | | | | | | |
|---|---|---|---|---|---|---|
| | Beta | 0.077 | 0.77 | 99 | 0.441 | 0.649 |
| | Gamma | −0.043 | −0.42 | 99 | 0.672 | 0.800 |
| Frontal–occipital Δ\|NetDI\| | Peak, 4–160 ms | **−0.370** | −3.42 | 74 | **0.001** | **0.004** |

*Panel B. ASC rating at the end of each resting-state epoch*

| Neural metric | Band | r | t | df | p | q (BH-FDR) |
|---|---|---|---|---|---|---|
| Aperiodic power | Broadband | 0.075 | 0.75 | 99 | 0.457 | 0.672 |
| | Delta | 0.079 | 0.79 | 99 | 0.431 | 0.672 |
| | Theta | −0.005 | −0.05 | 99 | 0.958 | 0.958 |
| | Alpha | −0.035 | −0.35 | 99 | 0.727 | 0.957 |
| | Beta | 0.006 | 0.05 | 99 | 0.956 | 0.958 |
| | Gamma | 0.055 | 0.55 | 99 | 0.584 | 0.811 |
| Periodic power | Broadband | **−0.389** | −4.21 | 99 | **< 0.001** | **< 0.001** |
| | Delta | 0.164 | 1.65 | 99 | 0.101 | 0.254 |
| | Theta | −0.011 | −0.11 | 99 | 0.916 | 0.958 |
| | Alpha | **−0.529** | −6.20 | 99 | **< 0.001** | **< 0.001** |
| | Beta | **−0.313** | −3.28 | 99 | **0.001** | **0.007** |
| | Gamma | 0.141 | 1.42 | 99 | 0.158 | 0.305 |
| LZC | Broadband | 0.153 | 1.54 | 99 | 0.127 | 0.288 |
| | Delta | −0.134 | −1.35 | 99 | 0.180 | 0.322 |
| | Theta | −0.143 | −1.44 | 99 | 0.152 | 0.305 |
| | Alpha | **0.490** | 5.59 | 99 | **< 0.001** | **< 0.001** |
| | Beta | **0.342** | 3.63 | 99 | **< 0.001** | **0.003** |
| | Gamma | −0.025 | −0.25 | 99 | 0.801 | 0.958 |
| wPLI nodal strength | Broadband | 0.008 | 0.08 | 99 | 0.938 | 0.958 |
| | Delta | 0.205 | 2.09 | 99 | 0.040 | 0.110 |
| | Theta | 0.214 | 2.18 | 99 | 0.032 | 0.100 |
| | Alpha | −0.223 | −2.27 | 99 | 0.025 | 0.090 |
| | Beta | 0.106 | 1.06 | 99 | 0.291 | 0.485 |
| | Gamma | −0.012 | −0.12 | 99 | 0.907 | 0.958 |
| Frontal–occipital Δ\|NetDI\| | Peak, 4–160 ms | **−0.298** | −2.69 | 74 | **0.009** | **0.037** |

**Supplemental Table S21: Correlations Between Peak-ASC Neural Metrics and Post-Session Survey Scores**

*(Total Breathwork Sample, n = 25). Pearson correlations (n = 25, df = 23) between each participant's peak-minus-baseline neural metric, taken from the resting-state epoch with the highest real-time ASC rating, and the corresponding post-session survey score. p-values are uncorrected. q = BH-FDR-adjusted p across the 25 tests within each questionnaire (4 neural metrics × 6 frequency bands + 1 NetDI measure; total power excluded); no test was significant at q < 0.05. Broadband = 1–45 Hz. Non-parametric (Spearman) results for DED, which deviated from normality, are in Supp. Table S14.*

*Panel A. MEQ30 total score*

| Neural metric | Band | r | t | df | p | q (BH-FDR) |
|---|---|---|---|---|---|---|
| Aperiodic power | Broadband | −0.035 | −0.17 | 23 | 0.869 | 0.999 |
| | Delta | −0.020 | −0.10 | 23 | 0.923 | 0.999 |
| | Theta | −0.151 | −0.73 | 23 | 0.470 | 0.999 |
| | Alpha | −0.153 | −0.74 | 23 | 0.466 | 0.999 |
| | Beta | −0.051 | −0.24 | 23 | 0.809 | 0.999 |
| | Gamma | 0.019 | 0.09 | 23 | 0.926 | 0.999 |
| Periodic power | Broadband | −0.063 | −0.30 | 23 | 0.765 | 0.999 |
| | Delta | −0.128 | −0.62 | 23 | 0.543 | 0.999 |
| | Theta | 0.139 | 0.67 | 23 | 0.507 | 0.999 |
| | Alpha | 0.026 | 0.13 | 23 | 0.901 | 0.999 |
| | Beta | −0.024 | −0.11 | 23 | 0.911 | 0.999 |
| | Gamma | −0.565 | −3.29 | 23 | 0.003 | 0.081 |
| LZC | Broadband | −0.006 | −0.03 | 23 | 0.977 | 0.999 |
| | Delta | 0.206 | 1.01 | 23 | 0.323 | 0.999 |
| | Theta | 0.154 | 0.75 | 23 | 0.463 | 0.999 |
| | Alpha | 0.233 | 1.15 | 23 | 0.263 | 0.999 |
| | Beta | 0.049 | 0.24 | 23 | 0.815 | 0.999 |
| | Gamma | 0.377 | 1.95 | 23 | 0.064 | 0.794 |
| wPLI nodal strength | Broadband | −0.099 | −0.47 | 23 | 0.639 | 0.999 |
| | Delta | −0.139 | −0.67 | 23 | 0.509 | 0.999 |
| | Theta | −0.044 | −0.21 | 23 | 0.836 | 0.999 |
| | Alpha | 0.000 | 0.00 | 23 | 0.999 | 0.999 |

| | | | | | | |
|---|---|---|---|---|---|---|
| | Beta | −0.157 | −0.76 | 23 | 0.454 | 0.999 |
| | Gamma | −0.013 | −0.06 | 23 | 0.951 | 0.999 |
| Frontal→occipital NetDI | Lag-normalized signed AUC | 0.293 | 1.47 | 23 | 0.155 | 0.999 |

*Panel B. 11D-ASC Oceanic Boundlessness (OBN)*

| Neural metric | Band | r | t | df | p | q (BH-FDR) |
|---|---|---|---|---|---|---|
| Aperiodic power | Broadband | −0.133 | −0.64 | 23 | 0.527 | 0.874 |
| | Delta | −0.113 | −0.54 | 23 | 0.592 | 0.874 |
| | Theta | −0.253 | −1.25 | 23 | 0.223 | 0.874 |
| | Alpha | −0.225 | −1.11 | 23 | 0.279 | 0.874 |
| | Beta | −0.067 | −0.32 | 23 | 0.750 | 0.874 |
| | Gamma | 0.048 | 0.23 | 23 | 0.821 | 0.874 |
| Periodic power | Broadband | −0.159 | −0.77 | 23 | 0.449 | 0.874 |
| | Delta | −0.136 | −0.66 | 23 | 0.518 | 0.874 |
| | Theta | 0.078 | 0.38 | 23 | 0.709 | 0.874 |
| | Alpha | −0.039 | −0.19 | 23 | 0.852 | 0.874 |
| | Beta | −0.142 | −0.69 | 23 | 0.500 | 0.874 |
| | Gamma | −0.449 | −2.41 | 23 | 0.024 | 0.607 |
| LZC | Broadband | 0.153 | 0.74 | 23 | 0.466 | 0.874 |
| | Delta | 0.163 | 0.79 | 23 | 0.436 | 0.874 |
| | Theta | 0.085 | 0.41 | 23 | 0.686 | 0.874 |
| | Alpha | 0.142 | 0.69 | 23 | 0.499 | 0.874 |
| | Beta | 0.037 | 0.18 | 23 | 0.862 | 0.874 |
| | Gamma | 0.269 | 1.34 | 23 | 0.194 | 0.874 |
| wPLI nodal strength | Broadband | 0.102 | 0.49 | 23 | 0.627 | 0.874 |
| | Delta | −0.169 | −0.82 | 23 | 0.420 | 0.874 |
| | Theta | 0.033 | 0.16 | 23 | 0.874 | 0.874 |
| | Alpha | 0.139 | 0.67 | 23 | 0.507 | 0.874 |
| | Beta | 0.063 | 0.30 | 23 | 0.766 | 0.874 |
| | Gamma | 0.037 | 0.18 | 23 | 0.862 | 0.874 |
| Frontal→occipital NetDI | Lag-normalized signed AUC | 0.259 | 1.28 | 23 | 0.212 | 0.874 |

*Panel C. 11D-ASC Visionary Restructuralization (VR)*

| Neural metric | Band | r | t | df | p | q (BH-FDR) |
|---|---|---|---|---|---|---|
| Aperiodic power | Broadband | −0.201 | −0.99 | 23 | 0.334 | 0.989 |
| | Delta | −0.212 | −1.04 | 23 | 0.308 | 0.989 |
| | Theta | −0.121 | −0.58 | 23 | 0.566 | 0.989 |
| | Alpha | 0.019 | 0.09 | 23 | 0.927 | 0.989 |
| | Beta | 0.086 | 0.41 | 23 | 0.684 | 0.989 |
| | Gamma | 0.063 | 0.30 | 23 | 0.765 | 0.989 |
| Periodic power | Broadband | −0.054 | −0.26 | 23 | 0.798 | 0.989 |
| | Delta | −0.402 | −2.11 | 23 | 0.046 | 0.576 |
| | Theta | 0.003 | 0.01 | 23 | 0.989 | 0.989 |
| | Alpha | 0.074 | 0.35 | 23 | 0.727 | 0.989 |
| | Beta | 0.025 | 0.12 | 23 | 0.906 | 0.989 |
| | Gamma | −0.419 | −2.21 | 23 | 0.037 | 0.576 |
| LZC | Broadband | 0.084 | 0.41 | 23 | 0.688 | 0.989 |
| | Delta | 0.094 | 0.45 | 23 | 0.656 | 0.989 |
| | Theta | 0.266 | 1.32 | 23 | 0.198 | 0.989 |
| | Alpha | 0.170 | 0.83 | 23 | 0.417 | 0.989 |
| | Beta | 0.019 | 0.09 | 23 | 0.928 | 0.989 |
| | Gamma | 0.149 | 0.72 | 23 | 0.478 | 0.989 |
| wPLI nodal strength | Broadband | −0.005 | −0.02 | 23 | 0.982 | 0.989 |
| | Delta | −0.007 | −0.03 | 23 | 0.975 | 0.989 |
| | Theta | −0.144 | −0.70 | 23 | 0.493 | 0.989 |
| | Alpha | 0.212 | 1.04 | 23 | 0.310 | 0.989 |
| | Beta | −0.063 | −0.30 | 23 | 0.763 | 0.989 |
| | Gamma | −0.053 | −0.26 | 23 | 0.800 | 0.989 |
| Frontal→occipital NetDI | Lag-normalized signed AUC | 0.318 | 1.61 | 23 | 0.121 | 0.989 |

*Panel D. 11D-ASC Dread of Ego Dissolution (DED)*

| Neural metric | Band | r | t | df | p | q (BH-FDR) |
|---|---|---|---|---|---|---|
| Aperiodic power | Broadband | −0.060 | −0.29 | 23 | 0.774 | 0.993 |
| | Delta | −0.103 | −0.50 | 23 | 0.624 | 0.993 |
| | Theta | 0.115 | 0.56 | 23 | 0.583 | 0.993 |

| | | | | | | |
|---|---|---|---|---|---|---|
| | Alpha | 0.246 | 1.22 | 23 | 0.235 | 0.969 |
| | Beta | 0.235 | 1.16 | 23 | 0.258 | 0.969 |
| | Gamma | 0.186 | 0.91 | 23 | 0.372 | 0.993 |
| Periodic power | Broadband | −0.048 | −0.23 | 23 | 0.818 | 0.993 |
| | Delta | −0.166 | −0.80 | 23 | 0.429 | 0.993 |
| | Theta | 0.037 | 0.18 | 23 | 0.861 | 0.993 |
| | Alpha | 0.054 | 0.26 | 23 | 0.797 | 0.993 |
| | Beta | −0.032 | −0.15 | 23 | 0.879 | 0.993 |
| | Gamma | −0.300 | −1.51 | 23 | 0.146 | 0.969 |
| LZC | Broadband | 0.001 | 0.01 | 23 | 0.996 | 0.996 |
| | Delta | 0.076 | 0.36 | 23 | 0.719 | 0.993 |
| | Theta | 0.072 | 0.34 | 23 | 0.733 | 0.993 |
| | Alpha | 0.135 | 0.65 | 23 | 0.520 | 0.993 |
| | Beta | 0.072 | 0.35 | 23 | 0.733 | 0.993 |
| | Gamma | 0.312 | 1.57 | 23 | 0.129 | 0.969 |
| wPLI nodal strength | Broadband | 0.232 | 1.14 | 23 | 0.265 | 0.969 |
| | Delta | 0.229 | 1.13 | 23 | 0.271 | 0.969 |
| | Theta | −0.011 | −0.05 | 23 | 0.958 | 0.996 |
| | Alpha | 0.051 | 0.25 | 23 | 0.808 | 0.993 |
| | Beta | 0.023 | 0.11 | 23 | 0.913 | 0.993 |
| | Gamma | 0.265 | 1.32 | 23 | 0.200 | 0.969 |
| Frontal→occipital NetDI | Lag-normalized signed AUC | 0.204 | 1.00 | 23 | 0.329 | 0.993 |

*DED (11D-ASC), Spearman's rho*

| Neural metric | Band | ρ | t | df | p | q (BH-FDR) |
|---|---|---|---|---|---|---|
| Aperiodic power | Broadband | −0.015 | −0.07 | 23 | 0.942 | 0.956 |
| | Delta | 0.058 | 0.28 | 23 | 0.784 | 0.956 |
| | Theta | −0.012 | −0.06 | 23 | 0.956 | 0.956 |
| | Alpha | 0.180 | 0.88 | 23 | 0.389 | 0.956 |
| | Beta | 0.221 | 1.09 | 23 | 0.289 | 0.956 |
| | Gamma | 0.188 | 0.92 | 23 | 0.369 | 0.956 |
| Periodic power | Broadband | −0.187 | −0.91 | 23 | 0.371 | 0.956 |
| | Delta | −0.076 | −0.37 | 23 | 0.718 | 0.956 |

| | | | | | | |
|---|---|---|---|---|---|---|
| | Theta | 0.018 | 0.08 | 23 | 0.933 | 0.956 |
| | Alpha | −0.077 | −0.37 | 23 | 0.715 | 0.956 |
| | Beta | −0.102 | −0.49 | 23 | 0.627 | 0.956 |
| | Gamma | −0.278 | −1.39 | 23 | 0.178 | 0.956 |
| LZC | Broadband | 0.016 | 0.08 | 23 | 0.939 | 0.956 |
| | Delta | 0.068 | 0.33 | 23 | 0.748 | 0.956 |
| | Theta | 0.128 | 0.62 | 23 | 0.541 | 0.956 |
| | Alpha | 0.228 | 1.13 | 23 | 0.272 | 0.956 |
| | Beta | 0.088 | 0.43 | 23 | 0.674 | 0.956 |
| | Gamma | 0.329 | 1.67 | 23 | 0.108 | 0.956 |
| wPLI nodal strength | Broadband | 0.275 | 1.37 | 23 | 0.184 | 0.956 |
| | Delta | 0.128 | 0.62 | 23 | 0.543 | 0.956 |
| | Theta | 0.097 | 0.47 | 23 | 0.645 | 0.956 |
| | Alpha | 0.015 | 0.07 | 23 | 0.945 | 0.956 |
| | Beta | 0.120 | 0.58 | 23 | 0.568 | 0.956 |
| | Gamma | 0.346 | 1.77 | 23 | 0.090 | 0.956 |
| Frontal–occipital Δ\|NetDI\| | Peak, 4–160 ms | −0.236 | −1.17 | 23 | 0.256 | 0.956 |

#### Supplemental Text S1: Breathwork Intervention & Facilitation Protocol

**Background**

The ESI RESET study explores the potential of conscious-connected breathwork (CCB) to induce altered states of consciousness, to reduce psychopathology, to improve well-being and to enhance socio-emotional functioning in experienced breathwork practitioners with a background of childhood trauma. To this end, we will conduct a 1-session intervention featuring one individual CCB session with in-person facilitation (see below).

**Participants**

Recruited participants have experienced at least 5 conscious-connected breathwork sessions, have experienced at least some adverse childhood events, and show mild depression scores. However, participants also undergo a structured clinical interview, and are excluded if psychopathological symptoms reach the level of a clinical diagnosis. Similarly, participants are screened beforehand for potential contraindications of CCB, including epilepsy, schizophrenia, psychosis, heart and lung conditions and pregnancy.

##### Structure of intervention

The intervention includes one individual CCB session, lasting approximately one hour, with preparation and integration before and after the session (see below). The breathwork session during the intervention will be led by a trained breathwork facilitator.

An optional control session includes one individual progressive muscle relaxation session, lasting approximately one hour, and led by a trained facilitator.

Survey questionnaires will be collected before and after the intervention, and self-reports of the breathwork experience itself are collected online after the CCB session.

During the intervention and optional control session, EEG, ECG, and respiratory measurements will be taken.

#### Intervention Session

##### Schedule (Timings are approximate)

| **Activity** | **Duration (min)** |
|---|---|
| Introduction to lab & consent forms (first session) | 5 |
| Intention setting & breathwork expectations (only facilitator & participant)<br>+ Explain experimental procedure:<br> + Timing of session structure<br> + ASC handsignals<br>+ Demonstrate breathing technique<br>+ Discuss the role of touch & touch consent<br> + Sign touch consent form<br>+ Set facilitation expectations<br>+ Intention setting | 20 |
| EEG electrode, respiration belt & ECG fitting<br>+ Fit cap onto participants head<br>+ Test breathing on the mattress with the cap<br>+ Gel electrodes<br>+ Participant lays on mattress<br>+ ECG electrode placement<br>+ Respiration belt fitting & signal testing | 40 |
| Meditative body scan<br>+ Remind participant of intentions<br>+ Conduct a guided felt sense meditation before experiment officially begins | 10 |
| Circular breathwork (13 min active breath + 4 minute rest x 4) | 80 |
| Sharing and integration (only facilitator & participant)<br>+ Offer participant time to return and unfit from the electrodes<br>+ Offer sink for hair washing | 15 |
| **BREAK** | **30** |
| Post- Intervention Surveys | 30 |
| Video annotation (Experimentor and participant, facilitator presence optional) | 30 |
| **TOTAL DURATION** | **4h30** |

Facilitators are available for 1 on 1 chats post-session!

##### Introduction & Experimental Procedure

Welcome to the study. Consent forms to be signed and pre-intervention surveys to be completed if not completed prior to visit.

Members of the facilitation and research team introduce themselves.

##### General remarks

- Everything the facilitator team says (apart from acute safety concerns) is an invitation
- Invitation to explore ‘one step outside the comfort zone’ – not too familiar, not too overwhelming
- Invitation to ‘pause’ intellectual analysis for a couple of hours and open up for experience-driven process (emphasizing that analysis is very welcome, but maybe more helpful before and after the session than during)

**Procedure explanation**

In both sessions, you will be fitted with an EEG cap, ECG electrodes, and a respiratory belt. These tools will measure your brain wave activity, the electrical activity of your heart, and your respiration. You will be guided through a CCB breathwork session. As you proceed through the breathwork session, we will ask you at regular intervals to report your subjective experience depth, reported as your “Altered State of Consciousness” (ASC) depth. 1 corresponds with normal waking consciousness, where 5 would indicate your upper threshold for your consciousness being altered, whatever that means to you. When prompted by a tone, you will raise your hand to report a number between 1-5 with your fingers to report your experience depth in that moment. Following this report, you will cease your conscious- connected breath, breathe normally, and lie still for 4 minutes to record your EEG signal. There will be another tone to signal the continuation of the conscious connected breathing pattern, along with guidance from the facilitator.

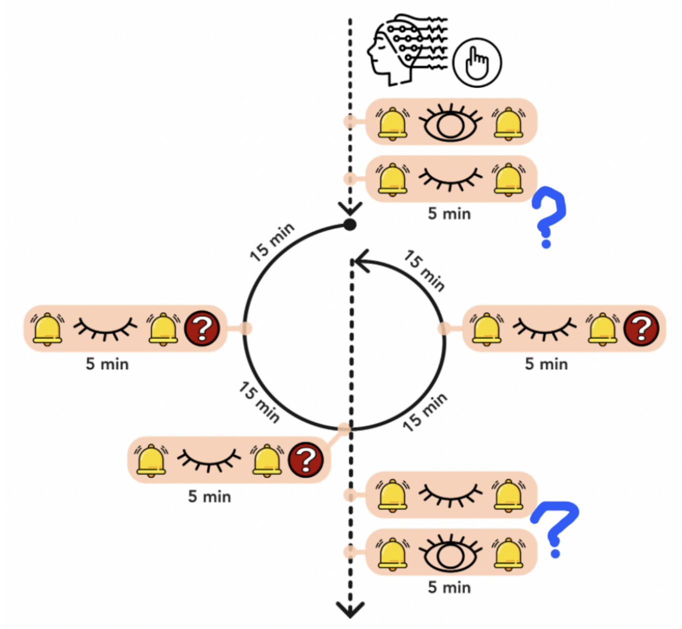

##### Conscious Connected Breathwork Demonstration

You are here because you are an experienced practitioner of Conscious Connected Breathwork (CCB). This demonstration will serve as a reminder of the breathing pattern used in our study. This CCB session will be 80 minutes, including four 13 minute bouts of CCB breathing followed by 4 minutes of stillness for the EEG recording.

##### Demonstrate breathing technique

- Deep inhale, relaxed exhale, connected in a continuous circle, with no pause between the inhale and exhale, or between the exhale and inhale
- Recommended breathing through mouth to increase the respiratory rate, but circular breath can be done through the nose.
    - Mention if the intensity of the mouth breath becomes too much, participant may continue circular breath through the nose
- Demonstrate the full expansion of the belly and rib cage, that our intention is to deepen the breath by using the full capacity of the respiratory system.
    - Optional: Explain the lungs expand in 360 degrees, not only forward or upward, this can increase somatic awareness of what a full breath looks like.
    - Optional: Mention that during breathing on a daily basis, there is a small pause at the bottom of the exhale and demonstrate what circular breath looks like in comparison.
- Describe that the focus of this practice is to increase respiratory volume, which can be achieved through increasing the depth of the breath, pace of the breath, or both. The intention is not on achieving a specific pace of breath, but just to maintain the circular pattern of deep breathing, though the respiratory rate can increase
- Invitation to practice some consciously-connected breaths together, increasing somatic awareness of the sensations of full belly & chest breathing

##### Potential Effects

o Typical initial breathwork effects: Skin tingling, dizziness, dry mouth, tetany (muscle cramps in the hands, feet, or jaw), temperature changes such a hot flashes, sweating, or cold hands and feet

o Typical elements of breathwork experiences: Physical (movements, noises); Emotional (whole spectrum of emotions); Cognitive (memories, new perspectives and insights); Spiritual (universal connection etc.); Visions (perceiving things that are not physically present); 'Lights out' (not in sense of coma but e.g. deep rest, losing track of time etc.); Coming up against cognitive defense mechanisms (e.g. 'Am I doing this right?', boredom, or desire to discontinue circular breath)

##### Breathing with the EEG cap

While our intention is to create as naturalistic a breathwork experience as possible, it is also of high importance to collect quality data for our study. There are a few points to keep in mind about breathing with the EEG cap fitted:

- During the breathing periods, you are free to move organically, but be mindful of unfitting the cap. For example, we discourage movements that might significantly alter your head position against the pillow.
- During the recording periods, you will lie as still as possible to ensure high quality recording

**The role of facilitation and supportive touch**

The role of the facilitator in this experimental session is to support a full naturalistic breathwork experience, including verbal, physical, and emotional support. This includes:

- Encouraging the deepening of the breath and expansion of the respiratory organs, verbally and with physical touch or massage
- Reminding the participant to stay in contact with their conscious connected breath, verbally or with physical touch
- Verbally guiding the participant through the segments of the session (ie: periods of breathing/ period of stillness), as well as guiding them through the progressive building of intensity, and the progressive settling after the peak
- Supporting emotional processes of participants, possibly relevant to the intentions set before the session
    - Facilitator will not guide or lead a participant into specific emotional experience, but support them in relevant ways, should an expression occur
- Facilitators are purely here to support participants' processes; open, direct feedback (e.g. if one needs more or less support than provided) is absolutely welcome
- Mention that if you received touch, it is not a 'correction', you are not doing the practice 'wrong', its an invitation to deepen into certain aspects of your physical or emotional experience
- Demonstrate examples of facilitator touch (e.g. touch on shoulder; touch on belly to assist breathing, support at feet)
- Invitation: If participants want to point out specific touch preferences (e.g. body parts that are off-limits), they can tell facilitators individually before the breathwork
- Invitation: Participants can always change their mind about consent to touch throughout the entire breathwork session, this can be done through either verbal or non-verbal cues. Non verbal cues may include shaking the head, or removal of facilitator's hands.
- Participants may also be encouraged to communicate forms of touch that they know they like, for example touches to the heart or the feet. This can also be communicated before or during the session verbally or non-verbally, for example "could you support my heart?" or placing of facilitators hands onto an (appropriate) body part such as the heart or hands

- Hands off practice: Explanation that even though open feedback is welcome, it can still be difficult to give that feedback in the session.
    - Practice: Facilitator applies a few of the demonstrated touches on participant lying down, participant is asked to practice both verbal (e.g. 'Hands off!') and non-verbal (e.g. pushing hand away) feedback to decline or redirect touch. Invitation is to practice 'impolite' forms of declining in order to allow participants to give direct feedback without worrying about politeness or considerations towards facilitators' feelings during the session.
    - Set the expectation that if one form of physical touch is not wanted, facilitator will verbally check in about other forms of touch before proceeding with touch
- Mention that if participants need support throughout the session, either with their breath or with an emotional process, they may raise one hand or explicitly communicate their need

##### Electrode fitting

EEG, ECG and respiratory belt fitting to be done by the experimenter and research team.

##### Intention Setting

Invitation: Put into words what is alive and present in you in the moment (i.e. it doesn't need to be a whole life story). Take a moment to feel what wants to be put into words. Invitation to specifically share what sensations are alive in the body, rather than 'stories' or 'explanations' of participants' experiences, the invitation is to become more aware of cues from the body.

Invite the participant to share if these sensations in the body are associated with any currents in their life, or processes they are in. While the facilitator will not actively guide the session with a theme, invite the participant to bring this intention with them into their internal practice.

##### Meditative body scan

Invite participant to lie down as if they were preparing for the breathwork session, including using pillows, blankets, or eye masks. Begin the session with a felt sense meditation, guiding the participants' awareness through a 'body scan', becoming aware of each part of their body from their feet to their face. The invitation of this practice is to relax each part of the body and let yourself be carried. Awareness of the breath can be encouraged here, while still breathing a normal breath through the nose. Invite the participant to share one sensation that is particularly noticeable, gathering details about this sensation. If the sensation is tension or negatively valenced, invite the participant to inquire as to where in their body feels the most opposite to this sensation. Inquire about the qualities of this sensation as well, and allow both to co-exist.

**General principles of navigating breathwork experiences (optional):**

- Experience tends to happen in waves
  - If experience comes up, go with it, when you feel like a chapter is coming to an end, return to the breath. These waves can appear as physical impulses, e.g. impulses to stomp the hands or feet, to touch certain parts of the body, to move or shake. They can also appear as emotional impulses such as crying or laughing. Expression of any such impulses is welcome.

- Better out than in
  - If sensations or feelings come up, it is generally recommended to express rather than suppress them. Sometimes suppression of emotions can occur with a desire to stop breathing or an unconscious return to normal breathing patterns. Invitation is to stay with the breath and follow the expressions as they arise.

- There are no distractions
  - Any experience that happens in a breathwork session (e.g. thoughts, body movements, sensations like feeling warm/cold) is very likely to be a potential entrance point into an experiential process. Invitation is to not try to push such sensations away but 'feel into them' i.e. give them awareness and presence.

**Breathwork session (80 minutes)**

- The practice begins with the invitation to move into a circular breathing pattern, with no pauses between inhale and exhale (and vice versa). This will begin through the nose, and at facilitators' discretion, participants can be guided to open the mouth and deepen their CCB, within the first minute or 2 of the session, after the participant is settled into the session.
- Session will be accompanied by evocative music chosen by the research team, with music intensifying towards a peak in the middle of the session, and decreasing in intensity in the second half
- Support by facilitator varies individually for each participant based on the individual experience. Typical needs that facilitators will respond to: Physical help with breathing depth and rhythm; supportive co-regulation during intense emotional experiences (e.g. supportive touch, see *Touch*

*Protocol*); validating feedback when experiencing uncertainty; bringing awareness to tensions in the body, or responding to participants requests for assistance

- For use of bodywork, (see separate *Touch Protocol*)
- Use of bodywork and touch will cease during the 4 minutes of EEG recording, following a prompt from researchers to report the ASC score. This timing will be communicated between researcher and facilitator
- Verbal guidance throughout the session is at the facilitator's discretion and will align with expectations set with the participant, and will include reminders to maintain the pace of the breath, invitations to deepen the breath or encouragement to further explore emotions and sensations as they arise
- After 13 minutes of CCB, participants will be prompted for the subjective experience depth. After researchers have recorded the depth, participants will be instructed to lie still for 4 minutes for EEG recording. No touch or guidance is to be offered during this time.
- After 4 minutes of stillness, participants will be guided to resume CCB for another 13 minutes
- Following the 4th round of CCB and 4 minutes of stillness for recording, an invitation will be made to begin returning to every-day consciousness by self-touch to the arms or heart, small movements such as lifting the knees, or a slow opening of the eyes and returning to the physical space, 'land' (i.e. return to every-day consciousness)
- An invitation to take care of physical needs, facilitators will keep an eye on participants needs for support in returning to the space, either by offering physical touch or a verbal check in

##### Post-session sharing

Post-session sharing is optional. The same protocol for sharing will be used as pre-session sharing.

Invitation to notice what wants to be put into words and witnessed by the group. Encourage participants' to stay with their experience in the moment and resist an urge to 'rationalize' or turn their experience into a 'story' to be understood. Sharing is an invitation to put into words any aspects of the experience that really would like to be witnessed or verbally expressed. This is also an opportunity for participants to provide feedback on the facilitator or their experience of the music and any touch offered.

##### Integration advice

Advice on integrating breathwork experiences into every-day life, mention the process of integration is about returning to wholeness and unifying the experiences and insights from the breathwork session into everyday life

- Invitation to handle oneself gently since perception and emotion are typically more sensitive than usual after a breathwork session, to notice one’s needs and if possible make space for them
- Advice to stay with feeling and sensations, and not land on one definite interpretation too quickly
- Advice to implement insights into practical actions step by step rather than immediately
- Invitation to reach out for support from facilitators if emotional processing should become intense
- Invitation to also drink plenty of water following the session, particularly if there were any intense emotional releases
- Reminder of integration zoom call next morning

**Video Annotation:**

- Following the session, participants will have the choice of conducting their video annotation in the presence of the facilitator and the experimenter, or just the experimenter
- Begin the video at the first

#### Supplemental Text S2: Protocol for Facilitator Touch Interventions

##### Background and Purpose

The purpose of this protocol is to provide clear guidelines for the use of touch by facilitators during breathwork sessions within the context of the RESET Study conducted by Ernst Strungmann Institute for Neuroscience. While touch has long been an element of breathwork for many practitioners, who have anecdotally reported it to be experienced as helpful, there is very little scientific evidence on when and how exactly supportive touch can benefit breathwork processes. A recent survey conducted by the study team with over 350 respondents who had experienced breathwork in different settings suggests that facilitator touch is perceived as helpful in a large majority of cases (Fig. 1, left), and that while most breathwork experiences are evaluated as beneficial in hindsight, sessions that featured touch were seen as slightly more beneficial on average (Fig. 1, right).

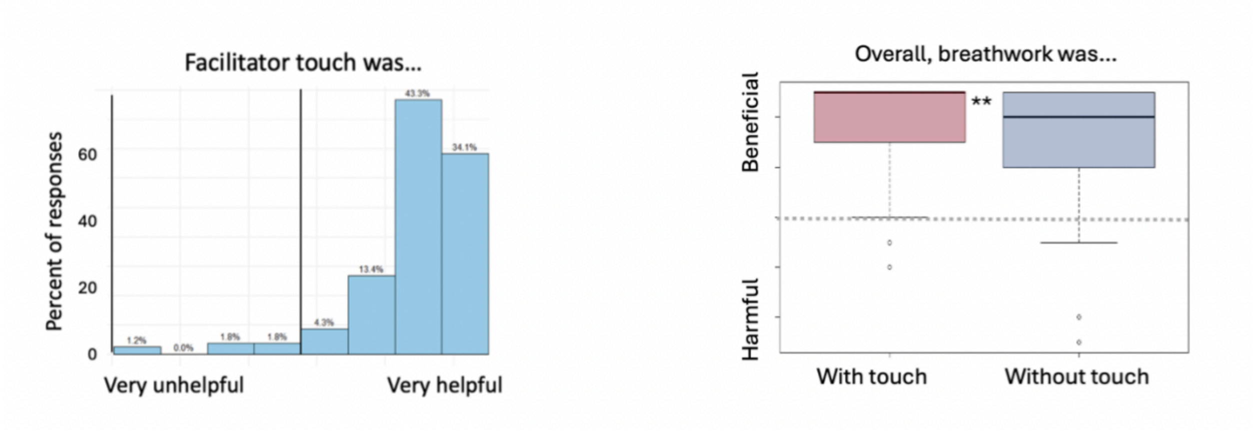

*Figure 1. Experiences of facilitator touch in 366 survey respondents across a spectrum of breathwork settings and experience levels, based on their most recent breathwork experience. Left: Facilitator touch was experienced as helpful by a large majority of respondents, but as unhelpful by approx. 5% of respondents. Right: Overall estimate of whether a breathwork session was experienced as more beneficial or more harmful, split by survey respondents whose sessions had and had not featured touch (approx. 50% of respondents each). While breathwork was rated as predominantly beneficial in both groups, respondents whose sessions had featured touch indicated rated the benefits somewhat higher (Hartter, Badanova, in prep.).*

Based on these insights, the RESET study welcomes careful, respectful and intentional facilitator touch (see below) as an element of its breathwork interventions. Documenting the role of facilitator touch in this study will also serve to strengthen the evidence base for designing future breathwork interventions. Based on the survey introduced above, we can identify two risks regarding the use of facilitator touch:

Too much and too little. As Figure 2 shows below, both incidents appear to be rather rare, with a probability of approx. 5% for experiences that featured clearly too much or too little touch. However, when such incidents occur, they have the potential to lead to aversive, and possibly even damaging, outcomes. Too much touch can evoke experiences of boundary crossing, disempowerment, disrespect and intrusiveness, while too little touch can cause experiences of overwhelm, panic or feeling left alone or abandoned in the face of difficulty. Since this study works with participants that may be particularly sensitive to such potentially retraumatizing experiences (see below), we put particular emphasis on using supportive facilitator touch in a thoughtful, respectful, empathetic, safe and of course fully consensual way.

As such, this protocol aims to ensure that touch interventions are consistent and rooted in the supportive nature of the breathwork process. Given the variability in facilitators' approach to offering supportive touch, the goal is to create a unified and explicitly stated approach to facilitator touch as a variable in the group session, enabling us to better understand its effects while ensuring a safe, healing, and respectful environment for all participants.

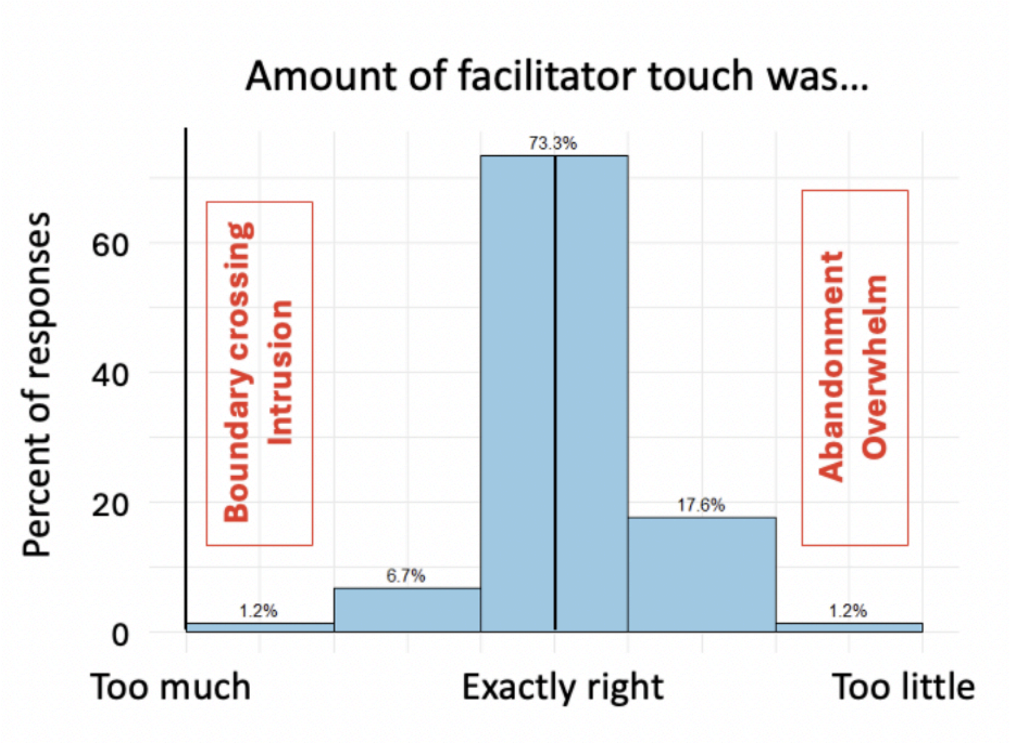

*Figure 2. Experiences of facilitator touch in 366 survey respondents regarding the amount of touch provided. While the majority of respondents (approx. 73%) reported that they experienced facilitator touch as exactly right for their needs, a percentage of respondents also experienced sessions that featured too much or too little touch. Red text boxes highlight potential themes of retraumatization at both ends of the spectrum (Hartter, Badanova, in prep.)*

**General Principles of facilitator touch**

● **Consent**: While supportive facilitator touch is offered as part of the breathwork intervention in this study, participants always have the choice to decline physical touch, and to communicate their preferences before and throughout the session. Facilitators will ask for verbal and non-verbal consent before the session (see below). Participants can change their consent to touch in general, or specific instances of touch, at any time during the session.

● **Verbal and non-verbal cues for consent** will be discussed and practiced between facilitators and participants prior to the session (see below)

● **Intentional Touch**: Facilitators must be clear on their intention before initiating touch. The intention for the use of touch during the group session is to provide emotional support or somatic awareness, not as a means of proactively eliciting arousal or activation ('triggering') in participants.

**Consent for Supportive Touch in the RESET Study**

Before the beginning of the breathwork session, facilitators will explain their intentions for the use of touch during the session and will demonstrate potential uses of supportive touch – especially including areas that participants may find startling or intrusive (e.g. the belly and heart region). Participants will be asked to indicate their basic comfort level with supportive touch via an open conversation.In addition, participants are invited to communicate more specific touch preferences (e.g. specific body areas that are off-limits for them) in one-on-one conversations with facilitators prior to the session and/or by writing them onto their stickers. Facilitators will take these baseline preferences into account when choosing touch-based interventions as set out below.

Participants are also informed that they can change and communicate their preferences towards touch at any time during the session. Facilitators will emphasize their understanding that providing real-time feedback during the breathwork session can feel difficult. To lower the threshold for real-time feedback, before the session participants will practice verbal and non-verbal ways of declining physical contact ('Hands off practice', see breathwork protocol)

If one form of physical touch is communicated to be unwanted during the session, facilitator is welcome to inquire if all touch is unwanted, or if just in specific areas. Verbal consent will be asked for prior to any other forms of touch, if one form of touch is unwanted.

**Guidelines for Intentional Touch**

The intention of touch interventions is either to assist the participant in deepening their breath, to increase somatic awareness of inner (somatic and/or emotional) processes and blockages, or to provide emotional support and grounding during vulnerable and/or intense experiences. As such, touch will be offered when there are signs that a participant…

● is not breathing deeply, has inadvertently deviated strongly from the recommended breathing rhythm or is experiencing other breathing issues

● is experiencing a visible emotional release that appears difficult to process alone

● Is experiencing persistent tensions (signaled e.g. by general stiffness in the body, by cramping of specific body areas, or by communicating other forms of physical or emotional discomfort like heat/cold, anxiety or confusion)

In response to such occurrences, facilitators may apply touch as consented to by the participants in service of the following aims :

- Reconnection and Continuation of Breath

  Example cases and touch interventions:

  - If a participant has stopped their deep circular breath or appears to be dissociating, touch may be used as a tool to help re-establish a connection to their body and breathing pattern.
  - If verbal cues to continue breathing or deepen the breath have not been successful, a gentle touch (e.g., on the hands, chest, or solar plexus) may help draw the participant's attention back to their body and encourage continued breathwork.
  - For example, if a participant is chest-breathing, the facilitator may gently place the participant's hand on the belly, and potentially add pressure with their own hand on top to encourage diaphragmatic breathing.
  - In the case of shallow breath, the facilitator may also apply touch or pressure to the diaphragm for the purpose of providing awareness of full contraction of the diaphragm. Additionally, a lifting of the ribcage may be used to provide the sensation of a full expansion of the lungs

- Emotional Support

  Example cases and touch interventions:

  - Touch can be used to offer emotional support, especially if intense emotions are aroused during the session. Facilitators may provide touch to signal to the participant that they are safe and not alone in their experience.

- Emotional support touch may involve placing hands on the feet, shoulders, chest, or hands, depending on participant's comfort level and facilitator's impression of the process.
- Depending on the nature of the emotional experience, touch may be more gentle (e.g. in vulnerable processes) or somewhat more robustly supportive (e.g. to counteract anxiety or overwhelm) - robust touch should of course always stay within an intensity that feels comfortable and unintrusive to the participant. If in doubt, facilitators should err on the side of applying less and more gentle touch.

- Somatic Awareness:

  Example cases and touch interventions:

  - Touch may be used to help participants become more aware of where they are holding tension in different areas of the body.
  - If a participant is tense or holding their breath, and verbal invitation to relax the chest/belly is not having an effect, a facilitator may carefully touch areas where tension is present, such as the belly, chest or shoulders, to encourage release of the tension. For potentially sensitive areas like the chest, it is also recommended to place the participant's hand on the area and one's own hand on top.
  - For example, if the participants hips and legs are particularly stiff, touch may be applied to the hip bones in a steady or a rocking motion to incorporate a sense of flow and movement back into the body.

Principles and aims of intentional touch are communicated to the participants before the session, and different touch locations and qualities are demonstrated.

**Amount of Touch**

The overall amount of supportive touch that is helpful to a process varies widely across individual breathwork experiences. In many cases, no facilitator touch is needed at all. In others, a few (1-3) touch interventions across the session will be enough to support specific points of the experience (see previous section). In rare cases, especially during emotionally intense or physically very expressive experiences, more sustained contact may be required. If in doubt, facilitators should err on the side of less and gentler touch.

**Appropriate Areas to Offer Touch**

Areas of the body where supportive touch may be offered include:

- Chest: to direct attention to ease/difficulty of breathing into the chest, and to offer emotional support around the heart region (keeping distance from the breast, see below)
- Shoulders: to provide emotional support or signal presence of the facilitator, and to bring awareness to tension that may be held in the shoulders
- Belly, Solar plexus: to increase somatic awareness of belly breathing, to direct attention to continuation of the breath, and to add awareness of tensions
- Hands: To convey presence of facilitator, to provide emotional support, and to give robust support e.g. if participants want to squeeze facilitator's hands during intense processes
- Feet: to convey presence and robust support (grounding)

**Areas that are off-limits for touch**

Areas that should not be touched include all areas commonly associated with sensual touch. This includes genitals, breasts, as well as the areas directly surrounding them (e.g. upper part of the upper thighs, lower belly). Areas that might be interpreted as close to off-limit zones (e.g. the heart region) must be handled with particular care with regards to touch, e.g. by placing the participant's hand on the area and one's own hand on top, and/or by using types of touch that are extremely unlikely to be received as potentially sensual (e.g. pressure with two fingers).

**Intentions for Offering Touch**

Before offering any touch, facilitators should be mindful of the following:

- *Clarity of Intention*

  Facilitators should ask themselves:

  - Why am I offering physical touch at this moment?
  - What specific type of support am I aiming to provide through this touch?

The facilitator's intention should align with the participant's needs, whether that be somatic awareness, breath support, or emotional connection.

- *Space for adjustment and feedback*

Facilitators need to be aware of the fact that their inferences concerning a participant's process and the resulting needs and wishes for touch may be wrong. The aim is therefore to leave as much space as possible to adjust to the participant's needs in real time. To this end, facilitators should keep in mind the following principles:

- If in doubt, provide less and more gentle touch.
- 'Start low and go slow': Increase touch gradually in steps, leaving time to notice verbal or nonverbal feedback from the participant (see below). This can involve starting from gentler touch qualities (e.g. soft touch or placing the participant's hand rather than one's own), and/or from less potentially intrusive touch locations (e.g. hands rather than belly); then if it seems indicated and has been consented to, facilitators can gradually increase the intensity of the touch intervention based on verbal and non-verbal cues (see below).
- As they apply touch, facilitators should be aware of body language and non-verbal cues. This especially includes nonverbal signs that may indicate discomfort, even if the participant has verbally consented to touch, for instance signs of physical withdrawal, startle, increased alertness or tension in response to touch. It can however also include signs that more touch is wished for, e.g. participants beginning to smile when they feel touch, reaching out for the facilitator's hand, indicating body areas that are experiencing discomfort, or making self-soothing motions (e.g. self-hugging) that do not seem to reduce their discomfort.

**Guidance for offering touch around electrodes**

Participants will be connected to an EEG and ECG electrodes as well as a respiration belt during the session. These electrodes and measuring tools are sensitive to movement, therefore physical touch should be offered with consideration for minimally interfering with these tools. No physical touch is to be offered during the 4 minutes of recording following the ASC report. Suggestions regarding touch placement are as follows:

- No touch of the head or neck that may interfere with or misplace the EEG cap
- No touch over the ECG electrode on the chest
- Mindful touch of the shoulders so as not to block the camera or misplace the EEG cap
- Mindful touch of the ribs and belly, assuring the respiration belt stays in place. If misplaced, facilitator will readjust the belt, communicating with the experimenter if signal is lost or regained

**Documentation and Monitoring**

To ensure consistency and improve on the role of touch in this study and in the design of future breathwork interventions, the use of touch will be recorded via a post-session debrief and participant surveys.

- After each session, facilitators will offer participants an opportunity to discuss their experience during the session in a post-session group share.

- Post-session surveys will invite participants to report on their experience of touch interventions and provide feedback on how touch influenced their session
- Before sessions 2-4, facilitators ask participants to share if they want more, less or the same amount of touch as in the previous session.

Facilitators will periodically (after every second session) review their facilitation experiences in the study in general, with particular emphasis on touch interventions, and reflect on how touch influenced the session. This will include evaluating their intentions and the impact of their touch on participants during facilitator intervision sessions.

**Conclusion**

Touch is a valuable tool in the breathwork process, offering somatic awareness, breath support, and emotional connection. However, it is crucial that touch interventions are conducted with clear intention, respect for participant autonomy, and a trauma-informed mindset. By following this protocol, facilitators help to maintain consistency and transparency of their interventions, contribute to the study’s understanding of touch as an important factor in breathwork experiences, and ensure that all participants have a safe, supportive experience throughout the RESET Breathwork Study.

#### Supplemental Text S3: Examples of verbal session annotations

*Excerpts below are drawn from post-session annotations of session video recordings by participants for time points that were rated as ASC intensities 4-5 during the session. Transcripts are lightly edited for readability (filler words and false starts removed; punctuation added; italics added to highlight important themes).*

**Participant 1:**

> "There was something going through my body and it felt then like a release, and afterwards everything was flowing more in the body — it started to feel like a wave in the body. *Energy blocks were released.* So the head is also moving. The wave went really through from my hips, or even my knees, to my hips, through the spine, until the head. I think it was when I had that feeling that I can move freely… I have on land my problems, cause of the accident, the scoliosis and all that stuff… It's there, but I can move… *I could really move freely without any stiffness or handicap*."

> "It's like a big cathedral with — how do you say, is it entities? Like spirits. Yeah, like spirits, a lot of spirits. And I then felt really not only connected, but in between… My dad was there, my mom was there — they're already dead, and a lot of dead people, yes, all around me. [What was that experience like?] It felt safe, known — I am known and seen and heard by them, but I also know that I'm not there, I have their support. Thank you, their support here on earth."

**Participant 2:**

> "A lot of anger coming through, for sure. This was very interesting because it was like a lasagna — a layer of anger, then something softer underneath. It was almost like a level of protection sitting on another emotion. The anger was clearing, and then it was like sadness or softness underneath. And that happened twice. You can see the trembling in the jaw — all this stored anger being released. And in the face, in the solar plexus, and also in the belly, hip space."

> "Gratitude flowing through, for this softness, for this experience. I can still feel a resonance right now in my body — this is how breathing should be."

> "There's this nervousness that I carry, and I know that my mother was very anxious when I was in the womb. And now that we're talking about this, it's almost like the release of this anxiousness — *even generational trauma that's being undone.* I can't even begin to describe it."

**Participant 3:**

> "This *massive energetic opening of my heart*, where suddenly all that space I hadn't felt able to find really opened, and it was specifically when you were [touching] on the back of my ribs, at the back of

the heart — the energy feeling was my heart was just expanding, and it had room. It was quite significant. My mouth — I mean, it's probably such small movements, but I could feel them inside — my mouth and my jaw were really vibrating in a way I haven't really experienced before; I've seen people have it when they're going into [an altered state]. And I remember wiping my eyes as well. At the same time that my mouth and jaw were vibrating, my eyelids were flickering — *there was like a whole reset thing going on*."

**Participant 4:**

"I could feel that tension in the shoulder — so just trying to create some space there, and then the left hand — quite quickly I could almost feel like *energy moving through the body*. It just really helps to move with it rather than resist it. So, dropping into a little bit more of a *surrendered space* of, okay, I'm *not controlling what the movements are,* just allowing what feels right and feels good for the body to move through, which is nice."

"Throughout the whole process, but especially when we're doing physical movements, it's not just *physical tension being released — it carries emotional content and energetic content.* At this point it wasn't crystal clear what I was unwinding, what the memory or the moment was — it just felt almost like a chronic rigidity and tension that isn't just physical but also carries emotional weight. By unwinding it, there was more of a sense of relief and peace that wasn't just physical — actually, *emotionally, I feel lighter.*"

"It was like [my] body knows what we're talking about here and … *another voice entirely outside of myself giving me affirmations of what I needed to hear to let go of what I've been personally carrying*…. "

"[This] big knot was kind of like unwound… now you can move, now you can float… almost like out of body; I don't really know where I was floating or being but it felt like pulling out of my body… absolutely everything was going to completely dissolve. Yeah, definitely felt like coming out of this physical space, *out of this physical body sensation*.

**Participant 5:**

"That was *pure unadulterated joy*… it was just this sense of *absolute joy*, like forget about all of the... the nonsense that you know life kind of like piles on top of you just let it go and just have fun and it was the most *beautiful sense of play* … I knew I was in the room but like I was also kind of like *traversing other dimensions* like I was completely in a different place… there was just *no real sense of time* to be perfectly honest it was just gone."

"The vibration in my body was insane. It felt like I could like shoot lightning bolts out of my fingertips…I felt like a sense of electricity in my body… I remember this sense of just this *powerful* *ball of energy that was literally malleable*, and it was palpable — I could actually play with it, like putty in my hands. It's really a nice feeling."

"This was literally the hand of God — there was a moment, how everything just kind of all came together, and it was almost like at the same time — we were in this whole thing together — and there was just this *explosion of light*, exploding from that mattress, boom, outwards, and it was this sudden *sense of absolute awe and wonder at this incredible miracle of being alive*, being these human beings in this cosmic realm — everything just felt like a *sense of absolute magic*. It was like an explosion of intensity that kind of just knocked me back."

**Participant 6:**

"The feeling of connection and being in the ocean, a little bit like when my mother was pregnant with me, in the womb… yeah like in the placenta, that's the the safest place for every human being and for me also. I don't know exactly why I cry, but it was a kind of, yeah, relief, just let go. It was there inside me and it was the moment to let go and I don't have to hold them back. It felt really relieving, it felt really positive, nearly like *blissful tears*, or was it really like maybe *awe*? Yeah, deep connection and yeah, wholeness. Yeah, wholeness.

"It was really *rebirth of my inner child* which is safe and connected. It's like a tunnel of life coming very deeply to myself. The head is quiet. It's like swimming in the ocean of life, being connected with me and the life… feeling very deeply safe and I don't have to do anything to get worth and just be there, the only thing to do is to be, nothing more and that's a very relaxing state… *Every human being wishes to be there in this state*."

**Participant 7:**

"It started to get like kind of psychedelic already a bit yeah, [the] dimensions of psychedelic was that a visual thing, yeah like more more colors… when I was closing my eyes, sometimes there were some patterns."

"I had like really a *deep relaxation and surrender*…I totally *felt I was dying actually.* It was really intense because I didn't expect it. I was not scared at all… it was really *a dying process* yeah, yeah…I was like just *completely letting go of all this attachment of all these emotions* and all these um stories and memories and everything like to really detangle… in a very loving way and peaceful way, deep peaceful, deep peaceful way of yeah just letting letting go… yeah….I felt very very safe with you both for sure but of course you were like very close and I think it helped me to let this stage happen."

"I had visuals and I was also not really aware about like '*am I in my body or not*,' I mean I knew kind of where I was, but [in the] very back in my head and was much more just floating. I had a similar experience on DMT about some entities but not so visually clear like on DMT because there I could see them very clear, maybe it just remembered me on that trip, it felt more like a remembrance but it was kind of surprising for me like ‘wow okay you're experiencing [that]’... It was really for me a breakthrough actually in my breathwork journeys.”

#### Supplemental Text S4: PMR Intervention Protocol

##### General Information Before the Session

You will have four 13-minute chunks in which I will guide your PMR session. In between these chunks are 4-minute-long periods where we will measure your brain signal. These sequences are separated by a long tone. As the measurement methods are quite sensitive, I will ask you to stay as still as possible. Additionally, I will check if the cap is still properly fitted on your head. If the cap must be adjusted, I will ask you to lift your head slightly and follow the specific movement *(explain and train with participant sitting)*.

In the beginning and the end of the session, we will conduct a baseline measurement with your eyes open and another one with your eyes closed, each for 5 minutes.

During the exercise, I will guide your awareness to the specific muscle. I will then ask you to tense the muscle. Only tense the muscle when I say "now." I ask you to hold the tension for a while and feel into it. I will then let you know when to release the tension and feel into the relaxation. Each muscle will be tensed and released twice.

##### Introduction

Please get ready to relax now. First, make sure you are lying comfortably. Your feet should be in a relaxed position and fall outwards a little. Your body is supported by the mattress. Your arms and hands rest loosely beside your body with your hands turned up. Your head is in a comfortable position.

We will now do two 5-minute measurements. The first one with your eyes open and the other one with your eyes closed. For the first measurement, with your eyes open, please fixate on the camera objective on the ceiling.

##### Start recording and experiment

##### First 13 Minutes

Let's start with feeling your legs on the mattress, your back, and your arms on the side. Mentally scan through your body and notice which muscles are tense and which are already quite relaxed.

If you feel like it, take a deep breath and exhale slowly. Then let the breath flow naturally, noticing how your belly rises as you inhale and falls as you exhale. You might also notice the cool air entering through your nose and warmed by the body as it leaves.

We will begin the exercises now. Pay attention to how your body feels when tensing and then relaxing the muscles. The goal is not to tense the muscles as much as possible, but to clearly feel the difference between tension and relaxation. Continue breathing normally as you tense the muscles. Please only tense the muscles when I say "now."

**PMR start:** Focus your attention on your right hand and forearm. Make a fist with your right hand — now. — Hold the tension for a moment and observe the sensations of tension in your hand and forearm. Continue to breathe normally while holding the tension. — And with the next exhale, release the tension in your hand and forearm, and relax the muscles. Notice the difference between the tension before and the relaxation now. Pay attention to what feeling develops in your hand and forearm — maybe a slight tingling, a feeling of heaviness, or warmth. Follow this feeling with each exhale.

And now, once more, make a fist with your right hand — now. — Again, notice the tension building in your hand and forearm. Hold it for a moment — and with the next exhale, release and relax again. Notice how the relaxation deepens this second time, spreading even more through your hand and forearm.

Move your attention to your right upper arm. Bend your elbow slightly with your hand open and lift your lower arm a little bit from the mattress — now. — Feel the tension in your right upper arm. Hold the tension for a moment — and with the next exhale, lower your arm and relax it. Again, notice the difference between the tension before and the relaxation now. Observe how the relaxation spreads in your right upper arm, in your right forearm, in your right hand, and in each finger.

And now, once more, bend your elbow ever so slightly, keep your hand open and lift your lower arm a little bit from the mattress — now. — Feel the tension again in your right upper arm. Hold it — and with the next exhale, lower your arm and relax more deeply. Notice how the relaxation increases, spreading even further through your right arm.

Let your right arm rest comfortably, and now focus your attention on your left hand. Make a fist with your left hand — now. — Observe the sensations of tension in your hand and forearm, hold it for a moment — and then, with the next exhale, release the tension and relax. Again, notice the difference between the tension before and the relaxation now.

And now, once more, make a fist with your left hand — now. — Feel the tension in your left hand and forearm. Hold it — and with the next exhale, release and relax even more deeply. Notice the relaxation spreading through your hand and forearm.

Now, move your attention to your left upper arm. Lift your left lower arm slightly off the mattress — now. — Feel the tension in your left upper arm, hold it. Continue to breathe normally while holding the tension. And then with the next exhale, lower your arm, relax the muscles, and let go of the tension. Notice how the muscles of your left upper arm, forearm, and hand relax with each exhale.

And now, once more, lift your lower arm slightly — now. — Feel the tension again in your left upper arm, hold it — and with the next exhale, release and relax even more. Notice how this second release deepens the relaxation in your left arm, all the way down to your fingers.

We will now have a measurement of 4 minutes. Please stay as still as possible. I will let you know when we continue.

**Second 13 Minutes**

Let both arms rest comfortably and focus your attention now on your feet. Curl your toes — now. — Hold the tension for a moment and observe the sensations of tension in both feet. — And with the next exhale, release the tension in your feet and relax them. Notice the difference between the tension before and the relaxation now. Follow the feeling of relaxation with each exhale.

And now, once more, curl your toes — now. — Feel the tension again in both feet. Hold it. Breathe normally while holding the tension. — and with the next exhale, release and relax even more deeply. Notice how the relaxation spreads further through your feet.

Now, move your attention to your lower legs. Point your feet — now. — Feel the tension in your calf muscles, hold it for a moment — and with the next exhale, release the tension and relax. Again, notice the difference between the tension before and the relaxation now. Try to relax your calf muscles even more with each exhale.

And now, once more, point your feet — now. — Feel the tension again in your calf muscles. Hold it — and with the next exhale, release and relax even more. Notice how the relaxation deepens in your calves with this second release.

Now, focus your attention on your thighs. Press both heels into the mattress and tense your thigh muscles — now. — Feel the tension in your thighs, hold it for a moment — and then, with the next exhale, release and relax. Notice how the relaxation spreads in your thighs with each exhale.

And now, once more, press both heels into the mattress — now. — Feel the tension again in your thighs, hold it — and with the next exhale, release and let the relaxation deepen. Notice how this second time, the relaxation spreads more fully into your thighs.

Now, move your attention to your buttocks. Squeeze your glutes together — now. — Feel the tension, hold it for a moment — and then, with the next exhale, release the tension and relax. Again, notice the difference between the tension before and the relaxation now. Follow the feeling of relaxation with each exhale.

And now, once more, squeeze your glutes together — now. — Feel the tension again, hold it — remember to breathe normally while holding the tension. And with the next exhale, release and let go even more. Notice how this second release allows the relaxation to spread deeper through your hips and buttocks.

We will now have a measurement of 4 minutes. Please stay as still as possible. I will let you know when we continue.

**Third 13 Minutes**

Now, focus your attention on your face, starting with your forehead. Wrinkle your forehead — now. — Feel the tension in your forehead and scalp, hold it for a moment — and then, with the next exhale, release the tension and smooth out your forehead, making it smooth like a blank surface. Notice the difference between the tension before and the relaxation now. Feel how the relaxation of your forehead spreads to your entire scalp.

And now, once more, wrinkle your forehead — now. — Hold the tension again, and with the next exhale, release and smooth it out even more. Notice how the relaxation deepens this second time, spreading through your forehead and scalp.

Now, focus your attention on your eyes. Squint your eyes slightly and wrinkle your nose — now. — Feel the tension in your entire upper face, hold it for a moment — and then, with the next exhale, release and relax. Again, notice the tension melting away and relaxation spreading.

And now, once more, squint your eyes and wrinkle your nose — now. — Feel the tension again, hold it — and with the next exhale, release and relax even more deeply. Notice how this second release softens your eyes and upper face completely.

Move your attention now to your jaw muscles. Press your teeth and lips together and press your tongue against the roof of your mouth — now. — Feel the tension, hold it — and then, with the next exhale, release and relax your jaw. Notice how, with the release of tension, a feeling of relaxation sets in. Follow this feeling with each exhale, allowing the relaxation to deepen with each breath.

And now, once more, press your teeth and lips together and your tongue against the roof of your mouth — now. — Feel the tension again, hold it — and with the next exhale, release and let your jaw relax even more. Notice how the relaxation spreads this second time through your jaw and cheeks.

Let your face be completely relaxed now, and shift your attention to your neck and throat. Pull your chin to your chest slightly and tense the back of your head by attempting to lift your head — now. — Feel the tension in your neck and the back of your head, hold it for a moment. Remember to breathe normally while holding the tension. — and then, with the next exhale, release the tension and relax the muscles. Again, notice the tension melting away and relaxation spreading. Follow the relaxation with each exhale.

And now, once more, pull your chin in slightly and tense the back of your head — now. — Feel the tension again, hold it — and with the next exhale, release and relax even more deeply. Notice how this second time the relaxation spreads further through your neck and throat.

We will now have a measurement of 4 minutes. Please stay as still as possible. I will let you know when we continue.

**Fourth 13 Minutes**

Now, focus your attention on your shoulders. Squeeze your shoulders back — now. — Feel the tension in your shoulders and upper back, hold it — and then, with the next exhale, release the tension and relax. Let your shoulders drop as low as possible. Let go of all the tension in your shoulders. A pleasant feeling of calm and relaxation spreads out. Follow this feeling and deepen it with each exhale.

And now, once more, squeeze your shoulders back — now. — Feel the tension again, hold it — and with the next exhale, release and relax even more deeply. Notice how this second release allows your shoulders to drop even further, letting go completely.

Now, move your attention to your chest muscles. Take a deep breath and hold it — now. — Feel the tension in your chest muscles, hold it — and then, with the next exhale, release the tension and relax. Notice how, with the release of tension, a feeling of relaxation sets in. Follow this relaxation with each exhale, allowing it to deepen with every breath.

And now, once more, take a deep breath and hold it — now. — Feel the tension again in your chest, hold it — and with the next exhale, release and relax even more. Notice how this second release spreads deeper relaxation through your chest.

Now, focus your attention on your lower back. Make a slight arch in your lower back — now. — Feel the tension in your lower back, hold it for a moment. Remember to breathe normally while holding the tension. — and then, with the next exhale, release and relax. Again, notice the difference between the tension before and the relaxation now. With each exhale, release more tension from your back.

And now, once more, make a slight arch in your lower back — now. — Feel the tension again, hold it — and with the next exhale, release and relax even more deeply. Notice how the second release allows relaxation to spread more fully through your back.

Now, shift your attention to your abdomen. Tighten your stomach muscles as if you were trying to catch a light blow — now. — Feel the tension in your belly, hold it for a moment — and then, with the next exhale, release and relax. Again, notice the difference between the tension before and the relaxation now. Let your abdominal muscles relax and deepen the relaxation with each exhale.

And now, once more, tighten your stomach muscles — now. — Feel the tension again, hold it — and with the next exhale, release and relax even more deeply. Notice how this second release allows your belly to soften completely.

And now, focus only on the pleasant feeling of relaxation. Follow this feeling and try to deepen it with every exhale. Let this peaceful feeling flow into every part of your body: your arms — and hands — and fingers — forehead and scalp — eyes — jaw and cheek muscles — neck and shoulders — down your back — your abdomen — buttocks — thighs — calves — all the way to your feet — to your toes. Let each exhale take you deeper and deeper into relaxation, as deeply as you want and as it feels comfortable. Enjoy this state of peace and relaxation for a little while longer. Take this feeling with you, so you can carry it through the rest of the day.

Now, please tell yourself that you will soon end the exercise. Slowly bring your attention back into this room. Gently tense both hands, bend your arms, stretch, and gently move your body. Take a few deep, energizing breaths — keep your eyes closed.

We will now have two 5-minute measurements again. The first one with your eyes still closed and the other one with your eyes open, looking at the camera. The long tone indicates when to open your eyes. I will also remind you when to open your eyes. For now, please stay as still as possible and keep your eyes closed.
